# Field-isolate recombinant tick-borne encephalitis viruses define reporter-stability guidelines for antiviral testing in flaviviruses

**DOI:** 10.64898/2026.02.23.707037

**Authors:** Benoit Besson, Damien Mouton, Yamina Boukenadel, Theo Nass, Soonju Park, Nakyung Lee, Jianhui Li, David Shum, Redmond Smyth, Stefanie C. Becker, Carine Meignin, Sébastien Pfeffer

**Affiliations:** Université de Strasbourg, Architecture et Réactivité de l’ARN, Institut de Biologie Moléculaire et Cellulaire du CNRS, Strasbourg, France; Université de Strasbourg, M3i UPR9022, Institut de Biologie Moléculaire et Cellulaire du CNRS, France; Screening Platform Laboratory, Institut Pasteur Korea, Seongnam, South Korea; Research Group for Vector-Associated Biodiversity and Infection, University of Veterinary Medicine Hanover, Germany; Research Center for Emerging Infections and Zoonoses, University of Veterinary Medicine Hanover, Germany

**Keywords:** tick-borne encephalitis virus, reverse genetics system, CPER, reporter virus, flavivirus

## Abstract

As arthropod-borne viruses continue to threaten populations globally, there is a pressing need for experimental systems that enable rapid antiviral discovery. Reverse-genetics platforms producing recombinant reporter orthoflaviviruses have been developed to address this gap. Here, we present two new recombinant tick-borne encephalitis viruses (TBEVrec) generated on a European-subtype Haselmühl Tiho1 isolate backbone. A reporter gene, either eGFP or Nluc, was inserted in the capsid-coding region of the genome downstream of the capsid RNA regulatory signal and separated from the complete viral polyprotein by a 2A self-cleaving peptide. TBEVrec was better rescued using the circular polymerase extension reaction (CPER) than with the infectious subgenomic amplicon (ISA) method. TBEVrec replicated efficiently in relevant human cell lines, with comparable replication to wild-type TBEV in a neuronal cell line and moderately reduced titers and RNA levels in immune-derived cell lines. Using either eGFP or Nluc, we illustrate how TBEVrec enabled high-content RNAi screening, highlighting Nucleolin and PRKD1 as potential TBEV host factors, and drug testing on a benchtop plate reader. Nanopore sequencing of the eGFP insert revealed that the reporter is excised without affecting flanking regions. Comparative analysis of eGFP and Nluc further shows that this instability is time-and cell type-dependent, and that Nluc is comparatively more stable. From these observations, we outline safeguards and design principles that are broadly applicable both to the rescue of existing constructs and to the design of future recombinant reporter virus platforms.

**HIGHLIGHTS:**

- GFP and Nluc TBEV reporters built from a field isolate.
- Determination of GFP reporter excision borders.
- Nanoluc reporter shows greater stability than GFP.
- Rescue and early-passaging conditions improve reporter stability.
- NCL and PRKD1 are candidate host factors for TBEV replication.

## 1. INTRODUCTION

In an age when climate change, growing populations, and increased travel redistribute ecosystems across continents, arthropod-borne viruses of the *Orthoflavivirus* genus represent a major global-health challenge (Pierson and Diamond, 2020; Postler et al., 2023). Mosquito-borne flaviviruses account for a substantial burden of disease in tropical and subtropical regions, whereas tick-associated flaviviruses circulate predominantly in temperate and boreal zones. Tick-borne encephalitis virus (TBEV) is the principal tick-borne viral pathogen in Europe and much of northern Asia, causing 10,000–15,000 human cases of encephalitis annually (Chiffi et al., 2023). Recent surveillance data indicate that EU notification rates for tick-borne encephalitis have increased since 2018, with TBEV remaining a persistent public health concern across large parts of Europe and northern Asia (Ackermann-Gäumann et al., 2024). TBEV is genetically diverse and is classically divided into three main phylogenetic subtypes: European (TBEV-Eu), Siberian (TBEV-Sib) and Far Eastern (TBEV-FE), with additional Baikalian and Himalayan lineages proposed more recently (Demina et al., 2010). Together, the expanding distribution and genetic diversity of TBEV highlight the need for constant development of new tools to study its replication and to develop new therapeutics (Bestehorn-Willmann et al., 2023).

The flavivirus genome is a capped, single-stranded, positive-sense ∼11kb RNA (gRNA) containing a single open reading frame (ORF) flanked by 5’ and 3’ untranslated regions (UTRs), but lacking a 3’ polyadenylation (polyA) signal (De Falco et al., 2021). The ORF encodes for a single polyprotein that is cleaved by viral and host proteins into three structural proteins: the capsid (C), pre-membrane (prM) and envelope (E), and at least seven non-structural (NS) proteins: NS1, NS2A, NS2B, NS3, NS4A, NS4B and NS5. The gRNA contains RNA structures in their 5’ and 3’UTRs, as well as other regions such as the hairpin in 5’ of the C gene (cHP), that are essential for the gRNA circularization, replication and translation.

An essential component of the experimental toolbox for flavivirus research is the generation of recombinant infectious viruses expressing reporter proteins, which enable rapid and quantitative readouts of viral replication via fluorescent or bioluminescent signals, such as green fluorescent protein (GFP) or nanoluciferase (Nluc). In that regard, while flaviviruses are particularly difficult to clone due to the instability of their genome in bacterial vectors, significant improvements in reverse genetics have been developed over the years to facilitate the rescue of recombinant flaviviruses (Aubry et al., 2015; Baker and Shi, 2020). Among the most robust strategies for rapidly generating recombinant flaviviruses are bacteria-free rescue systems that assemble circularized PCR amplicons, through variations of either the circular polymerase extension reaction (CPER) or infectious subgenomic amplicons (ISA) methods (Aubry et al., 2014; Edmonds et al., 2013). Most constructs place the reporter in the capsid with a downstream 2A self-cleavage site to allow co-translation of the reporter and viral polyprotein, while ensuring their efficient separation into independent products. Alternatively, insertions can be placed at NS1 or the 3′UTR, with diverse strategies to improve stability (see Table 1) (Baker and Shi, 2020; Shustov et al., 2007).

Recently, several recombinant infectious tick-borne orthoflaviviruses have been generated. For TBEV, these include TBEV-Eu Salland strain produced using the ISA approach (Hoornweg et al., 2023), as well as TBEV-Eu Hypr, TBEV-FE Oshima (low passage) and the Sofjin vaccine strain generated by CPER (Mitsunaga et al., 2025). Fluorescent reporter viruses have also been developed, with mCherry and TurboGFP inserted into the TBEV-Eu Hypr backbone upstream of the capsid, and produced by ISA (Berankova et al., 2025; Haviernik et al., 2021). In addition, eGFP-expressing reporter viruses have been described for other tick-borne flaviviruses, including Langat virus (LGTV) produced via full-length bacterial clone and Powassan virus (POWV) produced by CPER (Conde et al., 2023; Grabowski et al., 2017).

Here, we report two new recombinant TBEV reporter viruses, based on TBEV-Eu field isolate recovered from ticks in a persistent hotspot in southern Germany, and expressing either eGFP (TBEVgfp) or Nluc (TBEVnluc) reporters. We show that CPER is more efficient for rapid rescue of infectious TBEVgfp. TBEVgfp successfully infects relevant human cell lines and constitutes an ideal tool for high-content imaging assays. We further characterized genetic instability of GFP reporter as independent of the design of the recombinant system and suggest alternatives for current and future recombinant flavivirus design. In comparison, TBEVnluc is more stable and allows for rapid drug testing readout on standard benchtop instruments.

## 2. MATERIAL AND METHODS

### 2.1. Cells and virus

Baby hamster kidney BHK-21 cells (ATCC VR-1432), African green monkey kidney VeroE6 cells (ATCC, CRL-1586) and human embryonic kidney HEK293T cells (HEK293T/17; ATCC CRL-11268) were cultured in Dulbecco’s modified Eagle medium with GlutaMAX (DMEM, Gibco) supplemented with 10% fetal bovine serum (FBS, Clontech). Human pulmonary A549 cells expressing bovine viral diarrhoea virus NPro protein (A549/Npro, kindly provided by A. Khol, University of Glasgow, UK) were maintained in DMEM supplemented with 10% FBS and 10 μg/ml of blasticidin (InvivoGen). Human microglial CHME-5 cells (kindly provided by T. Loustau, University of Strasbourg, FR) were cultured in DMEM supplemented with 10% of FBS heat-inactivated for 30 min at 55°C (dFBS). Human monocytic THP-1 cells (ATCC, TIB-202) were maintained in Roswell Park Memorial Institute medium with GlutaMAX (RPMI, Gibco) supplemented with 10% of dFBS. Human neuroblastoma SK-SY5Y cells (ATCC, CRL-2266) were cultured in Ham’s nutrient mixture F12 and DMEM media (1:1, Gibco) supplemented with 15% FBS, 1% non-essential amino acids (Gibco), 2 mM L-glutamine (Gibco), 1 mM sodium pyruvate (Gibco) and 1% penicillin-streptomycin (respectively 100 units/mL and 100 µL/mL). All cell lines were maintained at 37°C in a humidified incubator with 5% CO_2_.

TBEV Haselmühl Tiho1 isolate (European subtype), also referred to as “Haselmühl 303/16” or “BaWa 12-203” (National Consulting Laboratory for TBE, Munich) the virus was passaged four times in mammalian cells: three times in A549 (including the initial isolation) and once in A549/Npro cells (Bestehorn-Willmann et al., 2023; Liebig et al., 2020). Recombinant TBEVgfp and TBEVnluc were produced by reverse genetic. The first passage after rescue (p1) was used unless indicated otherwise. Unless indicated otherwise, infections were performed at a multiplicity of infection (MOI) of 0.1 in serum-free media for 1h on a rocking platform and incubated for 48h in 2%-FBS media. Viral titers were determined by 50% tissue culture infectious dose (TCID_50_) endpoint dilution on A549/Npro cells, using either cytopathic effect (CPE), GFP or Nluc signal, and calculated according to the Reed and Muench method (Reed and Muench, 1938), and the MOI was approximated using 70% of TCID_50_ value. All cell culture work was performed under BSL-3 conditions following institutional safety protocols for TBEV manipulation.

### 2.2. Reverse transcription and PCR

Total RNA was extracted with TRIzol (Invitrogen) and 200-1000 ng (adjusted per experiment) were retrotranscribed with Marathon reverse transcriptase (kindly provided by N. Baumberger, University of Strasbourg, CNRS, FR) as described before (Bohn et al., 2023) using virus-specific primers, see supplementary table S1. PCR amplification of the corresponding amplicons was performed with the PrimeStar GXL PCR kit (Takara Bio) according to the manufacturer’s instructions, see supplementary table S1. Quantitative PCR (qPCR) analysis was performed using iTaq Universal SYBR Green Supermix (Bio-Rad) according to the manufacturer’s instructions on a QuantStudio5 instrument (ThermoFisher). Absolute RNA quantification was obtained by comparison to serial dilutions of pJET-3’UTR, a plasmid containing the TBEV 3′UTR amplicon cloned using the CloneJET PCR Cloning Kit (ThermoFisher).

### 2.3. Reverse genetic system

To generate recombinant infectious TBEV (TBEVrec), a bacteria-free circular DNA system was engineered to express TBEV genomic RNA under CMV, following previously described: CPER or ISA methods (Aubry et al., 2014; Edmonds et al., 2013). A 3’-5’UTR linker and an eGFP or Nluc reporter-2A sequence were synthesized (IDT technology), ligated by PrimeStar GXL PCR and cloned into a pJET plasmid to form the pJET-3’5’-reporter-linker. The 3’-5’ UTR linker includes sequentially TBEV Haselmühl last 83 nucleotides, hepatitis D virus (HDV) ribozyme, simian vacuolating virus 40 (SV40) polyA signal, cytomegalovirus (CMV) enhancer, CMV promoter and at the transcription start position, TBEV Haselmühl first 204 nucleotides. The reporter-2A sequence includes sequentially the reporter gene, porcine Teschovirus-1 2A site, the first 72 nucleotides of TBEV Haselmühl C gene including 27 silent mutations using the Synonymous Mutation Generator (Ong, 2021) and validated for Human/Ixodes compatibility with SnapGene, and the next 79 nucleotides of TBEV Haselmühl C gene (WT genome positions 205-283). SV40 polyA, CMV enhancer/promoter, eGFP and non-coding spacer sequences were derived from the peGFP-C2 plasmid (Promega).

To rescue TBEVrec, we used either the CPER or ISA method, as indicated. The 3’-5’ reporter linker and the genome, divided in 3 overlapping fragments covering the 5’truncated genome (positions 223 and 11098), were amplified by PrimeStar GXL PCR from the pJET-3’5’-reporter-linker or TBEV-infected cell cDNA, respectively, and agarose gel purified with Monarch DNA Gel Extraction Kit (New England Biolabs). Amplicons were combined for circular amplification (10 ng/amplicon, CPER) with PrimeStar GXL PCR using 20 cycles of 12 min extension. The whole PCR reaction (CPER) or purified amplicons (500 ng/amplicon, ISA) were transfected into 70-90% confluent cells with 2 µL of Lipofectamine 2000 in 12-well plates and supernatants were collected at the indicated time. Live GFP expression was monitored at indicated time on benchtop widefield EVOS M5000 microscope (ThermoFischer Scientific).

### 2.4. High-content imaging of siRNA-mediated gene silencing

HEK293T cells were reverse-transfected in 384-well plates with 50 nM of siRNA (Horizon Discovery) targeting the following genes: Non-Targeting Pool (siCTRL), GFP Duplex I (siGFP), Polo-like kinase 1 (siPLK1) or gene-specific ON-TARGETplus SmartPool (Nucleolin (siNCL), Serine/threonine-protein kinase 4 (siSTK4), and Protein Kinase D1 (siPRKD1)). For each well, 0.5 µL of Dharmafect1 transfection reagent (Horizon Discovery) was mixed in Opti-MEM (Gibco) to a total transfection volume of 10 µL and added to 2000 cells in 30 µL of DMEM (WelGene) containing 2% FBS (Gibco) and incubated for 48h. 10 µL of virus diluted in 2% FBS containing media was added to the transfected cells at a MOI of 0.1 and further incubated for another 48h.

After infection, cells were fixed with 2% paraformaldehyde (w/v) at room temperature. Fixed cells were permeabilized and blocked with PBS containing 0.4% Triton X-100 (v/v) and 2% BSA (v/v), stained with mouse anti-flavivirus glycoprotein E antibody (D1-4G2-4-15 (4G2), (ATCC, HB-112) kindly provided by K. Park, Institut Pasteur Korea, KR) at 1:200 dilution for 1h. Following three washes with PBS containing 0.2% Triton X-100 (v/v) and 0.2% BSA (v/v), cells were incubated with goat anti-mouse Alexa Fluor 647-conjugated secondary antibody (Invitrogen) at 1:1,000 dilution and Hoechst 33342 nuclear stain (Invitrogen) at 1:2,000 dilution for 1h at room temperature. Cells were washed three times with PBS before imaging. High-content confocal imaging was performed using an Operetta CLS automated microscope system (Revvity), acquiring Hoechst, eGFP and 4G2/Alexa647 channels using excitation/emission wavelengths of 355-385/430-550 nm, 460-490/500-550 nm, and 615-645/655-760 nm, respectively. For each well, five fields were imaged using a 20× objective, and images were analyzed using Signals Image Artist software (Revvity) and ImageJ.

### 2.5. Virus adaptation and nanopore sequencing

TBEVgfp was passaged four times in A549/Npro cells. RNA extraction, virus-specific reverse transcription and PCR over the reporter region were performed as described above, using 1µg of RNA as input. PCR products were purified from the PCR mixture with the Monarch DNA Gel Extraction Kit, omitting the agarose gel electrophoresis step. Purified DNA amplicon concentrations were measured with the Qubit dsDNA HS Assay Kit (Thermo Fisher Scientific). After normalizing the molar concentration of each sample, a total of 1,600 fmol DNA was used in library preparation, with 5 µL taken from each sample.

For each sample, dA-tailing and 5′ phosphorylation was performed in 6 µL reactions containing 5 µL DNA, 0.7 µL NEBNext End-Repair Buffer (New England Biolabs), and 0.3 µL NEBNext End-Repair Enzyme Mix (New England Biolabs). Reactions were incubated at 20 °C for 10 min, followed by 65 °C for 10 min. Barcoding ligation was performed using the SQK-NBD114-96 kit (Oxford Nanopore Technologies). For each sample, barcoding ligation was carried out in 5 µL reactions containing 1.5 µL end-repaired DNA, 1 µL barcode, and 2.5 µL NEB Blunt/TA Ligase Master Mix (New England Biolabs). Reactions were incubated at room temperature for 20 min, after which ligation was quenched by adding 1 µL EDTA. Subsequently, 5 µL of each barcoded sample was pooled, and the resulting pool was purified with 1× AMPure XP beads and washed twice with Short Fragment Buffer (SFB). The pooled barcoded DNA was eluted in 35 µL nuclease-free water. Adapter and motor protein ligation was performed in 56 µL reactions containing 35 µL pooled barcoded DNA, 11 µL NEBNext Quick Ligation Reaction Buffer (New England Biolabs), 5 µL Native Adapter and 5 µL high-concentration NEB T4 DNA Ligase (New England Biolabs). Reactions were incubated at room temperature for 20 min.

The library was then purified with 0.6× AMPure XP beads and washed twice with SFB, taking care to avoid drying of the beads during washing and prior to elution. The final library was loaded onto an R10.4.1 flow cell (Oxford Nanopore Technologies) and sequenced on a PromethION 2 Solo instrument (Oxford Nanopore Technologies) controlled by MinKNOW software (v25.09.16). Basecalling was performed with Dorado v1.1.1 (Oxford Nanopore Technologies) in super-accuracy (“sup”) mode using the dna_r10.4.1_e8.2_400bps_sup@v5.2.0 model.

Nanopore reads were quality assessed (FastQC, NanoPlot) and filtered (NanoFilt; Phred ≥16, read length 400–1500 nt corresponding to the approximate amplicon sizes). Filtered reads were aligned to the reference TBEV-GFP amplicon using minimap2 in splice-aware mode. Alignment files were processed with Samtools to add headers, convert SAM to BAM, sort, and index the BAM files. Per-base coverage along the amplicon was calculated and normalized to the maximal coverage value for each sample. Junction coordinates were extracted from splice-junction BED tracks exported from IGV. For each sample, junction-supporting reads were summed across strands per junction (start–end) and normalized to the sample’s total junction-overlapping reads; junctions with ≤2 reads were excluded.

### 2.6. Luciferase quantification of compound assay

CHME-5 cells were seeded at 2.5x10^4^ cells per well in a 96-well plate and infected at a MOI of 0.1. At one-hour post-infection, cells were treated with either DMSO (vehicle control) or a 1:3 serial dilution of 7-deaza-2’-C-methyladenosine (7DMA, Merck) and incubated for 48h before collection of supernatants and cell lysis. Nanoluciferase activity was quantified using the Nano-Glo luciferase assay system kit (Promega) and cell viability was assessed by measuring LDH levels in cell lysates with the CytoTox 96® Non-Radioactive Cytotoxicity (Promega). All were acquired on a Varioskan LUX instrument (Thermofisher). Relative luciferase and cell viability were normalized to the DMSO-treated control.

### 2.7. Phylogenetic analysis

The Haselmühl 3′ UTR was initially sequenced by Sanger using six independent bacterial clones. The complete genome of the TBEV Haselmühl strain was then obtained from the viral stock by nanopore sequencing of three overlapping amplicons spanning the full genome, see primers in supplementary table S1. Library preparation, sequencing and sequence analysis followed the TBEVgfp adaptation protocol, except for the read alignment. Reads were aligned with minimap2 (map-ont) to a composite reference comprising the Neudoerfl 5′ UTR, the Haselmühl coding sequence, and the Haselmühl 3′ UTR. A consensus sequence was derived with bcftools.

Phylogenetic trees and pairwise percentage identity were determined as previously described (Besson et al., 2022) with NGPhylogeny.fr, iTol and Mview (Brown et al., 1998; Lemoine et al., 2019; Letunic and Bork, 2021). Pairwise alignments were generated with EMBOSS needle and visualized in Jalview (Rice et al., 2000; Waterhouse et al., 2009). Reference sequences used are listed in supplementary Table S2 and Table S3.

### 2.8. Statistical analysis

Graphical representation and statistical analyses were performed using GraphPad Prism 10 software for non-sequencing, well-level data and in RStudio 2025.9.2.418 for sequencing data and single-cell data. Differences were tested for statistical significance at the well-level using t-test or Kruskal-Wallis, as specified, or for single-cell analysis using a linear mixed-effects model (LMM).^1^

## 3. RESULTS

### 3.1. Rescue of recombinant TBEV Haselmühl Tiho1 reporter virus

For the rapid rescue of a recombinant TBEV-Eu with a reporter gene inserted in the C region (Fig. 1A), we opted for a linker-circularized (Fig. 1B), amplicon-launched strategy (Fig. 1C) such as CPER or ISA (Aubry et al., 2014; Edmonds et al., 2013). This method allows for the recovery of an infectious virus with minimal cloning steps as it only requires the design of a modular synthetic 3’-5’ linker reporter cassette while 97% of the genome is amplified from viral cDNA.

**Figure 1.**
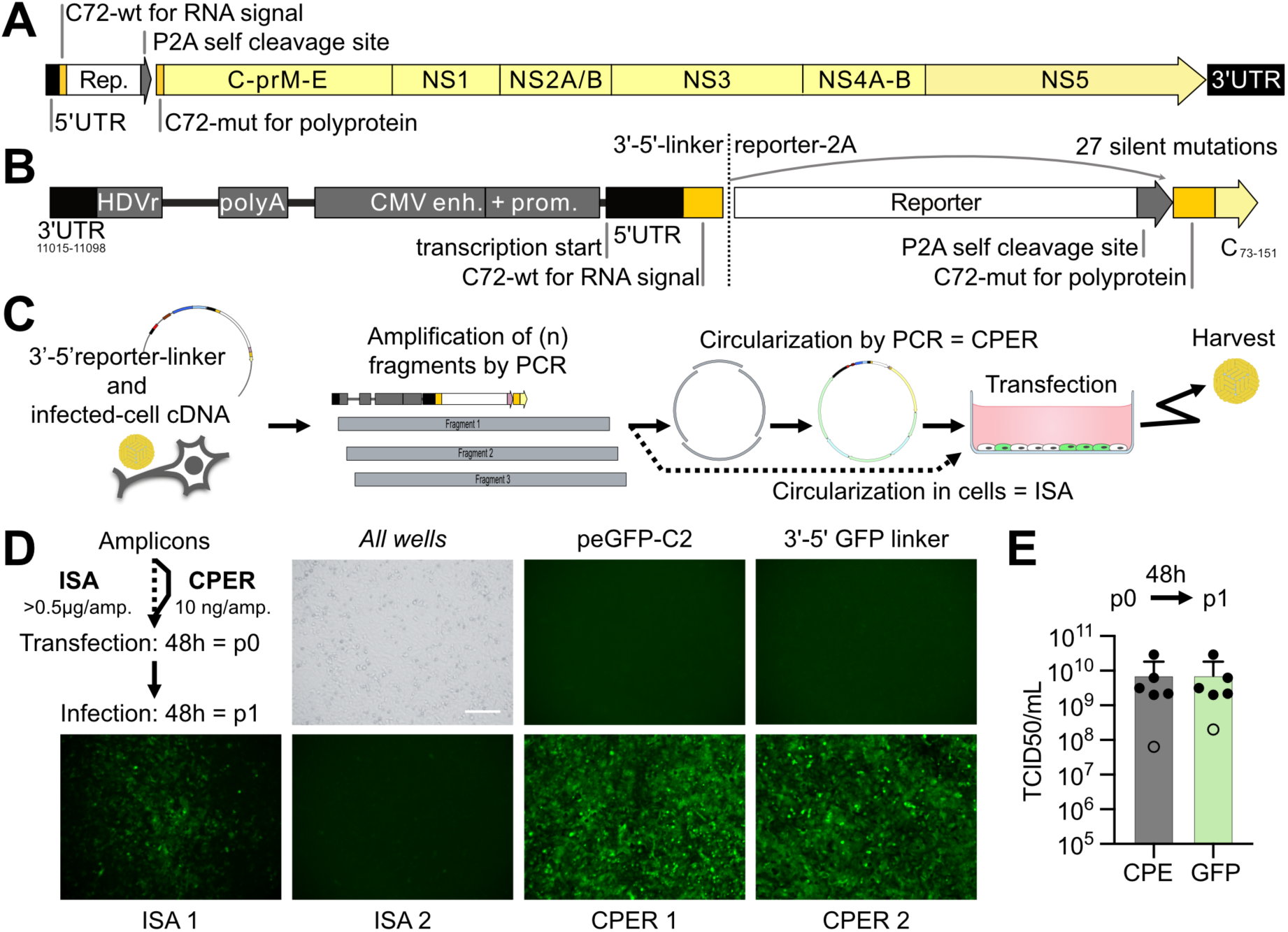
TBEVgfp rescue: design and strategy for recombinant reporter flavivirus systems. **(A)** Genetic map of recombinant infectious TBEV (TBEVrec), built using **(B)** a 3’UTR-5’UTR linker cassette (left) assembled by PCR with the reporter 2A cassette (right) into the 3’-5’reporter-linker cassette, for reporter insertion, DNA circularization and expression in mammalian cells. The viral RNA genome is transcribed in mammalian cells under CMV enhancer and promoter, auto-cleaved by HDV ribozyme and terminated by SV40 polyA signal. The reporter gene is inserted inside a partially duplicated viral capsid gene (C): the 5’UTR and native first 72nt of C (C72-wt) maintain the 5’ RNA viral regulation signal in, the PTV1 2A site (P2A) auto-cleaves the reporter from the polyprotein, and the first 72nt of C is repeated, including 27 silent mutations (C72-mut) to limit recombination and allow the assembly of a full functional polyprotein. **(C)** Rescue strategy: (1) a synthetic 3’-5’ reporter linker is assembled on a plasmid and long viral cDNA are retrotranscribed from TBEV-infected cells, (2) linker and genomic fragments are amplified by PCR to produce amplicons overlapping by 55-83 nt, (3) amplicons are then circularized by PCR (CPER) or directly transfected in cells (ISA) and (4) TBEVrec is harvested from the supernatant. **(D)** Rescue of TBEVgfp on A549/Npro cells. 3’-5’GFP-linker and genomic fragment 1-3 amplicons were either circularized *in vitro* by CPER or directly transfected in cells for ISA. As a control, cells were either transfected with a GFP-expressing plasmid (peGFP-C2) or the 3’-5’GFP-linker amplicon containing the GFP gene under CMV promoter. Supernatants were collected after 48h (p0) and 200µL were used to infect new cells for 48h (p1). Images were captured by widefield microscopy (20x) after 48h of p1 infection. Trans-illumination is shown once and is representative of all samples. GFP images are representative fields for each well; for CPER, two independent fields are shown, and for ISA, one field is centered on a focus and the other is from an area without foci. All transfections were done in duplicate. Scale bar corresponds to 100 μm and is the same for all panels. **(E)** TBEVgfp-p1 produced by CPER was titrated using either CPE or GFP signal as readout, observing one discordant readout (empty dot), n = 6 biological replicates. Cloning map created with SnapGene; virion schematic adapted from ViralZone; transfected well icon from Bioicons.

The whole viral genome is derived from TBEV-Eu Haselmühl Tiho1 (Haselmühl), a low-passage field isolate from ticks collected in 2016 at a TBEV hotspot in southern Germany (Bestehorn-Willmann et al., 2023; Borde et al., 2021; Liebig et al., 2021, 2020; Weidmann et al., 2013). Haselmühl was selected as a contemporary backbone close to Neudoerfl (Fig. S1A), TBEV-Eu reference strain. Sequence analysis of the viral stock is consistent with the CDS consensus sequence of the initial isolate at passage (Liebig et al., 2020). The complete Haselmühl genome was sequenced using Oxford Nanopore Technology and has been deposited in GenBank under accession number PZ023949. Haselmühl shares 97.8% identity over the CDS with closely related strains and Neudoerfl (Fig. S1B). Relying on their close identity, we used Neudoerfl-based primers to sequence Haselmühl UTRs revealing that Haselmühl shares 99.2% identity in the 5’UTR (Fig. S1C) and 91.78% identity in the 3’UTR with Neudoerfl (Fig. S1D). The decreased homology over the 3’UTR is mostly due to the absence of Neudoerfl internal “polyA sequence” in Haselmühl 3’UTR (Fig. S1E) (Kutschera and Wolfinger, 2022). The sequences of the recombinant TBEV (TBEVrec) constructs are representative of the Haselmühl field isolate and are available in Table S3.

We next evaluated Haselmülh growth in human cell lines relevant to pathogenesis: SH-SY5Y (neuronal), CHME-5 (also referenced as HCME-3; microglial) and THP-1 (monocytic) (Fig. S2). SH-SY5Y and CHME-5 were most permissive to viral replication, while THP-1 cells required higher MOI (≥ 1) to reach similar replication levels.

The 3’-5’ reporter linker cassette is assembled by PCR from two synthetic fragments: the 3’-5’linker cassette and the reporter-2A cassette (Fig. 1B). In the 3’-5’ linker, the CMV enhancer and promoter drives the transcription of the recombinant viral RNA, which is self-cleaved at the 3’ end by a HDV ribozyme, and terminated by a SV40 polyA signal. The reporter is inserted after the first “wild-type” 72 nt of the C coding sequence (C72-wt) (Haviernik et al., 2021), which contains the cHP RNA signal (Fig. S1E) required for viral replication, and immediately followed by P2A, the 2A self-cleaving peptide with the highest reported cleavage efficiency (Kim et al., 2011). This configuration results in expression of a reporter flanked by 24 residues from the capsid, and 21 residues from the 2A peptide. The first 72nt of the capsid is then repeated, including 27 synonymous mutations (C72-mut) to minimize the risk of recombination.

TBEVrec carrying an eGFP reporter (TBEVgfp) was rescued by CPER or ISA using the same 3′-5′ reporter linker cassette and viral fragment 1–3 amplicons (Fig. 1C-D). After the first passage (p1), CPER-rescued TBEVgfp produced a fully infected monolayer by 48 hours post-infection (hpi), whereas ISA-rescued TBEVgfp yielded only isolated foci (Fig. 1D). In the negative controls (peGFP-C2 plasmid or the 3′5′-GFP-linker amplicon), GFP expression was transient and GFP-positive cells were detected only after the initial transfection (Fig. S3A) but not detected in the subsequent infection (Fig. 1D). These results are consistent with previous reports of higher SARS-CoV-2 rescue efficiency with CPER compared with ISA (Park et al., 2025). TBEVgfp virus production reached ∼10^9^ TCID_50_/mL on average, whether quantified by cytopathic effect (CPE) or GFP readout (Fig. 1E). TBEVgfp replicated to equivalent titers across IFN-deficient cell lines, although complete monolayer destruction occurred by 3 days post-infection (dpi) in A549/Npro cells and by 4 dpi in BHK-21 cells, while VeroE6 cells showed minimal CPE through 5 dpi (Fig. S3B). Thus, CPER enabled fast and efficient TBEVgfp rescue compared to ISA, producing a viral stock with high titers under 5 days.

### 3.2. Characterization of TBEVgfp as a tool for high-content imaging

To validate TBEVrec as a model for TBEV infection, we compared wild-type TBEV (TBEVwt) and TBEVgfp in SH-SY5Y, CHME-5, THP-1 and HEK293T (Fig. 2A-B). TBEVgfp infected all cell lines but replicated slightly less efficiently than TBEVwt as indicated by lower levels of virus production at 48 hpi. In myeloid-lineage derived CHME-5 and THP-1 cells, TBEVgfp titers and RNA levels were ∼1 log lower than TBEVwt, whereas levels were comparable in SH-SY5Y and HEK293T, suggesting a modest fitness cost that is consistent with the added genetic load of the reporter gene (Berankova et al., 2025; Haviernik et al., 2021).

**Figure 2.**
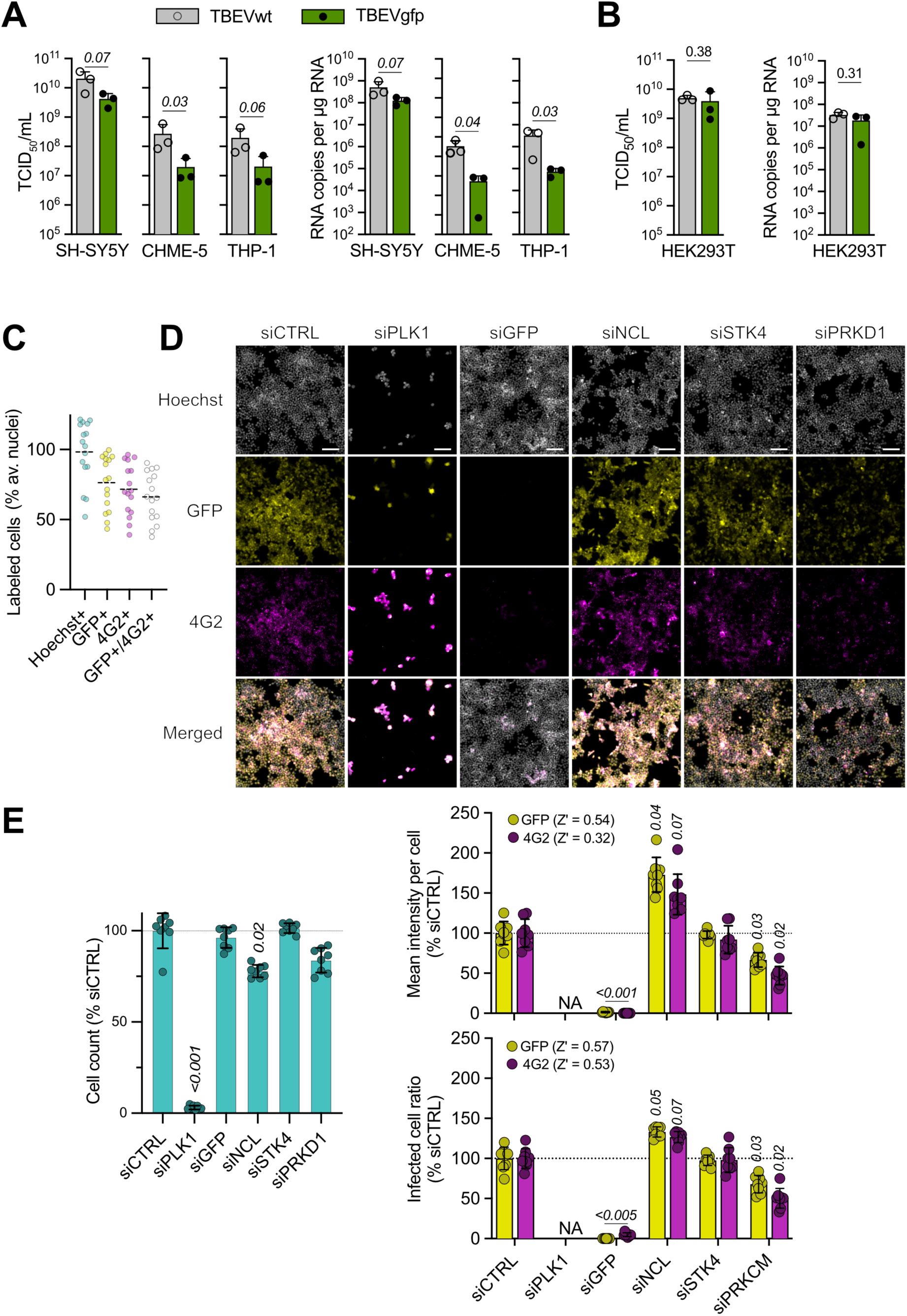
TBEVgfp characterization and application for high-content imaging screening. **(A-B)** Cells were infected for 48h with either TBEVwt or TBEVgfp at a MOI of 0.1 (or MOI 1 for THP-1 cells), and viral replication was quantified by either titration (left panel) or qPCR (right panel). Bar plots represent the means and standard deviation from 3 biological replicates. t-test p-values are shown in italics. **(C-E)** HEK293T cells were transfected with the indicated siRNA for 48h, then infected with TBEVgfp at MOI 0.1 and fixed at 48hpi with paraformaldehyde and labeled for imaging of Hoechst, GFP and 4G2/Alexa647. Images were acquired on a high-throughput compatible confocal microscope (n = 5 images per well). **(C)** Percentage of cells per well (Hoechst), and percentage of GFP and 4G2 positive-cells per well, normalized per well. N = 16 wells. **(D)** Representative images of each siRNA-transfected condition. N = 8 wells. Scale bar corresponds to 100 μm and is the same for all panels. **(E)** Image analysis based on Hoechst, GFP and 4G2 signals normalized as a percentage of the siCTRL. Bar plots represent the means and standard deviation from 8 wells. Kruskal–Wallis p-values are shown in italics.

To establish TBEVgfp for high-content screening, we used an automated confocal microscope to quantify RNAi-mediated effects on viral infection. In non-transfected wells, 71% of HEK293T cells infected at MOI 0.1 were positive for the anti-Orthoflavivirus antigen E (4G2 antibody), 76% of cells were GFP-positive and 66% of cells were positive for both, possibly due to the weaker 4G2 labelling and potential loss of GFP (Fig. 2C). In siCTRL wells, all 4G2/E-positive cells were also GFP-positive (Fig. 2D). Efficient siRNA delivery was verified by 97% cell death with cytotoxic siPLK1 (Fig. 2E, left panel). siGFP targeting the GFP-encoding viral genome abolished the GFP readout (0% GFP-positive), effectively extinguishing detectable replication of the GFP-tagged genome. A small fraction (∼5%) of cells remained 4G2/E-positive, suggesting the selection of GFP-negative TBEVrec variants under siGFP pressure (Fig 2D-E).

To investigate the contribution of host factors to TBEV replication, we silenced key cellular genes previously associated with flavivirus infection using siRNA. We targeted nucleolin (NCL), implicated in *Orthoflavivirus* infections and known to interact with dengue virus capsid and Japanese encephalitis virus NS5 protein (Balinsky et al., 2013; Deb et al., 2024; Naveed et al., 2025), STK4/MST1, a core kinase in the Hippo pathway that is critical in viral infections such as *Orthoflavivirus* Zika virus (Garcia et al., 2020; Wang et al., 2020), and PRKD1/PRKCM, involved in *Flaviviridae* hepatitis C virus vesicle trafficking and reported to increase in activity during TBEV infection (Coller et al., 2012; Sui et al., 2024). GFP and 4G2/E signal quantification indicated that *NCL* knockdown had a proviral effect while *PRKD1* knockdown had an antiviral effect (Fig. 2D-E). Consistent with the well-level analysis, single-cell mixed-effects modelling increased sensitivity by leveraging cell-level intensity distributions while accounting for clustering within wells (Fig. S4D). *STK4*-targeting siRNA treatment had no measurable effect on TBEVgfp replication, although efficient STK4 depletion was not independently confirmed.

Together, these results show that TBEVgfp is suitable for high-content imaging and can reveal host factors influencing TBEV replication, pending further validation. GFP readout correlates with 4G2-Ab/Alexa staining (Pearson r = 0.85-0.87, Fig. S4A) but signal intensity is 7.6-fold stronger with higher signal-to-background ratio (Fig. S4B), and avoids staining steps that can cause cell loss (Fig. S4C), underscoring the practical advantages of TBEVgfp.

### 3.3. Genetic instability of *Aequorea victoria* GFP variant in recombinant viruses

To assess the genetic stability of TBEVgfp, we serially passaged TBEVgfp in IFN-deficient A549/Npro cells for 48 h per passage. When titrating on A549/Npro or Vero E6 cells, discrepancies between CPE and GFP readouts were evident as early as passage 2 (Fig. 3A), consistent with the progressive loss of GFP signal from p1 to p4 (Fig. 3B). PCR amplification of the region spanning the GFP insert showed a gradual loss of the expected 1138-nt product and the appearance of a shorter ∼550-nt product (Fig. 3C). A minor GFP-less population was detectable at p1, and the 1138-nt product was no longer observed by p4, in line with prior reports. Loss of the reporter insert is common in recombinant flaviviruses and is well documented (Dong et al., 2023; Haviernik et al., 2021).

**Figure 3.**
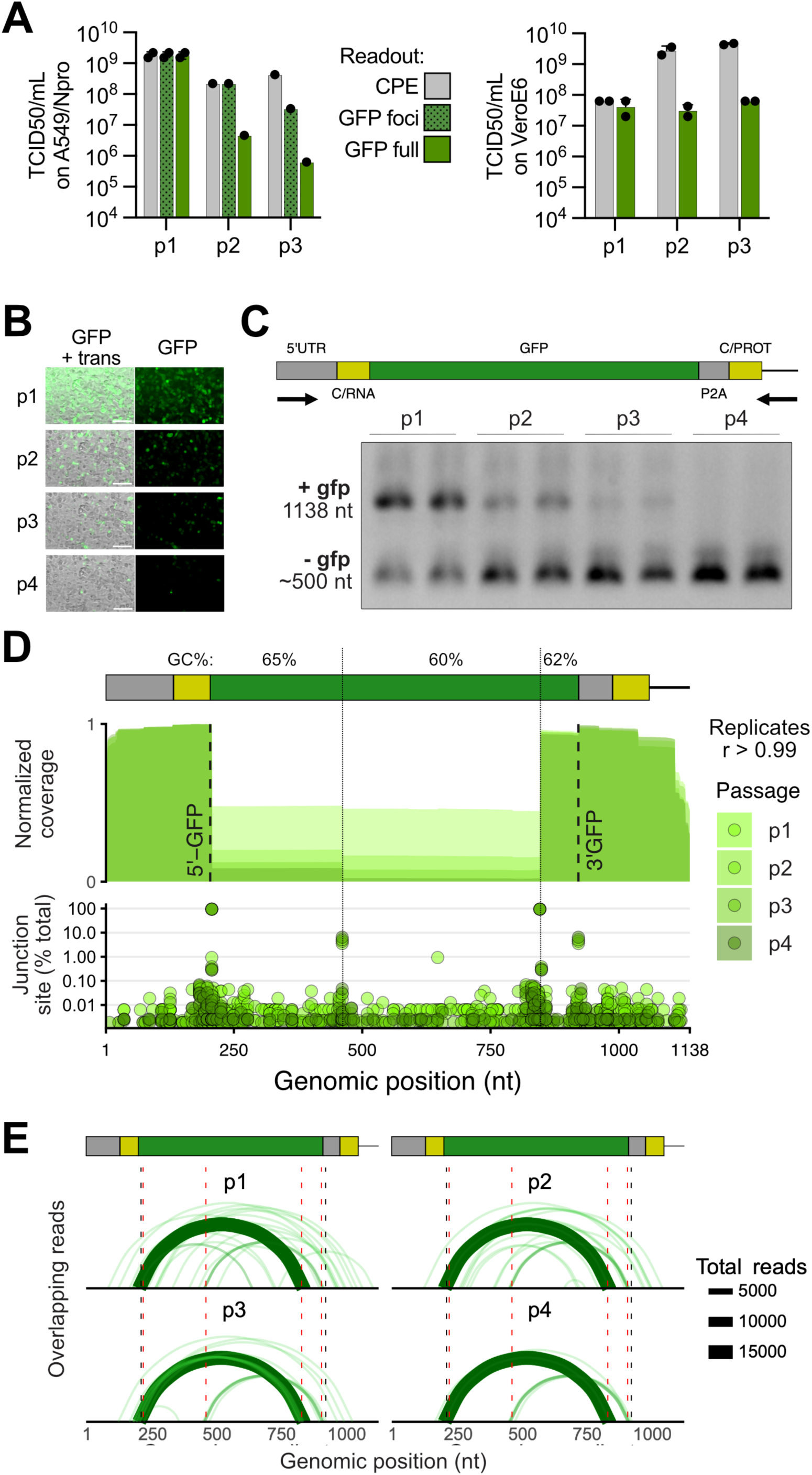
TBEVgfp genomic instability over serial passages. TBEVgfp was passaged 4 times (p1-4) on A549/Npro cells infected at MOI 1 (p1) and then MOI 0.01 (p2-4) and harvested at 48hpi (n = 2 biological replicates). **(A)** Supernatant were titrated on either on A549/Npro (left panel, n = 1-2) or VeroE6 cells (right panel, n = 2), using either CPE or GFP signal as readout. For GFP-positive wells, titers were determined using all wells with at least one GFP foci, or counting only wells with a full GFP monolayer (GFP full), ignoring wells with large CPE but isolated GFP foci. **(B)** Widefield microscopy imaging (20x) of A549/Npro cells at 48hpi. **(C)** PCR amplification over GFP sequence (1-1138 nt) in cell lysate cDNA. **(D)** Sequence coverage normalized to the maximum coverage (top panel) and relative abundance of each junction sites (bottom panel) over the GFP amplicon. Biological replicates were merged for each sample. Dashed line: boundaries of the GFP-coding sequence; thin line: boundaries of the sequence used to calculate GC%; r: Pearson’s r correlation. **(E)** Sashimi-like arcs over the GFP reporter amplicon; thickness is relative to supporting reads per junction; junctions with ≤2 reads were omitted and the top three per sample are highlighted. Major junctions are indicated with red dotted lines at positions 207, 461, ∼849 and ∼921.

To isolate the recombination points and identify weak points for future designs, PCR amplicons from each passage were then sequenced by nanopore sequencing. Nanopore data indicated a higher fraction of GFP-less reads than suggested by gel visualization, suggesting a sequencing bias toward shorter molecules. Coverage profiles revealed that most recombination junctions cluster within the GFP coding sequence (Fig. 3D). These junctions map preferentially to the GFP ORF, defining distinct sub-regions with multiple recombinant species at p1, but only 2–3 dominant recombinants by p4 (Fig. 3E). The most frequent recombination occurred between positions ∼207 and ∼846-849 (93-95% of junctions) on the recombinant genome, followed by a recombination between positions ∼461 and ∼921 (3-6 % of junctions). The third most frequent recombination was ∼207-647 at p1 and then ∼207-849 from p2 onward (<1 % junctions). Recombination starting or ending outside of the GFP coding region were substantially less frequent than recombination within GFP: 0.6% of total overlapping reads at p1, 0.3% at p2-3 and 0.2% at p4 (Fig. 3E). Within the GFP, the segment that is removed by the most abundant mutant with 93-95% of recorded junctions (207-846/9) has 65% GC while the rest of the GFP sequence contains 60-62% GC (Fig. 3D). Together, these observations indicate that the excision of the GFP occurs largely independently of the partially homologous flanking repeats, and may be facilitated by the presence of recombination-prone sites within the GFP coding region that enable partial excision of the insert.

### 3.4. TBEVnluc as stable reporter virus for rapid benchtop drug discovery

As an alternative to TBEVgfp, we generated a TBEVrec expressing a Nanoluciferase (Nluc) reporter (TBEVnluc). TBEVnluc was rescued by CPER, with TBEVgfp used as a control to monitor the appearance of infection foci, which correlated with a strong Nluc signal in cell lysates (Fig. 4A). TBEVnluc was then amplified, yielding ∼10^9^ TCID_50_/mL on average using either CPE or Nluc activity as the readout, comparable to the titers obtained for TBEVgfp (Fig. 4B).

**Figure 4.**
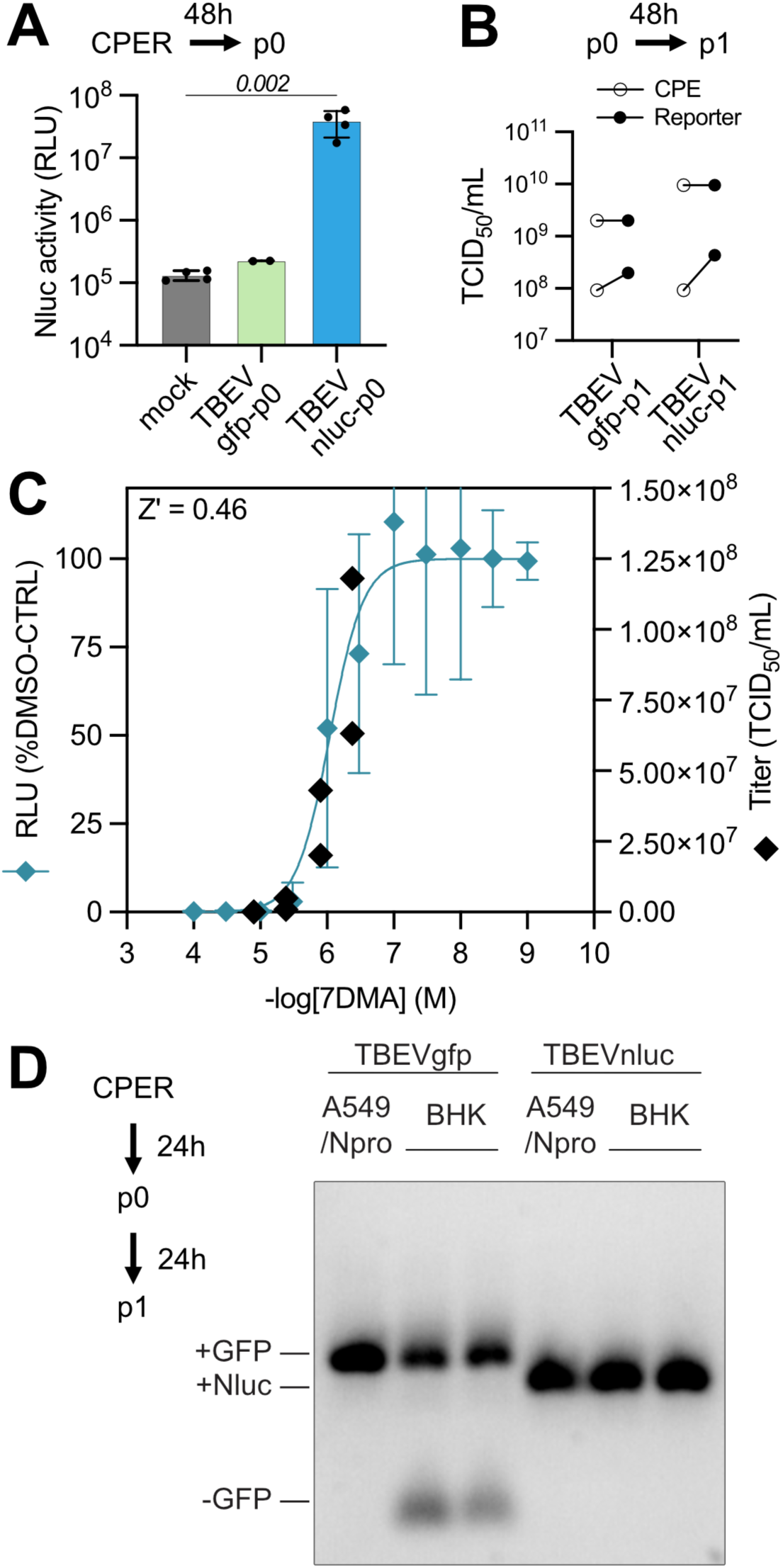
TBEVnluc: characterization of a more stable reporter for benchtop readout. **(A)** TBEVnluc (n = 4) was produced on CPER-transfected A549/Npro cells, using TBEVgfp (n = 2) and untransfected mock (n =4) as control. p0 supernatants were harvested after 48h and Nluc activity was measured in cell lysates. N = technical replicates. Kruskal–Wallis p-value is shown in italics. **(B)** TBEVnluc-p0 or TBEVgfp-p0 were amplified on A549/Npro to produce p1, titrated using both CPE and either GFP or Nluc reporter activity as readout. **(C)** CHME-5 cells were infected with TBEVnluc at a MOI of 0.1 and treated at 2hpi with serial dilutions of 7DMA. Luminescence was assessed at 48hpi on a benchtop plate reader (Varioskan, small dots and non-linear regression), and four 7DMA dilutions were selected for titration in duplicates (large dots). The indicated Z′ factor is the mean of plate-level values calculated from DMSO and maximum-inhibition controls. N = 8 biological replicates **(D)** TBEVgfp and TBEVnluc were rescued and collected 24h after CPER transfection (p0), and then amplified from MOI 0.01 for 24h after infection (p1) in the indicated cell lines. Amplicons covering the reporter gene was amplified by PCR from p1 cDNA. Expected size: 1138 nt (GFP) and 934 (Nluc)

7DMA, a 2′-C-methylated nucleoside with broad antiviral effect and potent anti-TBEV efficacy (Eyer et al., 2023, 2015), was used to validate the TBEVnluc reporter. To this end, we quantified the effect of 7DMA on the reporter virus replication by measuring Nluc activity in CHME-5 microglial cells. Dose–response analysis showed robust inhibition (pIC50 = 6.042), accompanied by a concordant drop in infectious titers after 48 hpi without detectable cytotoxicity (Fig. 4C; Fig. S5). Although we did not directly compare the antiviral potency of 7DMA on TBEVwt and TBEVnluc, these results indicate that the latter is suitable for antiviral testing and rapid readout on benchtop plate readers.

Finally, we compared the genetic stability of TBEVgfp and TBEVnluc in two cell lines (Fig. 4D). To reduce the likelihood of recombination, p0 rescue was restricted to 24 h and p1 was then generated at a low MOI (0.01) and amplified for 24 h. Under these conditions, no GFP-negative revertants were detected in virus stocks rescued and produced in A549/Npro cells, whereas such variants were already present in stocks generated in BHK-21 cells. Moreover, TBEVnluc did not yield Nluc-negative revertants in either cell line. These findings highlight that MOI, production timing, and producer cell line critically can shape recombinant virus stability, and that Nluc is more stably maintained than GFP within the TBEV genome, at least within this experimental timeframe, consistent with differences in reporter sequence composition (Table S4).

## 4. DISCUSSION

Tick-borne viruses cycle between ticks and diverse vertebrate hosts, from small rodents to larger mammals and birds. As such, they occupy specific ecological niches and occur in small, geographically separated foci where viral lineages evolve independently, promoting adaptation and co-evolution between TBEV and its local tick vectors and rodent reservoirs (Kauer et al., 2024; Liebig et al., 2021). This diversity contributes to variation in the effectiveness of existing vaccines against circulating strains (Bestehorn-Willmann et al., 2023). Moreover, no specific antiviral drugs are approved for TBEV, underscoring the need for tractable assay systems to study their replication and assess therapeutic solutions. Ruzek and colleagues (Berankova et al., 2025; Haviernik et al., 2021) recently generated mCherry and TurboGFP reporter viruses based on the Hypr strain, which is to be more closely related to K23, the strain used in the Encepur® vaccine (Bavarian Nordic GmbH). Here, we report two new TBEV-Eu reporter viruses, including the first TBEVnluc virus, based on the circulating Haselmühl strain, which is closely related to Neurdoerfl, the strain used for FSME-IMMUN® vaccine (also known as TICOVAC®, Pfizer, Vienna, Austria). By combining a contemporary wild TBEV-Eu isolate with direct CPER/ISA benchmarking and sequencing-based analysis of reporter-negative variants, this study extends existing reporter-virus work from construct generation to the optimization of rescue conditions and mechanistic investigation of reporter instability.

Multiple strategies have been used to generate recombinant viruses, ranging from bacteria-or yeast-propagated full-length infectious clones to amplicon-launched methods such as CPER and ISA (Aubry et al., 2015; Mitsunaga et al., 2025). Full-length clones facilitate stable genome maintenance but have notable drawbacks: large flavivirus constructs are prone to interference from cryptic bacterial promoters and difficult to clone. By contrast, amplicon-launched approaches are less labor-intensive and modular. PCR fragments can be combined to rapidly assemble recombinant viruses, potentially preserving sequence variants and avoiding sequence recoding required to stabilize clones. However, preservation of sequence diversity can be disadvantageous when a genetically defined virus is required, as input heterogeneity and PCR-derived mutations may generate non-clonal populations and complicate genotype–phenotype attribution, necessitating sequence validation of recovered viruses (Aubry et al., 2015; Dong et al., 2023). When comparing the two methods, CPER recovered recombinant virus more efficiently than ISA in the present study, aligning with reports where SARS-CoV-2 head-to-head tests indicated a length-related benefit for CPER, in contrast to dengue virus, for which separate studies showed similar success rates (Park et al., 2023, 2025; Tamura et al., 2022). Here, we perform such a comparison and reach the same conclusion: CPER outperforms ISA across genome lengths and across the ranges of amplicon sizes and numbers tested, an advantage that becomes critical when genetic instability is a concern.

Reporter orthoflaviviruses are widely used, however many studies report rapid loss of fluorescent inserts during passaging (Table 1) (Baker and Shi, 2020). The GFP from *Aequorea victoria* remains the predominant reporter in molecular biology. Consistent with this, orthoflaviviruses reporter systems frequently use GFP or its derivatives eGFP (Table S1), but use of eYFP, OxGFP, Clover2, mAmetrine, and split-GFP have also been reported (Cherkashchenko et al., 2023; Conde et al., 2023; Suphatrakul et al., 2018; Yi et al., 2011). Fluorescent proteins developed from non-*A. victoria* sources (mCherry, bfloGFP and others, Fig. S6) are not necessarily more stable in these configurations (Cherkashchenko et al., 2023; Haviernik et al., 2021; Mutso et al., 2017; Suphatrakul et al., 2018), although mCherry has been reported to be stable until p3 (Li et al., 2020). Despite this ubiquity, stability depends on genomic context, design and rescue parameters, as illustrated by the data presented here.

Our design follows the common insertion at the capsid N-terminus, separated by a 2A site and distinguished by a codon-scrambled, duplicated capsid N-terminus. Recent studies introduced protective configurations such as an upstream 2A peptide, which has been punctually supported by viability-penalizing safeguard (frameshift or lethal mutation upstream of the first 2A site) to force recombination events to produce non-viable genomes (Baker et al., 2020; Suphatrakul et al., 2018; Volkova et al., 2020). In addition, the duplication of the C72wt upstream of the reporter was also shown to reduce viable deletants although the mechanism remains unresolved (Berankova et al., 2025). In our system, Nluc remained stable through the first passage, whereas GFP-negative variants emerged as early as passage one, with sequencing indicating that excision events mapped within the GFP coding sequence. Collectively, the data implicate recombination-prone motifs within the GFP ORF that enable partial excision of the reporter. In contrast, variants directly joining the flanking regions were extremely rare (<1% events) and their contribution to reporter excision remains unresolved, although flanking-region engineering can improve reporter stability and suppress recombinant outgrowth.

Our observations explain the trends observed in the literature (Table 1). Progressive loss of GFP and variants over passages p1–p4 has been reported for capsid-duplication designs in dengue virus type-2 (DENV2), Zika virus (ZIKV), Kunjin virus, and Culex flavivirus across mammalian and mosquito cells (Cherkashchenko et al., 2023; Dong et al., 2023; Gadea et al., 2016; Park et al., 2023; Schoggins et al., 2012; Suphatrakul et al., 2018). Recent protective configurations (dual 2A linkers, frameshift, duplicated C72wt) delay the emergence of GFP excised variants (Berankova et al., 2025; Suphatrakul et al., 2018). Moreover, even when eGFP persisted to p10, background GFP excision was evident, potentially confined to defective genomes (Volkova et al., 2020). Outliers are a Japanese encephalitis virus (JEV) where eGFP remained stable for 10 passages and a ZIKV construct in which eGFP remained stable for 4 passages (Li et al., 2022; Mutso et al., 2017), while other reports offered limited or no genetic-stability data (Gao et al., 2021; Yi et al., 2011).

Practically, stability at passage one is sufficient for many applications, including drug or RNAi screening as well as for dissecting mechanisms underlying viral replication. We therefore used each reporter according to assay requirements: TBEVgfp for siRNA experiments, where high-content imaging quantified infection together with cell number and image-based quality-control parameters, and TBEVnluc for antiviral dose-response testing, where a rapid homogeneous plate-reader readout provided robust sensitivity. The comparable production yields of TBEVgfp and TBEVnluc in A549/Npro cells, together with CHME-5 titers falling within the same broad range as TBEVwt/TBEVgfp, further support similar replication capacities across these viruses, although the separate assay formats preclude direct quantitative comparison. Using eGFP fluorescence or immunofluorescence of the envelope protein provided the same result and allowed us to identify Nucleolin and PRKD1 as respectively an antiviral and a proviral factor for TBEV. Of note, the antiviral and RNAi experiments performed in this study were intended as proof-of-principle applications rather than comprehensive validation.

Bioluminescent reporters tend to be more durable but are not immune to sequence ejection. Nluc or Rluc were progressively excised from capsid-duplication designs in yellow fever virus (YFV) and JEV between p2 and p5, whereas viability-penalization strategies supported stability up to p10. Even then, recombination between reporter and capsid has been observed (Baker and Shi, 2020; Dong et al., 2021; Li et al., 2017; Volkova et al., 2020). E–NS1 junction insertions are often stable (Bonaldo et al., 2007; Kassar et al., 2017), although results are not universal (Volkova et al., 2020), and stability at the NS1 C-terminus remains sparsely documented (Conde et al., 2023). By contrast, 3′UTR insertions (GFP or luciferase) are typically lost by ∼p2–p5 across cell types (Pierson et al., 2005; Yun et al., 2020; Zhang et al., 2020).

The preferential excision of fluorescent reporters relative to bioluminescent reporters is associated with a higher GC content. *Aequorea victoria*–derived GFP variants, mCherry or Turbo-GFP are GC-rich (∼62-64% GC), whereas commonly used bioluminescent reporters have lower GC content (Table S4). Notably, Nluc offers a favorable combination of shorter length (515 nt) and GC content closer to the viral background (∼54%). Additionally, the lower recovery efficiency reported for DENV2 and DENV4 relative to ZIKV or KUNV is consistent with differences in viral genome GC content (Table S4) (Cherkashchenko et al., 2023). We therefore suggest that adjusting reporter GC content toward the viral genome could improve reporter gene stability in recombinant orthoflaviviruses, as reported in other virus families (Kanai et al., 2023), and reduce zinc-finger antiviral protein (ZAP)-associated selective pressure on flavivirus genomes (Fros et al., 2021; Van Bree et al., 2025).

Importantly, recombination and stability are context-dependent and extend beyond reporter sequence composition or reported variations between viruses (Cherkashchenko et al., 2023). Production variables also influence stability: at low MOI, GFP-negative revertants arose in BHK-21 but were not detected in A549/Npro under matched conditions. By contrast, TBEVnluc did not yield Nluc-negative revertants in either cell line within the experimental timeframe tested. These findings underscore the impact of MOI, harvest timing, and producer cell line on the genetic stability of reporter viruses. Accordingly, routine quality control by long-read sequencing (e.g., Oxford Nanopore Technology) of virus stocks can help confirm reporter integrity and detect emerging variants. A limitation of the present study is that the relative contributions of reporter identity, sequence composition, and genomic context to stability remain to be resolved. This will require matched serial-passaging experiments with GFP, Nluc and other non-GFP fluorescent reporters to quantify reporter-negative outgrowth and to map deletion junctions, together with direct comparisons of wild-type GFP versus GC-and codon-tuned GFP variants in recombinant flavivirus genomes.

Our data and prior work suggest that (i) GC-rich GFP variants (and alternative fluorescent proteins such as mCherry) are prone to recombination in capsid-insertion designs; (ii) bioluminescent reporters can be more stable; and (iii) protective configurations such as dual 2A linkers, recombination “dead-ends” and duplicated C72wt can cumulatively reduce, but not eliminate, the emergence of reporter-negative genomes (Cherkashchenko et al., 2023; Haviernik et al., 2021; Suphatrakul et al., 2018; Volkova et al., 2020). Given the apparent recombination hotspots within GFP coding sequence in our system, insert choice and genomic context should be tailored to the intended application. When longer-term stability is needed, strategies such as codon tuning toward viral GC content, protective safeguards, E–NS1 insertion designs, and luciferase reporters are particularly attractive.

Overall, our results help to refine practical guidelines for generating recombinant reporter flaviviruses spanning construct design, reporter choice, and recovery conditions. Favoring luciferases when durability is required and tuning MOI, timing, and producer cell lines improve stability at rescue and early passage. These principles generalize to other reporters, with GFP variants particularly prone to loss, and could be used to stabilize or optimize existing GFP-based systems.

## ACKNOWLEDGEMENTS

This work of the Interdisciplinary Thematic Institute IMCbio+, as part of the ITI 2021-2028 program of the University of Strasbourg, CNRS and Inserm, was supported by Agence Nationale de la Recherche through IdEx Unistra (ANR-10-IDEX-0002), SFRI-STRAT’US project (ANR-20-SFRI-0012), and EUR IMCBio (IMCBio ANR-17-EURE-0023) under the framework of the French Investments for the Future Program. It was also supported by an Agence Nationale pour la Recherche PRCI grant to S.P. and C.M. (ANR-21-CE35-0018-01), European Union’s Horizon Europe research and innovation program under the Marie Skłodowska-Curie grant agreement #101109395 (MSCA Postdoctoral Fellowship to B.B.). Additional support for this research was provided by the German Research Foundation (Deutsche Forschungsgemeinschaft, DFG; grant BE 5748/9-1 to S.C.B.) and the National Research Foundation (NRF) funded by the Korean government (MSIT) (RS-2024-00398073). We thank Nicolas Baumberger (University of Strasbourg, CNRS, FR) for producing the Marathon reverse transcriptase. We thank Alain Khol (University of Glasgow, UK) for providing the A549/Npro cell line. We thank Thomas Loustau for providing the CHME-5 cell line.

