## Supplementary material for "Field-isolate recombinant tick-borne encephalitis viruses define reporter-stability guidelines for antiviral testing in flaviviruses": besson_REV_supp_data.docx

### SUPPLEMENTARY TABLES

#### Table S1. Primer list

| **Use** | **Orientation** | **Sequence** | **Region** |
| --- | --- | --- | --- |
| 3'5' linker PCR (cloning)  and 3'5' reporter linker PCR (rescue) | for | GTTCTTGTTCTCCCTGAGCC | 3'UTR |
| 3'5' linker PCR (cloning) | rev | CGTTGCGGTCTCTTTCGACAC | C gene |
| Complete TBEV genomic fragment 1 PCR (sequencing)  and reporter PCR (analysis) | for | AGATTTTCTTGCACGTGCATGC | 5'UTR |
| reporter 2A PCR (cloning) | for | GTGTCGAAAGAGACCGCAACG | C gene |
| 3'5' reporter linker (rescue)  and reporter PCR (analysis) | rev | CTACGGCATGCCACAAGATC | C gene |
| reporter RT (analysis) | rev | GCCAATGCCACTCCTAATAGAAGC | C gene |
| Truncated TBEV genomic fragment 1 PCR (rescue) | for | GTCCAAATGCCAAATGGGCT | C gene |
| TBEV genomic fragment 1 RT and PCR (rescue, sequencing) | rev | CTCAAACACAGCCTGGAGTAGC | NS3 gene |
| TBEV genomic fragment 2 PCR (rescue) | for | GATCACATTCCACCTTGAGCTAGG | NS3 gene |
| TBEV genomic fragment 2 RT and PCR (rescue, sequencing) | rev | CACTATGGCCAACACCAGACTC | NS5 gene |
| TBEV genomic fragment 3 PCR (rescue, sequencing) | for | GGATGTCATCAACCCATTCGGG | NS5 gene |
| 3'UTR PCR (cloning) | for | CTCAGACTGGAGAGCTCAATAATC | 3'UTR |
| TBEV genomic fragment 3 RT, PCR (rescue, sequencing) 3'UTR PCR (cloning) and qPCR RT (analysis) | rev | AGCGGGTGTTTTTCCGAGTC | 3'UTR |
| TBEV qPCR (analysis) | for | ACTCACAGAGATAGAGCTCGGA | 3'UTR |
| TBEV qPCR (analysis) | rev | CCCACAAACCGCTATGAAGC | 3'UTR |

#### Table S2. TBEV-Eu genome references used

| **Strain** | **GenBank** |
| --- | --- |
| Haselmuhl tiho1 | PZ023949 |
| Neudoerfl | NC_001672 |
| Hypr | OP037819 |
| Salem | FJ572210 |
| Salland | ON502378 |
| Sipoo-8 | MK801805 |
| NL | LC171402 |
| N5-17 | MG243699 |
| KEM-127 | MG210947 |
| Absettarov | KJ000002 |
| K23 | AM600965 |
| BaWa11-171 | KX268728 |
| Oshima | AB753012 |
| SofjinKSY | JF819648.2 |

#### Table S3. Complete TBEV Haselmühl wild-type and recombinant system sequences

| **Name** | **Sequence** |
| --- | --- |
| Haselmühl tiho1 complete genome | agattttcttgcacgtgcatgcgtttgcttcggacagcaatagcagcggttggtttgaaagagatattcttttgtttctaccagtcgtgaacgtgttgagaaaaagacagcttaggagaacaagagctggggATGGTCAAGAAGGCCATCCTGAAAGGTAAGGGGGGCGGTCCCCCTCGACGAGTGTCGAAAGAGACCGCAACGAAGACGCGTCAACCCAGAGTCCAAATGCCAAATGGGCTTGTGTTGATGCGCATGATGGGGATCTTGTGGCATGCCGTAGCTGGCACCGCGAGAAACCCCGTATTGAAGGCGTTTTGGAACTCGGTCCCTCTGAAACAGGCCACAGCAGCACTGCGGAAGATCAAAAGGACGGTGAGTGCTCTCATGGTTGGCTTGCAAAAACGTGGGAAAAGGAGGTCAGCGACGGACTGGATGAGCTGGTTGCTGGTCATTACTCTGTTGGGGATGACGCTTGCTGCAACGGTGAGGAAAGAAAGGGACGGCTCAACTGTGATCAGAGCTGAAGGAAAGGATGCAGCAACTCAGGTGCGTGTGGAGAATGGCACCTGTGTGATCCTGGCTACTGACATGGGGTCATGGTGTGATGATTCACTGTCCTATGAGTGTGTGACCATAGATCAAGGAGAGGAGCCTGTTGACGTGGACTGCTTTTGCCGGAACGTTGATGGAGTCTATCTGGAGTATGGACGCTGTGGGAAACAGGAAGGCTCACGGACAAGGCGCTCAGTGCTGATCCCATCCCATGCTCAGGGAGAGCTGACGGGGAGGGGACACAAATGGCTAGAAGGAGACTCGCTGCGAACACACCTCACTAGAGTTGAGGGATGGGTCTGGAAGAACAAGCTACTTGCCTTGGCGATGGTTACCGTTGTGTGGTTGACCCTGGAGAGTGTGGTGACCAGGGTCGCCGTTCTGGTTGTGCTCCTGTGTTTGGCACCGGTCTACGCTTCTCGTTGCACACACTTGGAAAACAGGGACTTTGTGACTGGAACTCAGGGGACTACGAGGGTCACCTTGGTGCTGGAACTGGGTGGATGTGTTACCATAACAGCTGAGGGGAAGCCTTCGATGGATGTGTGGCTTGACGCCATTTACCAGGAGAACCCTGCTAAGACACGTGAGTACTGTTTGCACGCCAAGTTGTCGGACACTAAGGTTGCAGCCAGATGTCCAACAATGGGACCAGCCACTTTGGCTGAAGAACACCAGGGTGGCACAGTGTGCAAGAGAGATCAGAGTGATCGAGGCTGGGGCAACCACTGTGGACTGTTTGGAAAGGGTAGCATTGTGGCCTGTGTCAAGGCGGCTTGTGAGGCAAAAAAGAAAGCCACAGGACATGTGTACGACGCCAACAAAATAGTGTACACGGTTAAAGTCGAACCACACACGGGAGACTATGTTGCCGCAAACGAGACACACAGTGGGAGGAAGACGGCATCCTTCACGGTCTCTTCAGAGAAAACCATTCTAACTATGGGTGAATATGGAGATGTGTCTTTGTTGTGCAGGGTCGCTAGTGGCGTTGACTTGGCCCAGACCGTCATCCTTGAGCTTGACAAGACAGTGGAACACCTTCCAACGGCTTGGCAGGTCCACAGGGACTGGTTCAATGATCTGGCTCTGCCATGGAAACATGAGGGAGCGCAAAACTGGAACAACGCAGAAAGACTGGTTGAATTTGGGGCTCCTCACGCTGTCAAGATGGACGTGTACAACCTCGGAGACCAGACTGGAGTGTTACTGAAGGCTCTCGCTGGGGTTCCTGTGGCACACATTGAGGGAACCAAGTACCACCTGAAGAGTGGCCACGTGACCTGCGAAGTGGGACTGGAAAAACTGAAGATGAAAGGTCTTACGTACACAATGTGTGACAAAACAAAGTTCACATGGAAGAGAGCTCCAACAGACAGTGGGCATGATACAGTGGTCATGGAAGTCACATTCTCTGGAACAAAGCCCTGTAGGATCCCAGTCAGGGCAGTGGCACATGGATCTCCAGATGTGAACGTGGCCATGCTGATAACGCCAAACCCAACAATTGAAAACAATGGAGGTGGCTTCATAGAGATGCAGCTGCCCCCAGGGGATAACATCATCTATGTTGGGGAACTGAGTCATCAATGGTTCCAAAAAGGGAGCAGCATCGGAAGGGTTTTCCAAAAGACCAAGAAAGGCATAGAAAGATTGACAGTGATAGGAGAGCACGCCTGGGACTTCGGTTCTGCTGGAGGCTTTCTGAGTTCAATTGGGAAGGCGGTGCACACGGTCCTTGGTGGTGCTTTCAACAGCATCTTCGGGGGAGTGGGGTTTCTACCAAAGCTTCTATTAGGAGTGGCATTGGCTTGGTTGGGCCTGAACATGAGAAACCCTACAATGTCCATGAGCTTTCTCTTGGCTGGAGGTCTGGTCTTGGCCATGACCCTTGGAGTGGGGGCTGATGTTGGCTGCGCTGTGGACACGGAACGAATGGAGCTCCGCTGTGGCGAGGGCCTGGTCGTGTGGAGAGAGGTCTCAGAATGGTATGACAACTATGCCTACTACCCGGAGACACCGGGGGCCCTTGCATCAGCCATAAAGGAGACATTTGAGGAGGGAAGTTGTGGCGTAGTCCCCCAGAACAGGCTCGAGATGGCCATGTGGAGAAACTCGGTCACAGAGCTGAATCTGGCTCTGGCGGAAGGAGAGGCAAATCTCACAGTGGTGGTGGACAAGTTTGACCCCACTGACTACCGAGGTGGTGTCCCTGGTTTACTGAAAAAAGGAAAGGACATAAAAGTCTCCTGGAAAAGCTGGGGCCATTCAATGATCTGGAGCATTCCTGAGGCCCCCCGTCGCTTCATGGTGGGCACGGAAGGACAAAGTGAGTGTCCCCTAGAGAGACGGAAGACAGGTGTTTTCACGGTGGCAGAATTCGGGGTTGGCCTGAGAACAAAGGTCTTCTTGGATTTCAGACAGGAACCAACACATGAGTGTGACACAGGAGTGATGGGAGCTGCCGTCAAGAACGGCATGGCAGTCCACACAGATCAAAGTCTCTGGATGAGGTCAATGAAAAATGACACAGGCACTTACATAGTTGAACTTTTGGTCACTGACCTGAGGAACTGCTCGTGGCCTGCTAGCCACACTATCGATAATGCTGACGTGGTGGACTCGGAGTTATTCCTTCCGGCGAGCCTGGCAGGACCCAGATCCTGGTACAACAGGATACCTGGTTATTCAGAGCAGGTGAAAGGGCCATGGAAGTACACGCCTATCCGAGTCATCAGAGAGGAGTGTCCCGGCACGACCGTTACCATCAACGCCAAGTGTGACAAAAGAGGAGCATCTGTGAGGAGTACCACAGAGAGCGGCAAGGTTATCCCAGAATGGTGCTGCCGAGCATGCACAATGCCACCAGTGACGTTCCGGACTGGAACTGATTGCTGGTATGCCATGGAAATACGGCCAGTCCATGACCAGGGGGGGCTTGTTCGCTCAATGGTGGTTGCGGACAATGGTGAATTACTTAGTGAGGGAGGGGTCCCCGGAATAGTGGCATTGTTTGTGGTCCTTGAATACATCATCCGTAGGAGACCCTCCACGGGAGCGACGGTTGTGTGGGGGGGCATCGTCGTTCTCGCTCTGCTTGTCACCGGGATGGTCAGGATAGAGAGCCTGGTGCGCTATGTGGTGGCAGTGGGGATCACATTCCACCTTGAGCTAGGGCCAGAGATCGTGGCATTGATGCTACTCCAGGCTGTGTTTGAGCTGAGGGTGGGTTTGCTCAGCGCATTTGCACTGCGTAGAAGCCTCACCGTCCGAGAGATGGTGACCACCTACTTTCTCTTGCTGGTCCTGGAATTGGGGCTGCCGGGTGCGAGCCTTGAGGATTTCTGGAAATGGGGTGATGCACTGGCTATGGGGGCGCTGATATTCAGGGCTTGCACGGCGGAAGGAAAGACTGGAGCGGGGCTCTTGCTCATGGCTCTCATGACACAGCAGGATGTGGTGACTGTGCATCATGGACTGGTGTGCTTCCTGAGTGTAGCTTCGGCTTGCTCGGTCTGGAGGCTGCTCAAGGGACACAGAGAGCAGAAGGGATTGACCTGGATTGTCCCCCTGGCTAGATTGCTTGGGGGAGAGGGCTCTGGAATCAGACTGCTGGCGTTTTGGGAGCTGTCAGCTCGCAGAGGGAGACGATCCTTCAGTGAACCACTAACTGTGGTAGGAGTCATGCTAACACTGGCCAGCGGCATGATGCGACACACCTCCCAGGAGGCTCTCTGTGCACTCGCAGTGGCCTCGTTTCTCTTGTTGATGCTGGTGCTGGGGACAAGAAAGATGCAGCTGGTTGCCGAATGGAGTGGCTGCGTGGAATGGCACCCGGAACTAGTGAATGAGGGTGGAGAGGTTAGCCTGCGGGTCCGTCAGGACGCGATGGGAAACTTTCACTTGACTGAGCTCGAGAAAGAAGAGAGAATGATGGCTTTTTGGCTGCTTGCCGGCTTGGCAGCCTCGGCCATTCATTGGTCAGGCATTCTTGGTGTGATGGGACTGTGGACGCTCACGGAAATGCTGAGGTCATCCCGAAGGTCTGACCTGATTTTCTCTGGACAGGGGGGTCGAGAGCGTGGTGACAGACCTTTCGAGGTTAGGGACGGTGTCTACAGGATTTTCAGCCCCGGCTTGTTCTGGGGTCAGAACCAGGTGGGAGTTGGCTACGGTTCCAAGGGTGTTTTACACACGATGTGGCACGTGACAAGAGGAGCGGCGCTGTCTATTGATGATGCTGTAGCCGGTCCCTACTGGGCTGATGTGAGGGAAGATGTCGTGTGCTACGGAGGAGCCTGGAGTCTGGAGGAAAAATGGAAAGGTGAAACAGTACAGGTCCATGCCTTCCCACCGGGGAAGGCCCATGAGGTGCATCAGTGCCAGCCTGGGGAGTTGATCCTTGACACCGGAAGGAAGCTCGGGGCAATACCAATTGATTTGGTAAAAGGAACATCAGGCAGCCCCATTCTTAACGCCCAGGGAGTGGTTGTGGGGCTATATGGAAATGGCCTAAAAACTAATGAGACCTACGTCAGCAGCATTGCTCAAGGAGAAGCAGAGAGGAGTCGACCCAACCTTCCACGGGCTGTTGTGGGTACTGGCTGGACATCAAAGGGTCAGATCACAGTGCTGGACATGCACCCAGGCTCGGGGAAGACCCACAGAGTCCTCCCGGAGCTCATTCGCCAATGCATTGACAGGCGCCTGAGAACGTTGGTGTTGGCTCCAACTCGTGTGGTACTCAAAGAAATGGAGCGCGCCTTGAATGGGAAACGGGTCAGGTTCCACTCACCAGCAGTCAGTGACCAACAGGCTGGAGGGGCAATTGTCGATGTGATGTGTCACGCAACCTATGTCAACAGAAGGCTACTCCCACAGGGGAGACAAAATTGGGAGGTGGCAATCATGGATGAGGCCCACTGGACGGACCCCCACAGCATAGCTGCCAGAGGTCATTTGTACACTCTGGCAAAAGAAAACAAGTGTGCACTGGTCTTGATGACAGCGACACCTCCTGGCAAGAGTGAACCCTTTCCGGAGTCTAACGGAGCCATTACTAGTGAGGAAAGACAGATTCCTGATGGAGAGTGGCGCGACGGGTTTGACTGGATCACTGAGTATGAAGGGCGCACCGCTTGGTTTGTCCCTTCGATTGCAAAAGGTGGGGCTATAGCTCGCACCTTGAGACAGAAGGGGAAAAGTGTGATCTGTTTGAACAGCAAAACCTTTGAAAAGGACTACTCCAGAGTGAGGGATGAGAAGCCTGACTTTGTGGTGACGACTGATATTTCGGAGATGGGAGCCAACCTTGACGTGAGCCGCGTCATAGATGGGAGGACAAACATCAAGCCTGAAGAGGTTGATGGGAAGGTCGAGCTCACCGGGACCAGGCGAGTGACCACGGCTTCCGCTGCCCAACGGCGCGGGAGAGTTGGTCGGCAAGACGGACGAACAGATGAATACATATACTCTGGACAGTGTGATGATGATGACAGTGGATTAGTGCAATGGAAAGAGGCGCAAATACTTCTTGACAACATAACAACCTTGCGGGGGCCCGTGGCCACCTTCTATGGACCAGAACAAGACAAGATGCCGGAGGTGGCTGGTCACTTTCGACTCACTGAAGAGAAAAGAAAGCACTTCCGACATCTTCTCACCCATTGTGACTTCACACCGTGGCTGGCATGGCACGTCGCAGCAAATGTGTCCAGCGTCACGGATCGAAGCTGGACATGGGAAGGGCCGGAGGCAAATGCCGTGGATGAGGCCAGTGGTGACTTGGTCACCTTTAGGAGCCCGAATGGGGCGGAGAGAACTCTCAGGCCGGTGTGGAAGGACGCACGCATGTTCAAAGAGGGACGTGACATCAAAGAGTTCGTGGCGTACGCGTCTGGGCGTCGCAGCTTTGGAGATGTTCTGACAGGAATGTCGGGAGTTCCTGAGCTTTTGCGGCACAGGTGCGTCAGTGCCCTGGATGTCTTCTACACACTTATGCATGAGGAACCTGGCAGCAGGGCAATGAGAATGGCGGAGAGAGATGCCCCAGAGGCCTTTCTGACTATGGTTGAGATGATGGTACTGGGCTTGGCAACCCTGGGTGTCATCTGGTGCTTCGTCGTCCGGACTTCAATCAGCCGTATGATGCTGGGCACGCTGGTCCTGCTGGCCTCCTTGCTACTTTTGTGGGCAGGTGGCGTCGGCTATGGAAATATGGCCGGAGTGGCTCTCATCTTTTACACGTTGCTGACAGTGCTGCAGCCCGAGGCGGGAAAACAGAGAAGCAGTGACGACAACAAACTGGCATATTTCTTGCTGACGCTCTGCAGCCTTGCTGGACTGGTTGCAGCCAATGAGATGGGTTTTCTGGAGAAGACCAAGGCAGACTTGTCCACGGTGCTGTGGAGTGAACAGGAGGAACCCCGGCCATGGAGTGAATGGACGAATGTGGACATCCAGCCAGCGAGGTCCTGGGGGACCTACGTGCTGGTGGTGTCTCTGTTTACACCGTACATCATCCACCAACTGCAAACCAAAATCCAACAACTTGTCAACAGTGCCGTGGCATCTGGTGCACAGGCCATGAGAGACCTTGGGGGAGGTGCCCCCTTCTTTGGTGTGGCGGGACATGTCATGACCCTCGGGGTGGTGTCACTGATTGGGGCTACTCCCACCTCACTGATGGTGGGCGTTGGCTTGGCGGCACTCCATCTGGCCATTGTGGTGTCTGGCCTGGAGGCTGAATTGACACAGAGAGCTCATAAGGTCTTTTTCTCTGCAATGGTGCGCAACCCCATGGTGGATGGGGATGTCATCAACCCATTCGGGGAGGGGGAGGCAAAACCTGCTCTATATGAAAGGAAAATGAGTCTGGTGTTGGCCATAGTGTTGTGCCTCATGTCGGTGGTCATGAACCGAACGGTGGCTTCCATAACAGAAGCTTCAGCTGTGGGACTGGCAGCAGCGGGACAGCTGCTTAGACCGGAGGCTGACACGCTGTGGACGATGCCGGTTGCTTGTGGCATGAGTGGTGTGGTCAGGGGTAGCCTGTGGGGGTTTCTGCCTCTTGGGCATAGACTCTGGCTTCGAGCTTCTGGGGGCAGACGTGGCGGTTCTGAGGGAGACACGCTTGGAGATCTCTGGAAACGGAGGCTGAACAACTGCACCAGGGAGGAATTCTTTGTGTACAGGCGCACTGGCATCCTTGAGACGGAACGTGACAAGGCTAGAGAGTTGCTCAGAAGAGGAGAGACCAATATGGGATTGGCTGTCTCTCGGGGCACGGCAAAGCTTGCCTGGCTTGAGGAACGCGGATATGCCACCCTCAAGGGAGAGGTGGTAGATCTTGGATGTGGAAGGGGCGGCTGGTCCTACTATGCGGCATCCCGACCGGCAGTCATGAGTGTCAGGGCGTACACCATTGGTGGAAGAGGGCACGAGGCTCCAAAGATGGTAACAAGCCTGGGTTGGAACTTGATTAAATTTAGATCAGGAATGGACGTGTTCAGCATGCAGCCACACCGGGCTGACACTGTCATGTGTGACATCGGAGAGAGCAGCCCAGATGCCGCTGTGGAAGGTGAGAGGACAAGGAAAGTGATATTGCTCATGGAGCAATGGAAAAACAGGAATCCCACGGCTGCCTGTGTGTTCAAGGTGCTGGCCCCATACCGCCCAGAAGTGATAGAAGCACTGCACAGATTCCAACTGCAGTGGGGGGGGGGTCTGGTGAGGACCCCTTTTTCAAGGAACTCCACCCATGAGATGTATTACTCAACAGCTGTCACTGGGAACATAGTGAACTCCGTCAATGTACAGTCGAGGAAACTTTTGGCTCGGTTTGGAGACCAGAGAGGGCCAACCAGGGTGCCTGAACTTGACCTGGGAGTTGGAACGAGGTGTGTGGTCTTAGCTGAGGACAAGGTGAAAGAACAAGACGTACAAGAGAGGATCAGAGCGTTGCGTGAGCAATACAGCGAAACCTGGCATATGGACGAGGAACACCCGTACCGGACATGGCAGTATTGGGGCAGCTACCGCACGGCACCAACCGGCTCGGCGGCGTCACTGATCAATGGAGTTGTGAAACTTCTCAGCTGGCCATGGAACGCACGGGAAGATGTGGTGCGCATGGCTATGACTGACACAACGGCTTTTGGACAGCAGAGAGTGTTCAAGGACAAAGTTGACACAAAGGCACAGGAGCCTCAGCCCGGTACAAGAGTCATCATGAGAGCAGTAAATGATTGGATTTTGGAACGACTGGCGCAGAAAAGCAAACCGCGCATGTGCAGCAAAGAAGAATTCATAGCAAAAGTGAAATCAAATGCAGCCTTGGGAGCTTGGTCAGATGAGCAAAACAGATGGGCAAGTGCAAGAGAGGCTGTAGAGGATCCTGCATTTTGGCACCTCGTGGATGAAGAGAGAGAAAGGCACCTCATGGGGAGATGCGCGCACTGCGTGTACAACATGATGGGCAAGAGAGAGAAGAAACTGGGAGAGTTCGGAGTGGCAAAAGGAAGTCGGGCCATTTGGTACATGTGGCTGGGGAGTCGCTTTCTGGAGTTCGAAGCTCTTGGATTCTTGAATGAAGACCATTGGGCCTCTAGAGAGTCCAGTGGAGCTGGAGTTGAAGGAATAAGCTTGAACTACCTGGGCTGGCACCTCAAGAAGTTGTCAACCCTGAATGGAGGACTCTTCTATGCAGATGACACAGCTGGCTGGGACACGAAAGTTACCAATGCAGACTTAGAGGATGAAGAACAGATCCTACGGTACATGGAGGGTGAGCACAAACAATTGGCAACCACAATAATGCAAAAAGCATACCATGCCAAAGTCGTGAAGGTCGCGAGGCCCTCCCGTGATGGAGGCTGCATCATGGATGTCATCACAAGAAGAGACCAAAGAGGCTCGGGCCAGGTTGTGACCTATGCCCTTAACACCCTCACCAACATAAAGGTGCAACTAATCCGAATGATGGAAGGGGAAGGGGTCATAGAGGCAGCGGATGCACACAACCCGAGACTGCTTCGAGTGGAGCGCTGGCTGAAAGAACATGGAGAAGAGCGTCTTGGAAGAATGCTCGTTAGTGGTGACGATTGTGTGGTGAGGCCCTTGGATGACAGATTTGGCAAAGCACTCTACTTTCTGAATGACATGGCCAAGACCAGGAAGGACATTGGGGAATGGGAGCACTCGGCCGGCTTTTCAAGCTGGGAGGAGGTCCCCTTTTGTTCACACCATTTCCACGAGCTAGTGATGAAGGACGGACGCACCCTGGTGGTGCCGTGCCGAGACCAAGATGAACTCGTTGGGAGGGCGCGCATCTCACCGGGGTGCGGCTGGAGTGTCCGCGAGACGGCCTGCCTTTCAAAAGCCTACGGGCAGATGTGGCTGCTGAGCTATTTCCACCGGCGAGACCTGAGGACGCTCGGGCTCGCCATCAACTCAGCAGTGCCTGTCGATTGGGTTCCCACCGGCCGCACGACGTGGAGCATCCATGCCAGTGGGGCCTGGATGACCACAGAAGACATGCTGGACGTTTGGAACCGGGTGTGGATCCTGGACAACCCTTTCATGCAGAACAAGGAAAAGGTCATGGAGTGGAGGGATGTTCCGTACCTCCCTAAAGCTCAGGACATGTTATGTTCCTCCCTTGTTGGGAGGAGAGAAAGAGCAGAATGGGCTAAGAACATCTGGGGAGCGGTGGAAAAGGTGAGGAAGATGATAGGTCCTGAAAAGTTCAAGGACTATCTCTCCTGTATGGACCGCCATGACCTGCACTGGGAGCTCAGACTGGAGAGCTCAATAATCtaaacccagactgtgacagagcaaaacccggagggctcgcaaaagattgtccggaaccaaaacagaagcaagcaactcacagagatagagctcggactggagagctctttaaacaaaaaagccagaattgagctgaacctggagagctcattaaatatagtccagacaaaacaaaacatgacaaagtaaataggctgagctaaaagctcccaccacgggactgcttcatagcggtttgtggggggaggctaggaggcgaagccacagatcatggagtgatgcggcagcgcgcgagagcgacggggaagtggtcgcacccgacgcactatccatgaagcaatacttcgtgagacccccctggccagcaaagggggcagactggtcaggggtaagggatgcccccagagtgcattacggcagcacgccagtgagagtggcgacgggaaaatggtcgatcccgacgtagggcactctgaaaaattttgtgagaccccctgcatcatgacaaggccgaacatggtgcatgaaaaggggaggcccccggaagcacgcttccgggaggagggaagagagaaattggcagctctcttcaggatttttcctcctcctatacaaaattccccctcggtagagggggggcggttcttgttctccctgagccaccatcacccagacacagatagtctgacaaggaggtgatgtgtgactcggaaaaacacccgct |
| circular TBEVgfp | GTGATGCGGTTTTGGCAGTACATCAATGGGCGTGGATAGCGGTTTGACTCACGGGGATTTCCAAGTCTCCACCCCATTGACGTCAATGGGAGTTTGTTTTGGCACCAAAATCAACGGGACTTTCCAAAATGTCGTAACAACTCCGCCCCATTGACGCAAATGGGCGGTAGGCGTGTACGGTGGGAGGTCTATATAAGCAGAGCTGGTTTAGTGAACCGagattttcttgcacgtgcatgcgtttgcttcggacagcaatagcagcggttggtttgaaagagatattcttttgtttctaccagtcgtgaacgtgttgagaaaaagacagcttaggagaacaagagctggggATGGTCAAGAAGGCCATCCTGAAAGGTAAGGGGGGCGGTCCCCCTCGACGAGTGTCGAAAGAGACCGCAACGatggtgagcaagggcgaggagctgttcaccggggtggtgcccatcctggtcgagctggacggcgacgtaaacggccacaagttcagcgtgtccggcgagggcgagggcgatgccacctacggcaagctgaccctgaagttcatctgcaccaccggcaagctgcccgtgccctggcccaccctcgtgaccaccctgacctacggcgtgcagtgcttcagccgctaccccgaccacatgaagcagcacgacttcttcaagtccgccatgcccgaaggctacgtccaggagcgcaccatcttcttcaaggacgacggcaactacaagacccgcgccgaggtgaagttcgagggcgacaccctggtgaaccgcatcgagctgaagggcatcgacttcaaggaggacggcaacatcctggggcacaagctggagtacaactacaacagccacaacgtctatatcatggccgacaagcagaagaacggcatcaaggtgaacttcaagatccgccacaacatcgaggacggcagcgtgcagctcgccgaccactaccagcagaacacccccatcggcgacggccccgtgctgctgcccgacaaccactacctgagcacccagtccgccctgagcaaagaccccaacgagaagcgcgatcacatggtcctgctggagttcgtgaccgccgccgggatcactctcggcatggacgagctgtacaagggaagcggagctactaacttcagcctgctgaagcaggctggagacgtggaggagaaccctggacctATGGTGAAAAAAGCTATTCTCAAGGGCAAAGGCGGAGGACCTCCAAGGAGGGTCAGCAAGGAAACAGCCACCAAGACGCGTCAACCCAGAGTCCAAATGCCAAATGGGCTTGTGTTGATGCGCATGATGGGGATCTTGTGGCATGCCGTAGCTGGCACCGCGAGAAACCCCGTATTGAAGGCGTTTTGGAACTCGGTCCCTCTGAAACAGGCCACAGCAGCACTGCGGAAGATCAAAAGGACGGTGAGTGCTCTCATGGTTGGCTTGCAAAAACGTGGGAAAAGGAGGTCAGCGACGGACTGGATGAGCTGGTTGCTGGTCATTACTCTGTTGGGGATGACGCTTGCTGCAACGGTGAGGAAAGAAAGGGACGGCTCAACTGTGATCAGAGCTGAAGGAAAGGATGCAGCAACTCAGGTGCGTGTGGAGAATGGCACCTGTGTGATCCTGGCTACTGACATGGGGTCATGGTGTGATGATTCACTGTCCTATGAGTGTGTGACCATAGATCAAGGAGAGGAGCCTGTTGACGTGGACTGCTTTTGCCGGAACGTTGATGGAGTCTATCTGGAGTATGGACGCTGTGGGAAACAGGAAGGCTCACGGACAAGGCGCTCAGTGCTGATCCCATCCCATGCTCAGGGAGAGCTGACGGGGAGGGGACACAAATGGCTAGAAGGAGACTCGCTGCGAACACACCTCACTAGAGTTGAGGGATGGGTCTGGAAGAACAAGCTACTTGCCTTGGCGATGGTTACCGTTGTGTGGTTGACCCTGGAGAGTGTGGTGACCAGGGTCGCCGTTCTGGTTGTGCTCCTGTGTTTGGCACCGGTCTACGCTTCTCGTTGCACACACTTGGAAAACAGGGACTTTGTGACTGGAACTCAGGGGACTACGAGGGTCACCTTGGTGCTGGAACTGGGTGGATGTGTTACCATAACAGCTGAGGGGAAGCCTTCGATGGATGTGTGGCTTGACGCCATTTACCAGGAGAACCCTGCTAAGACACGTGAGTACTGTTTGCACGCCAAGTTGTCGGACACTAAGGTTGCAGCCAGATGTCCAACAATGGGACCAGCCACTTTGGCTGAAGAACACCAGGGTGGCACAGTGTGCAAGAGAGATCAGAGTGATCGAGGCTGGGGCAACCACTGTGGACTGTTTGGAAAGGGTAGCATTGTGGCCTGTGTCAAGGCGGCTTGTGAGGCAAAAAAGAAAGCCACAGGACATGTGTACGACGCCAACAAAATAGTGTACACGGTTAAAGTCGAACCACACACGGGAGACTATGTTGCCGCAAACGAGACACACAGTGGGAGGAAGACGGCATCCTTCACGGTCTCTTCAGAGAAAACCATTCTAACTATGGGTGAATATGGAGATGTGTCTTTGTTGTGCAGGGTCGCTAGTGGCGTTGACTTGGCCCAGACCGTCATCCTTGAGCTTGACAAGACAGTGGAACACCTTCCAACGGCTTGGCAGGTCCACAGGGACTGGTTCAATGATCTGGCTCTGCCATGGAAACATGAGGGAGCGCAAAACTGGAACAACGCAGAAAGACTGGTTGAATTTGGGGCTCCTCACGCTGTCAAGATGGACGTGTACAACCTCGGAGACCAGACTGGAGTGTTACTGAAGGCTCTCGCTGGGGTTCCTGTGGCACACATTGAGGGAACCAAGTACCACCTGAAGAGTGGCCACGTGACCTGCGAAGTGGGACTGGAAAAACTGAAGATGAAAGGTCTTACGTACACAATGTGTGACAAAACAAAGTTCACATGGAAGAGAGCTCCAACAGACAGTGGGCATGATACAGTGGTCATGGAAGTCACATTCTCTGGAACAAAGCCCTGTAGGATCCCAGTCAGGGCAGTGGCACATGGATCTCCAGATGTGAACGTGGCCATGCTGATAACGCCAAACCCAACAATTGAAAACAATGGAGGTGGCTTCATAGAGATGCAGCTGCCCCCAGGGGATAACATCATCTATGTTGGGGAACTGAGTCATCAATGGTTCCAAAAAGGGAGCAGCATCGGAAGGGTTTTCCAAAAGACCAAGAAAGGCATAGAAAGATTGACAGTGATAGGAGAGCACGCCTGGGACTTCGGTTCTGCTGGAGGCTTTCTGAGTTCAATTGGGAAGGCGGTGCACACGGTCCTTGGTGGTGCTTTCAACAGCATCTTCGGGGGAGTGGGGTTTCTACCAAAGCTTCTATTAGGAGTGGCATTGGCTTGGTTGGGCCTGAACATGAGAAACCCTACAATGTCCATGAGCTTTCTCTTGGCTGGAGGTCTGGTCTTGGCCATGACCCTTGGAGTGGGGGCTGATGTTGGCTGCGCTGTGGACACGGAACGAATGGAGCTCCGCTGTGGCGAGGGCCTGGTCGTGTGGAGAGAGGTCTCAGAATGGTATGACAACTATGCCTACTACCCGGAGACACCGGGGGCCCTTGCATCAGCCATAAAGGAGACATTTGAGGAGGGAAGTTGTGGCGTAGTCCCCCAGAACAGGCTCGAGATGGCCATGTGGAGAAACTCGGTCACAGAGCTGAATCTGGCTCTGGCGGAAGGAGAGGCAAATCTCACAGTGGTGGTGGACAAGTTTGACCCCACTGACTACCGAGGTGGTGTCCCTGGTTTACTGAAAAAAGGAAAGGACATAAAAGTCTCCTGGAAAAGCTGGGGCCATTCAATGATCTGGAGCATTCCTGAGGCCCCCCGTCGCTTCATGGTGGGCACGGAAGGACAAAGTGAGTGTCCCCTAGAGAGACGGAAGACAGGTGTTTTCACGGTGGCAGAATTCGGGGTTGGCCTGAGAACAAAGGTCTTCTTGGATTTCAGACAGGAACCAACACATGAGTGTGACACAGGAGTGATGGGAGCTGCCGTCAAGAACGGCATGGCAGTCCACACAGATCAAAGTCTCTGGATGAGGTCAATGAAAAATGACACAGGCACTTACATAGTTGAACTTTTGGTCACTGACCTGAGGAACTGCTCGTGGCCTGCTAGCCACACTATCGATAATGCTGACGTGGTGGACTCGGAGTTATTCCTTCCGGCGAGCCTGGCAGGACCCAGATCCTGGTACAACAGGATACCTGGTTATTCAGAGCAGGTGAAAGGGCCATGGAAGTACACGCCTATCCGAGTCATCAGAGAGGAGTGTCCCGGCACGACCGTTACCATCAACGCCAAGTGTGACAAAAGAGGAGCATCTGTGAGGAGTACCACAGAGAGCGGCAAGGTTATCCCAGAATGGTGCTGCCGAGCATGCACAATGCCACCAGTGACGTTCCGGACTGGAACTGATTGCTGGTATGCCATGGAAATACGGCCAGTCCATGACCAGGGGGGGCTTGTTCGCTCAATGGTGGTTGCGGACAATGGTGAATTACTTAGTGAGGGAGGGGTCCCCGGAATAGTGGCATTGTTTGTGGTCCTTGAATACATCATCCGTAGGAGACCCTCCACGGGAGCGACGGTTGTGTGGGGGGGCATCGTCGTTCTCGCTCTGCTTGTCACCGGGATGGTCAGGATAGAGAGCCTGGTGCGCTATGTGGTGGCAGTGGGGATCACATTCCACCTTGAGCTAGGGCCAGAGATCGTGGCATTGATGCTACTCCAGGCTGTGTTTGAGCTGAGGGTGGGTTTGCTCAGCGCATTTGCACTGCGTAGAAGCCTCACCGTCCGAGAGATGGTGACCACCTACTTTCTCTTGCTGGTCCTGGAATTGGGGCTGCCGGGTGCGAGCCTTGAGGATTTCTGGAAATGGGGTGATGCACTGGCTATGGGGGCGCTGATATTCAGGGCTTGCACGGCGGAAGGAAAGACTGGAGCGGGGCTCTTGCTCATGGCTCTCATGACACAGCAGGATGTGGTGACTGTGCATCATGGACTGGTGTGCTTCCTGAGTGTAGCTTCGGCTTGCTCGGTCTGGAGGCTGCTCAAGGGACACAGAGAGCAGAAGGGATTGACCTGGATTGTCCCCCTGGCTAGATTGCTTGGGGGAGAGGGCTCTGGAATCAGACTGCTGGCGTTTTGGGAGCTGTCAGCTCGCAGAGGGAGACGATCCTTCAGTGAACCACTAACTGTGGTAGGAGTCATGCTAACACTGGCCAGCGGCATGATGCGACACACCTCCCAGGAGGCTCTCTGTGCACTCGCAGTGGCCTCGTTTCTCTTGTTGATGCTGGTGCTGGGGACAAGAAAGATGCAGCTGGTTGCCGAATGGAGTGGCTGCGTGGAATGGCACCCGGAACTAGTGAATGAGGGTGGAGAGGTTAGCCTGCGGGTCCGTCAGGACGCGATGGGAAACTTTCACTTGACTGAGCTCGAGAAAGAAGAGAGAATGATGGCTTTTTGGCTGCTTGCCGGCTTGGCAGCCTCGGCCATTCATTGGTCAGGCATTCTTGGTGTGATGGGACTGTGGACGCTCACGGAAATGCTGAGGTCATCCCGAAGGTCTGACCTGATTTTCTCTGGACAGGGGGGTCGAGAGCGTGGTGACAGACCTTTCGAGGTTAGGGACGGTGTCTACAGGATTTTCAGCCCCGGCTTGTTCTGGGGTCAGAACCAGGTGGGAGTTGGCTACGGTTCCAAGGGTGTTTTACACACGATGTGGCACGTGACAAGAGGAGCGGCGCTGTCTATTGATGATGCTGTAGCCGGTCCCTACTGGGCTGATGTGAGGGAAGATGTCGTGTGCTACGGAGGAGCCTGGAGTCTGGAGGAAAAATGGAAAGGTGAAACAGTACAGGTCCATGCCTTCCCACCGGGGAAGGCCCATGAGGTGCATCAGTGCCAGCCTGGGGAGTTGATCCTTGACACCGGAAGGAAGCTCGGGGCAATACCAATTGATTTGGTAAAAGGAACATCAGGCAGCCCCATTCTTAACGCCCAGGGAGTGGTTGTGGGGCTATATGGAAATGGCCTAAAAACTAATGAGACCTACGTCAGCAGCATTGCTCAAGGAGAAGCAGAGAGGAGTCGACCCAACCTTCCACGGGCTGTTGTGGGTACTGGCTGGACATCAAAGGGTCAGATCACAGTGCTGGACATGCACCCAGGCTCGGGGAAGACCCACAGAGTCCTCCCGGAGCTCATTCGCCAATGCATTGACAGGCGCCTGAGAACGTTGGTGTTGGCTCCAACTCGTGTGGTACTCAAAGAAATGGAGCGCGCCTTGAATGGGAAACGGGTCAGGTTCCACTCACCAGCAGTCAGTGACCAACAGGCTGGAGGGGCAATTGTCGATGTGATGTGTCACGCAACCTATGTCAACAGAAGGCTACTCCCACAGGGGAGACAAAATTGGGAGGTGGCAATCATGGATGAGGCCCACTGGACGGACCCCCACAGCATAGCTGCCAGAGGTCATTTGTACACTCTGGCAAAAGAAAACAAGTGTGCACTGGTCTTGATGACAGCGACACCTCCTGGCAAGAGTGAACCCTTTCCGGAGTCTAACGGAGCCATTACTAGTGAGGAAAGACAGATTCCTGATGGAGAGTGGCGCGACGGGTTTGACTGGATCACTGAGTATGAAGGGCGCACCGCTTGGTTTGTCCCTTCGATTGCAAAAGGTGGGGCTATAGCTCGCACCTTGAGACAGAAGGGGAAAAGTGTGATCTGTTTGAACAGCAAAACCTTTGAAAAGGACTACTCCAGAGTGAGGGATGAGAAGCCTGACTTTGTGGTGACGACTGATATTTCGGAGATGGGAGCCAACCTTGACGTGAGCCGCGTCATAGATGGGAGGACAAACATCAAGCCTGAAGAGGTTGATGGGAAGGTCGAGCTCACCGGGACCAGGCGAGTGACCACGGCTTCCGCTGCCCAACGGCGCGGGAGAGTTGGTCGGCAAGACGGACGAACAGATGAATACATATACTCTGGACAGTGTGATGATGATGACAGTGGATTAGTGCAATGGAAAGAGGCGCAAATACTTCTTGACAACATAACAACCTTGCGGGGGCCCGTGGCCACCTTCTATGGACCAGAACAAGACAAGATGCCGGAGGTGGCTGGTCACTTTCGACTCACTGAAGAGAAAAGAAAGCACTTCCGACATCTTCTCACCCATTGTGACTTCACACCGTGGCTGGCATGGCACGTCGCAGCAAATGTGTCCAGCGTCACGGATCGAAGCTGGACATGGGAAGGGCCGGAGGCAAATGCCGTGGATGAGGCCAGTGGTGACTTGGTCACCTTTAGGAGCCCGAATGGGGCGGAGAGAACTCTCAGGCCGGTGTGGAAGGACGCACGCATGTTCAAAGAGGGACGTGACATCAAAGAGTTCGTGGCGTACGCGTCTGGGCGTCGCAGCTTTGGAGATGTTCTGACAGGAATGTCGGGAGTTCCTGAGCTTTTGCGGCACAGGTGCGTCAGTGCCCTGGATGTCTTCTACACACTTATGCATGAGGAACCTGGCAGCAGGGCAATGAGAATGGCGGAGAGAGATGCCCCAGAGGCCTTTCTGACTATGGTTGAGATGATGGTACTGGGCTTGGCAACCCTGGGTGTCATCTGGTGCTTCGTCGTCCGGACTTCAATCAGCCGTATGATGCTGGGCACGCTGGTCCTGCTGGCCTCCTTGCTACTTTTGTGGGCAGGTGGCGTCGGCTATGGAAATATGGCCGGAGTGGCTCTCATCTTTTACACGTTGCTGACAGTGCTGCAGCCCGAGGCGGGAAAACAGAGAAGCAGTGACGACAACAAACTGGCATATTTCTTGCTGACGCTCTGCAGCCTTGCTGGACTGGTTGCAGCCAATGAGATGGGTTTTCTGGAGAAGACCAAGGCAGACTTGTCCACGGTGCTGTGGAGTGAACAGGAGGAACCCCGGCCATGGAGTGAATGGACGAATGTGGACATCCAGCCAGCGAGGTCCTGGGGGACCTACGTGCTGGTGGTGTCTCTGTTTACACCGTACATCATCCACCAACTGCAAACCAAAATCCAACAACTTGTCAACAGTGCCGTGGCATCTGGTGCACAGGCCATGAGAGACCTTGGGGGAGGTGCCCCCTTCTTTGGTGTGGCGGGACATGTCATGACCCTCGGGGTGGTGTCACTGATTGGGGCTACTCCCACCTCACTGATGGTGGGCGTTGGCTTGGCGGCACTCCATCTGGCCATTGTGGTGTCTGGCCTGGAGGCTGAATTGACACAGAGAGCTCATAAGGTCTTTTTCTCTGCAATGGTGCGCAACCCCATGGTGGATGGGGATGTCATCAACCCATTCGGGGAGGGGGAGGCAAAACCTGCTCTATATGAAAGGAAAATGAGTCTGGTGTTGGCCATAGTGTTGTGCCTCATGTCGGTGGTCATGAACCGAACGGTGGCTTCCATAACAGAAGCTTCAGCTGTGGGACTGGCAGCAGCGGGACAGCTGCTTAGACCGGAGGCTGACACGCTGTGGACGATGCCGGTTGCTTGTGGCATGAGTGGTGTGGTCAGGGGTAGCCTGTGGGGGTTTCTGCCTCTTGGGCATAGACTCTGGCTTCGAGCTTCTGGGGGCAGACGTGGCGGTTCTGAGGGAGACACGCTTGGAGATCTCTGGAAACGGAGGCTGAACAACTGCACCAGGGAGGAATTCTTTGTGTACAGGCGCACTGGCATCCTTGAGACGGAACGTGACAAGGCTAGAGAGTTGCTCAGAAGAGGAGAGACCAATATGGGATTGGCTGTCTCTCGGGGCACGGCAAAGCTTGCCTGGCTTGAGGAACGCGGATATGCCACCCTCAAGGGAGAGGTGGTAGATCTTGGATGTGGAAGGGGCGGCTGGTCCTACTATGCGGCATCCCGACCGGCAGTCATGAGTGTCAGGGCGTACACCATTGGTGGAAGAGGGCACGAGGCTCCAAAGATGGTAACAAGCCTGGGTTGGAACTTGATTAAATTTAGATCAGGAATGGACGTGTTCAGCATGCAGCCACACCGGGCTGACACTGTCATGTGTGACATCGGAGAGAGCAGCCCAGATGCCGCTGTGGAAGGTGAGAGGACAAGGAAAGTGATATTGCTCATGGAGCAATGGAAAAACAGGAATCCCACGGCTGCCTGTGTGTTCAAGGTGCTGGCCCCATACCGCCCAGAAGTGATAGAAGCACTGCACAGATTCCAACTGCAGTGGGGGGGGGGTCTGGTGAGGACCCCTTTTTCAAGGAACTCCACCCATGAGATGTATTACTCAACAGCTGTCACTGGGAACATAGTGAACTCCGTCAATGTACAGTCGAGGAAACTTTTGGCTCGGTTTGGAGACCAGAGAGGGCCAACCAGGGTGCCTGAACTTGACCTGGGAGTTGGAACGAGGTGTGTGGTCTTAGCTGAGGACAAGGTGAAAGAACAAGACGTACAAGAGAGGATCAGAGCGTTGCGTGAGCAATACAGCGAAACCTGGCATATGGACGAGGAACACCCGTACCGGACATGGCAGTATTGGGGCAGCTACCGCACGGCACCAACCGGCTCGGCGGCGTCACTGATCAATGGAGTTGTGAAACTTCTCAGCTGGCCATGGAACGCACGGGAAGATGTGGTGCGCATGGCTATGACTGACACAACGGCTTTTGGACAGCAGAGAGTGTTCAAGGACAAAGTTGACACAAAGGCACAGGAGCCTCAGCCCGGTACAAGAGTCATCATGAGAGCAGTAAATGATTGGATTTTGGAACGACTGGCGCAGAAAAGCAAACCGCGCATGTGCAGCAAAGAAGAATTCATAGCAAAAGTGAAATCAAATGCAGCCTTGGGAGCTTGGTCAGATGAGCAAAACAGATGGGCAAGTGCAAGAGAGGCTGTAGAGGATCCTGCATTTTGGCACCTCGTGGATGAAGAGAGAGAAAGGCACCTCATGGGGAGATGCGCGCACTGCGTGTACAACATGATGGGCAAGAGAGAGAAGAAACTGGGAGAGTTCGGAGTGGCAAAAGGAAGTCGGGCCATTTGGTACATGTGGCTGGGGAGTCGCTTTCTGGAGTTCGAAGCTCTTGGATTCTTGAATGAAGACCATTGGGCCTCTAGAGAGTCCAGTGGAGCTGGAGTTGAAGGAATAAGCTTGAACTACCTGGGCTGGCACCTCAAGAAGTTGTCAACCCTGAATGGAGGACTCTTCTATGCAGATGACACAGCTGGCTGGGACACGAAAGTTACCAATGCAGACTTAGAGGATGAAGAACAGATCCTACGGTACATGGAGGGTGAGCACAAACAATTGGCAACCACAATAATGCAAAAAGCATACCATGCCAAAGTCGTGAAGGTCGCGAGGCCCTCCCGTGATGGAGGCTGCATCATGGATGTCATCACAAGAAGAGACCAAAGAGGCTCGGGCCAGGTTGTGACCTATGCCCTTAACACCCTCACCAACATAAAGGTGCAACTAATCCGAATGATGGAAGGGGAAGGGGTCATAGAGGCAGCGGATGCACACAACCCGAGACTGCTTCGAGTGGAGCGCTGGCTGAAAGAACATGGAGAAGAGCGTCTTGGAAGAATGCTCGTTAGTGGTGACGATTGTGTGGTGAGGCCCTTGGATGACAGATTTGGCAAAGCACTCTACTTTCTGAATGACATGGCCAAGACCAGGAAGGACATTGGGGAATGGGAGCACTCGGCCGGCTTTTCAAGCTGGGAGGAGGTCCCCTTTTGTTCACACCATTTCCACGAGCTAGTGATGAAGGACGGACGCACCCTGGTGGTGCCGTGCCGAGACCAAGATGAACTCGTTGGGAGGGCGCGCATCTCACCGGGGTGCGGCTGGAGTGTCCGCGAGACGGCCTGCCTTTCAAAAGCCTACGGGCAGATGTGGCTGCTGAGCTATTTCCACCGGCGAGACCTGAGGACGCTCGGGCTCGCCATCAACTCAGCAGTGCCTGTCGATTGGGTTCCCACCGGCCGCACGACGTGGAGCATCCATGCCAGTGGGGCCTGGATGACCACAGAAGACATGCTGGACGTTTGGAACCGGGTGTGGATCCTGGACAACCCTTTCATGCAGAACAAGGAAAAGGTCATGGAGTGGAGGGATGTTCCGTACCTCCCTAAAGCTCAGGACATGTTATGTTCCTCCCTTGTTGGGAGGAGAGAAAGAGCAGAATGGGCTAAGAACATCTGGGGAGCGGTGGAAAAGGTGAGGAAGATGATAGGTCCTGAAAAGTTCAAGGACTATCTCTCCTGTATGGACCGCCATGACCTGCACTGGGAGCTCAGACTGGAGAGCTCAATAATCtaaacccagactgtgacagagcaaaacccggagggctcgcaaaagattgtccggaaccaaaacagaagcaagcaactcacagagatagagctcggactggagagctctttaaacaaaaaagccagaattgagctgaacctggagagctcattaaatatagtccagacaaaacaaaacatgacaaagtaaataggctgagctaaaagctcccaccacgggactgcttcatagcggtttgtggggggaggctaggaggcgaagccacagatcatggagtgatgcggcagcgcgcgagagcgacggggaagtggtcgcacccgacgcactatccatgaagcaatacttcgtgagacccccctggccagcaaagggggcagactggtcaggggtaagggatgcccccagagtgcattacggcagcacgccagtgagagtggcgacgggaaaatggtcgatcccgacgtagggcactctgaaaaattttgtgagaccccctgcatcatgacaaggccgaacatggtgcatgaaaaggggaggcccccggaagcacgcttccgggaggagggaagagagaaattggcagctctcttcaggatttttcctcctcctatacaaaattccccctcggtagagggggggcggttcttgttctccctgagccaccatcacccagacacagatagtctgacaaggaggtgatgtgtgactcggaaaaacacccgctGGGTCGGCATGGCATCTCCACCTCCTCGCGGTCCGACCTGGGCATCCGAAGGAGGACGTCGTCCACTCGGATGGCTAAGGGAGAGCTCggatccaccggatctagataactgatcataatcagccataccacatttgtagaggttttacttgctttaaaaaacctcccacacctccccctgaacctgaaacataaaatgaatgcaattgttgttgttAACTTGTTTATTGCAGCTTATAATGGTTACAAATAAAGCAATAGCATCACAAATTTCACAAATAAAGCATTTTTTTCACTGCATTCTAGTTGTGGTTTGTCCAAACTCATCAATGTATCTTAtagtaatcaattacggggtcattagttcatagcccatatatggagttccgCGTTACATAACTTACGGTAAATGGCCCGCCTGGCTGACCGCCCAACGACCCCCGCCCATTGACGTCAATAATGACGTATGTTCCCATAGTAACGCCAATAGGGACTTTCCATTGACGTCAATGGGTGGAGTATTTACGGTAAACTGCCCACTTGGCAGTACATCAAGTGTATCATATGCCAAGTACGCCCCCTATTGACGTCAATGACGGTAAATGGCCCGCCTGGCATTATGCCCAGTACATGACCTTATGGGACTTTCCTACTTGGCAGTACATCTACGTATTAGTCATCGCTATTACCATG |
| infectious TBEVgfp | agattttcttgcacgtgcatgcgtttgcttcggacagcaatagcagcggttggtttgaaagagatattcttttgtttctaccagtcgtgaacgtgttgagaaaaagacagcttaggagaacaagagctggggATGGTCAAGAAGGCCATCCTGAAAGGTAAGGGGGGCGGTCCCCCTCGACGAGTGTCGAAAGAGACCGCAACGatggtgagcaagggcgaggagctgttcaccggggtggtgcccatcctggtcgagctggacggcgacgtaaacggccacaagttcagcgtgtccggcgagggcgagggcgatgccacctacggcaagctgaccctgaagttcatctgcaccaccggcaagctgcccgtgccctggcccaccctcgtgaccaccctgacctacggcgtgcagtgcttcagccgctaccccgaccacatgaagcagcacgacttcttcaagtccgccatgcccgaaggctacgtccaggagcgcaccatcttcttcaaggacgacggcaactacaagacccgcgccgaggtgaagttcgagggcgacaccctggtgaaccgcatcgagctgaagggcatcgacttcaaggaggacggcaacatcctggggcacaagctggagtacaactacaacagccacaacgtctatatcatggccgacaagcagaagaacggcatcaaggtgaacttcaagatccgccacaacatcgaggacggcagcgtgcagctcgccgaccactaccagcagaacacccccatcggcgacggccccgtgctgctgcccgacaaccactacctgagcacccagtccgccctgagcaaagaccccaacgagaagcgcgatcacatggtcctgctggagttcgtgaccgccgccgggatcactctcggcatggacgagctgtacaagggaagcggagctactaacttcagcctgctgaagcaggctggagacgtggaggagaaccctggacctATGGTGAAAAAAGCTATTCTCAAGGGCAAAGGCGGAGGACCTCCAAGGAGGGTCAGCAAGGAAACAGCCACCAAGACGCGTCAACCCAGAGTCCAAATGCCAAATGGGCTTGTGTTGATGCGCATGATGGGGATCTTGTGGCATGCCGTAGCTGGCACCGCGAGAAACCCCGTATTGAAGGCGTTTTGGAACTCGGTCCCTCTGAAACAGGCCACAGCAGCACTGCGGAAGATCAAAAGGACGGTGAGTGCTCTCATGGTTGGCTTGCAAAAACGTGGGAAAAGGAGGTCAGCGACGGACTGGATGAGCTGGTTGCTGGTCATTACTCTGTTGGGGATGACGCTTGCTGCAACGGTGAGGAAAGAAAGGGACGGCTCAACTGTGATCAGAGCTGAAGGAAAGGATGCAGCAACTCAGGTGCGTGTGGAGAATGGCACCTGTGTGATCCTGGCTACTGACATGGGGTCATGGTGTGATGATTCACTGTCCTATGAGTGTGTGACCATAGATCAAGGAGAGGAGCCTGTTGACGTGGACTGCTTTTGCCGGAACGTTGATGGAGTCTATCTGGAGTATGGACGCTGTGGGAAACAGGAAGGCTCACGGACAAGGCGCTCAGTGCTGATCCCATCCCATGCTCAGGGAGAGCTGACGGGGAGGGGACACAAATGGCTAGAAGGAGACTCGCTGCGAACACACCTCACTAGAGTTGAGGGATGGGTCTGGAAGAACAAGCTACTTGCCTTGGCGATGGTTACCGTTGTGTGGTTGACCCTGGAGAGTGTGGTGACCAGGGTCGCCGTTCTGGTTGTGCTCCTGTGTTTGGCACCGGTCTACGCTTCTCGTTGCACACACTTGGAAAACAGGGACTTTGTGACTGGAACTCAGGGGACTACGAGGGTCACCTTGGTGCTGGAACTGGGTGGATGTGTTACCATAACAGCTGAGGGGAAGCCTTCGATGGATGTGTGGCTTGACGCCATTTACCAGGAGAACCCTGCTAAGACACGTGAGTACTGTTTGCACGCCAAGTTGTCGGACACTAAGGTTGCAGCCAGATGTCCAACAATGGGACCAGCCACTTTGGCTGAAGAACACCAGGGTGGCACAGTGTGCAAGAGAGATCAGAGTGATCGAGGCTGGGGCAACCACTGTGGACTGTTTGGAAAGGGTAGCATTGTGGCCTGTGTCAAGGCGGCTTGTGAGGCAAAAAAGAAAGCCACAGGACATGTGTACGACGCCAACAAAATAGTGTACACGGTTAAAGTCGAACCACACACGGGAGACTATGTTGCCGCAAACGAGACACACAGTGGGAGGAAGACGGCATCCTTCACGGTCTCTTCAGAGAAAACCATTCTAACTATGGGTGAATATGGAGATGTGTCTTTGTTGTGCAGGGTCGCTAGTGGCGTTGACTTGGCCCAGACCGTCATCCTTGAGCTTGACAAGACAGTGGAACACCTTCCAACGGCTTGGCAGGTCCACAGGGACTGGTTCAATGATCTGGCTCTGCCATGGAAACATGAGGGAGCGCAAAACTGGAACAACGCAGAAAGACTGGTTGAATTTGGGGCTCCTCACGCTGTCAAGATGGACGTGTACAACCTCGGAGACCAGACTGGAGTGTTACTGAAGGCTCTCGCTGGGGTTCCTGTGGCACACATTGAGGGAACCAAGTACCACCTGAAGAGTGGCCACGTGACCTGCGAAGTGGGACTGGAAAAACTGAAGATGAAAGGTCTTACGTACACAATGTGTGACAAAACAAAGTTCACATGGAAGAGAGCTCCAACAGACAGTGGGCATGATACAGTGGTCATGGAAGTCACATTCTCTGGAACAAAGCCCTGTAGGATCCCAGTCAGGGCAGTGGCACATGGATCTCCAGATGTGAACGTGGCCATGCTGATAACGCCAAACCCAACAATTGAAAACAATGGAGGTGGCTTCATAGAGATGCAGCTGCCCCCAGGGGATAACATCATCTATGTTGGGGAACTGAGTCATCAATGGTTCCAAAAAGGGAGCAGCATCGGAAGGGTTTTCCAAAAGACCAAGAAAGGCATAGAAAGATTGACAGTGATAGGAGAGCACGCCTGGGACTTCGGTTCTGCTGGAGGCTTTCTGAGTTCAATTGGGAAGGCGGTGCACACGGTCCTTGGTGGTGCTTTCAACAGCATCTTCGGGGGAGTGGGGTTTCTACCAAAGCTTCTATTAGGAGTGGCATTGGCTTGGTTGGGCCTGAACATGAGAAACCCTACAATGTCCATGAGCTTTCTCTTGGCTGGAGGTCTGGTCTTGGCCATGACCCTTGGAGTGGGGGCTGATGTTGGCTGCGCTGTGGACACGGAACGAATGGAGCTCCGCTGTGGCGAGGGCCTGGTCGTGTGGAGAGAGGTCTCAGAATGGTATGACAACTATGCCTACTACCCGGAGACACCGGGGGCCCTTGCATCAGCCATAAAGGAGACATTTGAGGAGGGAAGTTGTGGCGTAGTCCCCCAGAACAGGCTCGAGATGGCCATGTGGAGAAACTCGGTCACAGAGCTGAATCTGGCTCTGGCGGAAGGAGAGGCAAATCTCACAGTGGTGGTGGACAAGTTTGACCCCACTGACTACCGAGGTGGTGTCCCTGGTTTACTGAAAAAAGGAAAGGACATAAAAGTCTCCTGGAAAAGCTGGGGCCATTCAATGATCTGGAGCATTCCTGAGGCCCCCCGTCGCTTCATGGTGGGCACGGAAGGACAAAGTGAGTGTCCCCTAGAGAGACGGAAGACAGGTGTTTTCACGGTGGCAGAATTCGGGGTTGGCCTGAGAACAAAGGTCTTCTTGGATTTCAGACAGGAACCAACACATGAGTGTGACACAGGAGTGATGGGAGCTGCCGTCAAGAACGGCATGGCAGTCCACACAGATCAAAGTCTCTGGATGAGGTCAATGAAAAATGACACAGGCACTTACATAGTTGAACTTTTGGTCACTGACCTGAGGAACTGCTCGTGGCCTGCTAGCCACACTATCGATAATGCTGACGTGGTGGACTCGGAGTTATTCCTTCCGGCGAGCCTGGCAGGACCCAGATCCTGGTACAACAGGATACCTGGTTATTCAGAGCAGGTGAAAGGGCCATGGAAGTACACGCCTATCCGAGTCATCAGAGAGGAGTGTCCCGGCACGACCGTTACCATCAACGCCAAGTGTGACAAAAGAGGAGCATCTGTGAGGAGTACCACAGAGAGCGGCAAGGTTATCCCAGAATGGTGCTGCCGAGCATGCACAATGCCACCAGTGACGTTCCGGACTGGAACTGATTGCTGGTATGCCATGGAAATACGGCCAGTCCATGACCAGGGGGGGCTTGTTCGCTCAATGGTGGTTGCGGACAATGGTGAATTACTTAGTGAGGGAGGGGTCCCCGGAATAGTGGCATTGTTTGTGGTCCTTGAATACATCATCCGTAGGAGACCCTCCACGGGAGCGACGGTTGTGTGGGGGGGCATCGTCGTTCTCGCTCTGCTTGTCACCGGGATGGTCAGGATAGAGAGCCTGGTGCGCTATGTGGTGGCAGTGGGGATCACATTCCACCTTGAGCTAGGGCCAGAGATCGTGGCATTGATGCTACTCCAGGCTGTGTTTGAGCTGAGGGTGGGTTTGCTCAGCGCATTTGCACTGCGTAGAAGCCTCACCGTCCGAGAGATGGTGACCACCTACTTTCTCTTGCTGGTCCTGGAATTGGGGCTGCCGGGTGCGAGCCTTGAGGATTTCTGGAAATGGGGTGATGCACTGGCTATGGGGGCGCTGATATTCAGGGCTTGCACGGCGGAAGGAAAGACTGGAGCGGGGCTCTTGCTCATGGCTCTCATGACACAGCAGGATGTGGTGACTGTGCATCATGGACTGGTGTGCTTCCTGAGTGTAGCTTCGGCTTGCTCGGTCTGGAGGCTGCTCAAGGGACACAGAGAGCAGAAGGGATTGACCTGGATTGTCCCCCTGGCTAGATTGCTTGGGGGAGAGGGCTCTGGAATCAGACTGCTGGCGTTTTGGGAGCTGTCAGCTCGCAGAGGGAGACGATCCTTCAGTGAACCACTAACTGTGGTAGGAGTCATGCTAACACTGGCCAGCGGCATGATGCGACACACCTCCCAGGAGGCTCTCTGTGCACTCGCAGTGGCCTCGTTTCTCTTGTTGATGCTGGTGCTGGGGACAAGAAAGATGCAGCTGGTTGCCGAATGGAGTGGCTGCGTGGAATGGCACCCGGAACTAGTGAATGAGGGTGGAGAGGTTAGCCTGCGGGTCCGTCAGGACGCGATGGGAAACTTTCACTTGACTGAGCTCGAGAAAGAAGAGAGAATGATGGCTTTTTGGCTGCTTGCCGGCTTGGCAGCCTCGGCCATTCATTGGTCAGGCATTCTTGGTGTGATGGGACTGTGGACGCTCACGGAAATGCTGAGGTCATCCCGAAGGTCTGACCTGATTTTCTCTGGACAGGGGGGTCGAGAGCGTGGTGACAGACCTTTCGAGGTTAGGGACGGTGTCTACAGGATTTTCAGCCCCGGCTTGTTCTGGGGTCAGAACCAGGTGGGAGTTGGCTACGGTTCCAAGGGTGTTTTACACACGATGTGGCACGTGACAAGAGGAGCGGCGCTGTCTATTGATGATGCTGTAGCCGGTCCCTACTGGGCTGATGTGAGGGAAGATGTCGTGTGCTACGGAGGAGCCTGGAGTCTGGAGGAAAAATGGAAAGGTGAAACAGTACAGGTCCATGCCTTCCCACCGGGGAAGGCCCATGAGGTGCATCAGTGCCAGCCTGGGGAGTTGATCCTTGACACCGGAAGGAAGCTCGGGGCAATACCAATTGATTTGGTAAAAGGAACATCAGGCAGCCCCATTCTTAACGCCCAGGGAGTGGTTGTGGGGCTATATGGAAATGGCCTAAAAACTAATGAGACCTACGTCAGCAGCATTGCTCAAGGAGAAGCAGAGAGGAGTCGACCCAACCTTCCACGGGCTGTTGTGGGTACTGGCTGGACATCAAAGGGTCAGATCACAGTGCTGGACATGCACCCAGGCTCGGGGAAGACCCACAGAGTCCTCCCGGAGCTCATTCGCCAATGCATTGACAGGCGCCTGAGAACGTTGGTGTTGGCTCCAACTCGTGTGGTACTCAAAGAAATGGAGCGCGCCTTGAATGGGAAACGGGTCAGGTTCCACTCACCAGCAGTCAGTGACCAACAGGCTGGAGGGGCAATTGTCGATGTGATGTGTCACGCAACCTATGTCAACAGAAGGCTACTCCCACAGGGGAGACAAAATTGGGAGGTGGCAATCATGGATGAGGCCCACTGGACGGACCCCCACAGCATAGCTGCCAGAGGTCATTTGTACACTCTGGCAAAAGAAAACAAGTGTGCACTGGTCTTGATGACAGCGACACCTCCTGGCAAGAGTGAACCCTTTCCGGAGTCTAACGGAGCCATTACTAGTGAGGAAAGACAGATTCCTGATGGAGAGTGGCGCGACGGGTTTGACTGGATCACTGAGTATGAAGGGCGCACCGCTTGGTTTGTCCCTTCGATTGCAAAAGGTGGGGCTATAGCTCGCACCTTGAGACAGAAGGGGAAAAGTGTGATCTGTTTGAACAGCAAAACCTTTGAAAAGGACTACTCCAGAGTGAGGGATGAGAAGCCTGACTTTGTGGTGACGACTGATATTTCGGAGATGGGAGCCAACCTTGACGTGAGCCGCGTCATAGATGGGAGGACAAACATCAAGCCTGAAGAGGTTGATGGGAAGGTCGAGCTCACCGGGACCAGGCGAGTGACCACGGCTTCCGCTGCCCAACGGCGCGGGAGAGTTGGTCGGCAAGACGGACGAACAGATGAATACATATACTCTGGACAGTGTGATGATGATGACAGTGGATTAGTGCAATGGAAAGAGGCGCAAATACTTCTTGACAACATAACAACCTTGCGGGGGCCCGTGGCCACCTTCTATGGACCAGAACAAGACAAGATGCCGGAGGTGGCTGGTCACTTTCGACTCACTGAAGAGAAAAGAAAGCACTTCCGACATCTTCTCACCCATTGTGACTTCACACCGTGGCTGGCATGGCACGTCGCAGCAAATGTGTCCAGCGTCACGGATCGAAGCTGGACATGGGAAGGGCCGGAGGCAAATGCCGTGGATGAGGCCAGTGGTGACTTGGTCACCTTTAGGAGCCCGAATGGGGCGGAGAGAACTCTCAGGCCGGTGTGGAAGGACGCACGCATGTTCAAAGAGGGACGTGACATCAAAGAGTTCGTGGCGTACGCGTCTGGGCGTCGCAGCTTTGGAGATGTTCTGACAGGAATGTCGGGAGTTCCTGAGCTTTTGCGGCACAGGTGCGTCAGTGCCCTGGATGTCTTCTACACACTTATGCATGAGGAACCTGGCAGCAGGGCAATGAGAATGGCGGAGAGAGATGCCCCAGAGGCCTTTCTGACTATGGTTGAGATGATGGTACTGGGCTTGGCAACCCTGGGTGTCATCTGGTGCTTCGTCGTCCGGACTTCAATCAGCCGTATGATGCTGGGCACGCTGGTCCTGCTGGCCTCCTTGCTACTTTTGTGGGCAGGTGGCGTCGGCTATGGAAATATGGCCGGAGTGGCTCTCATCTTTTACACGTTGCTGACAGTGCTGCAGCCCGAGGCGGGAAAACAGAGAAGCAGTGACGACAACAAACTGGCATATTTCTTGCTGACGCTCTGCAGCCTTGCTGGACTGGTTGCAGCCAATGAGATGGGTTTTCTGGAGAAGACCAAGGCAGACTTGTCCACGGTGCTGTGGAGTGAACAGGAGGAACCCCGGCCATGGAGTGAATGGACGAATGTGGACATCCAGCCAGCGAGGTCCTGGGGGACCTACGTGCTGGTGGTGTCTCTGTTTACACCGTACATCATCCACCAACTGCAAACCAAAATCCAACAACTTGTCAACAGTGCCGTGGCATCTGGTGCACAGGCCATGAGAGACCTTGGGGGAGGTGCCCCCTTCTTTGGTGTGGCGGGACATGTCATGACCCTCGGGGTGGTGTCACTGATTGGGGCTACTCCCACCTCACTGATGGTGGGCGTTGGCTTGGCGGCACTCCATCTGGCCATTGTGGTGTCTGGCCTGGAGGCTGAATTGACACAGAGAGCTCATAAGGTCTTTTTCTCTGCAATGGTGCGCAACCCCATGGTGGATGGGGATGTCATCAACCCATTCGGGGAGGGGGAGGCAAAACCTGCTCTATATGAAAGGAAAATGAGTCTGGTGTTGGCCATAGTGTTGTGCCTCATGTCGGTGGTCATGAACCGAACGGTGGCTTCCATAACAGAAGCTTCAGCTGTGGGACTGGCAGCAGCGGGACAGCTGCTTAGACCGGAGGCTGACACGCTGTGGACGATGCCGGTTGCTTGTGGCATGAGTGGTGTGGTCAGGGGTAGCCTGTGGGGGTTTCTGCCTCTTGGGCATAGACTCTGGCTTCGAGCTTCTGGGGGCAGACGTGGCGGTTCTGAGGGAGACACGCTTGGAGATCTCTGGAAACGGAGGCTGAACAACTGCACCAGGGAGGAATTCTTTGTGTACAGGCGCACTGGCATCCTTGAGACGGAACGTGACAAGGCTAGAGAGTTGCTCAGAAGAGGAGAGACCAATATGGGATTGGCTGTCTCTCGGGGCACGGCAAAGCTTGCCTGGCTTGAGGAACGCGGATATGCCACCCTCAAGGGAGAGGTGGTAGATCTTGGATGTGGAAGGGGCGGCTGGTCCTACTATGCGGCATCCCGACCGGCAGTCATGAGTGTCAGGGCGTACACCATTGGTGGAAGAGGGCACGAGGCTCCAAAGATGGTAACAAGCCTGGGTTGGAACTTGATTAAATTTAGATCAGGAATGGACGTGTTCAGCATGCAGCCACACCGGGCTGACACTGTCATGTGTGACATCGGAGAGAGCAGCCCAGATGCCGCTGTGGAAGGTGAGAGGACAAGGAAAGTGATATTGCTCATGGAGCAATGGAAAAACAGGAATCCCACGGCTGCCTGTGTGTTCAAGGTGCTGGCCCCATACCGCCCAGAAGTGATAGAAGCACTGCACAGATTCCAACTGCAGTGGGGGGGGGGTCTGGTGAGGACCCCTTTTTCAAGGAACTCCACCCATGAGATGTATTACTCAACAGCTGTCACTGGGAACATAGTGAACTCCGTCAATGTACAGTCGAGGAAACTTTTGGCTCGGTTTGGAGACCAGAGAGGGCCAACCAGGGTGCCTGAACTTGACCTGGGAGTTGGAACGAGGTGTGTGGTCTTAGCTGAGGACAAGGTGAAAGAACAAGACGTACAAGAGAGGATCAGAGCGTTGCGTGAGCAATACAGCGAAACCTGGCATATGGACGAGGAACACCCGTACCGGACATGGCAGTATTGGGGCAGCTACCGCACGGCACCAACCGGCTCGGCGGCGTCACTGATCAATGGAGTTGTGAAACTTCTCAGCTGGCCATGGAACGCACGGGAAGATGTGGTGCGCATGGCTATGACTGACACAACGGCTTTTGGACAGCAGAGAGTGTTCAAGGACAAAGTTGACACAAAGGCACAGGAGCCTCAGCCCGGTACAAGAGTCATCATGAGAGCAGTAAATGATTGGATTTTGGAACGACTGGCGCAGAAAAGCAAACCGCGCATGTGCAGCAAAGAAGAATTCATAGCAAAAGTGAAATCAAATGCAGCCTTGGGAGCTTGGTCAGATGAGCAAAACAGATGGGCAAGTGCAAGAGAGGCTGTAGAGGATCCTGCATTTTGGCACCTCGTGGATGAAGAGAGAGAAAGGCACCTCATGGGGAGATGCGCGCACTGCGTGTACAACATGATGGGCAAGAGAGAGAAGAAACTGGGAGAGTTCGGAGTGGCAAAAGGAAGTCGGGCCATTTGGTACATGTGGCTGGGGAGTCGCTTTCTGGAGTTCGAAGCTCTTGGATTCTTGAATGAAGACCATTGGGCCTCTAGAGAGTCCAGTGGAGCTGGAGTTGAAGGAATAAGCTTGAACTACCTGGGCTGGCACCTCAAGAAGTTGTCAACCCTGAATGGAGGACTCTTCTATGCAGATGACACAGCTGGCTGGGACACGAAAGTTACCAATGCAGACTTAGAGGATGAAGAACAGATCCTACGGTACATGGAGGGTGAGCACAAACAATTGGCAACCACAATAATGCAAAAAGCATACCATGCCAAAGTCGTGAAGGTCGCGAGGCCCTCCCGTGATGGAGGCTGCATCATGGATGTCATCACAAGAAGAGACCAAAGAGGCTCGGGCCAGGTTGTGACCTATGCCCTTAACACCCTCACCAACATAAAGGTGCAACTAATCCGAATGATGGAAGGGGAAGGGGTCATAGAGGCAGCGGATGCACACAACCCGAGACTGCTTCGAGTGGAGCGCTGGCTGAAAGAACATGGAGAAGAGCGTCTTGGAAGAATGCTCGTTAGTGGTGACGATTGTGTGGTGAGGCCCTTGGATGACAGATTTGGCAAAGCACTCTACTTTCTGAATGACATGGCCAAGACCAGGAAGGACATTGGGGAATGGGAGCACTCGGCCGGCTTTTCAAGCTGGGAGGAGGTCCCCTTTTGTTCACACCATTTCCACGAGCTAGTGATGAAGGACGGACGCACCCTGGTGGTGCCGTGCCGAGACCAAGATGAACTCGTTGGGAGGGCGCGCATCTCACCGGGGTGCGGCTGGAGTGTCCGCGAGACGGCCTGCCTTTCAAAAGCCTACGGGCAGATGTGGCTGCTGAGCTATTTCCACCGGCGAGACCTGAGGACGCTCGGGCTCGCCATCAACTCAGCAGTGCCTGTCGATTGGGTTCCCACCGGCCGCACGACGTGGAGCATCCATGCCAGTGGGGCCTGGATGACCACAGAAGACATGCTGGACGTTTGGAACCGGGTGTGGATCCTGGACAACCCTTTCATGCAGAACAAGGAAAAGGTCATGGAGTGGAGGGATGTTCCGTACCTCCCTAAAGCTCAGGACATGTTATGTTCCTCCCTTGTTGGGAGGAGAGAAAGAGCAGAATGGGCTAAGAACATCTGGGGAGCGGTGGAAAAGGTGAGGAAGATGATAGGTCCTGAAAAGTTCAAGGACTATCTCTCCTGTATGGACCGCCATGACCTGCACTGGGAGCTCAGACTGGAGAGCTCAATAATCtaaacccagactgtgacagagcaaaacccggagggctcgcaaaagattgtccggaaccaaaacagaagcaagcaactcacagagatagagctcggactggagagctctttaaacaaaaaagccagaattgagctgaacctggagagctcattaaatatagtccagacaaaacaaaacatgacaaagtaaataggctgagctaaaagctcccaccacgggactgcttcatagcggtttgtggggggaggctaggaggcgaagccacagatcatggagtgatgcggcagcgcgcgagagcgacggggaagtggtcgcacccgacgcactatccatgaagcaatacttcgtgagacccccctggccagcaaagggggcagactggtcaggggtaagggatgcccccagagtgcattacggcagcacgccagtgagagtggcgacgggaaaatggtcgatcccgacgtagggcactctgaaaaattttgtgagaccccctgcatcatgacaaggccgaacatggtgcatgaaaaggggaggcccccggaagcacgcttccgggaggagggaagagagaaattggcagctctcttcaggatttttcctcctcctatacaaaattccccctcggtagagggggggcggttcttgttctccctgagccaccatcacccagacacagatagtctgacaaggaggtgatgtgtgactcggaaaaacacccgct |
| circular TBEVnluc | GTGATGCGGTTTTGGCAGTACATCAATGGGCGTGGATAGCGGTTTGACTCACGGGGATTTCCAAGTCTCCACCCCATTGACGTCAATGGGAGTTTGTTTTGGCACCAAAATCAACGGGACTTTCCAAAATGTCGTAACAACTCCGCCCCATTGACGCAAATGGGCGGTAGGCGTGTACGGTGGGAGGTCTATATAAGCAGAGCTGGTTTAGTGAACCGagattttcttgcacgtgcatgcgtttgcttcggacagcaatagcagcggttggtttgaaagagatattcttttgtttctaccagtcgtgaacgtgttgagaaaaagacagcttaggagaacaagagctggggATGGTCAAGAAGGCCATCCTGAAAGGTAAGGGGGGCGGTCCCCCTCGACGAGTGTCGAAAGAGACCGCAACGATGGTCTTCACACTCGAAGATTTCGTTGGGGACTGGCGACAGACAGCCGGCTACAACCTGGACCAAGTCCTTGAACAGGGAGGTGTGTCCAGTTTGTTTCAGAATCTCGGGGTGTCCGTAACTCCGATCCAAAGGATTGTCCTGAGCGGTGAAAATGGGCTGAAGATCGACATCCATGTCATCATCCCGTATGAAGGTCTGAGCGGCGACCAAATGGGCCAGATCGAAAAAATTTTTAAGGTGGTGTACCCTGTGGATGATCATCACTTTAAGGTGATCCTGCACTATGGCACACTGGTAATCGACGGGGTTACGCCGAACATGATCGACTATTTCGGACGGCCGTATGAAGGCATCGCCGTGTTCGACGGCAAAAAGATCACTGTAACAGGGACCCTGTGGAACGGCAACAAAATTATCGACGAGCGCCTGATCAACCCCGACGGCTCCCTGCTGTTCCGAGTAACCATCAACGGAGTGACCGGCTGGCGGCTGTGCGAACGCATTCTGGCGGGAAGCGGAGCTACTAACTTCAGCCTGCTGAAGCAGGCTGGAGACGTGGAGGAGAACCCTGGACCTATGGTGAAAAAAGCTATTCTCAAGGGCAAAGGCGGAGGACCTCCAAGGAGGGTCAGCAAGGAAACAGCCACCAAGACGCGTCAACCCAGAGTCCAAATGCCAAATGGGCTTGTGTTGATGCGCATGATGGGGATCTTGTGGCATGCCGTAGCTGGCACCGCGAGAAACCCCGTATTGAAGGCGTTTTGGAACTCGGTCCCTCTGAAACAGGCCACAGCAGCACTGCGGAAGATCAAAAGGACGGTGAGTGCTCTCATGGTTGGCTTGCAAAAACGTGGGAAAAGGAGGTCAGCGACGGACTGGATGAGCTGGTTGCTGGTCATTACTCTGTTGGGGATGACGCTTGCTGCAACGGTGAGGAAAGAAAGGGACGGCTCAACTGTGATCAGAGCTGAAGGAAAGGATGCAGCAACTCAGGTGCGTGTGGAGAATGGCACCTGTGTGATCCTGGCTACTGACATGGGGTCATGGTGTGATGATTCACTGTCCTATGAGTGTGTGACCATAGATCAAGGAGAGGAGCCTGTTGACGTGGACTGCTTTTGCCGGAACGTTGATGGAGTCTATCTGGAGTATGGACGCTGTGGGAAACAGGAAGGCTCACGGACAAGGCGCTCAGTGCTGATCCCATCCCATGCTCAGGGAGAGCTGACGGGGAGGGGACACAAATGGCTAGAAGGAGACTCGCTGCGAACACACCTCACTAGAGTTGAGGGATGGGTCTGGAAGAACAAGCTACTTGCCTTGGCGATGGTTACCGTTGTGTGGTTGACCCTGGAGAGTGTGGTGACCAGGGTCGCCGTTCTGGTTGTGCTCCTGTGTTTGGCACCGGTCTACGCTTCTCGTTGCACACACTTGGAAAACAGGGACTTTGTGACTGGAACTCAGGGGACTACGAGGGTCACCTTGGTGCTGGAACTGGGTGGATGTGTTACCATAACAGCTGAGGGGAAGCCTTCGATGGATGTGTGGCTTGACGCCATTTACCAGGAGAACCCTGCTAAGACACGTGAGTACTGTTTGCACGCCAAGTTGTCGGACACTAAGGTTGCAGCCAGATGTCCAACAATGGGACCAGCCACTTTGGCTGAAGAACACCAGGGTGGCACAGTGTGCAAGAGAGATCAGAGTGATCGAGGCTGGGGCAACCACTGTGGACTGTTTGGAAAGGGTAGCATTGTGGCCTGTGTCAAGGCGGCTTGTGAGGCAAAAAAGAAAGCCACAGGACATGTGTACGACGCCAACAAAATAGTGTACACGGTTAAAGTCGAACCACACACGGGAGACTATGTTGCCGCAAACGAGACACACAGTGGGAGGAAGACGGCATCCTTCACGGTCTCTTCAGAGAAAACCATTCTAACTATGGGTGAATATGGAGATGTGTCTTTGTTGTGCAGGGTCGCTAGTGGCGTTGACTTGGCCCAGACCGTCATCCTTGAGCTTGACAAGACAGTGGAACACCTTCCAACGGCTTGGCAGGTCCACAGGGACTGGTTCAATGATCTGGCTCTGCCATGGAAACATGAGGGAGCGCAAAACTGGAACAACGCAGAAAGACTGGTTGAATTTGGGGCTCCTCACGCTGTCAAGATGGACGTGTACAACCTCGGAGACCAGACTGGAGTGTTACTGAAGGCTCTCGCTGGGGTTCCTGTGGCACACATTGAGGGAACCAAGTACCACCTGAAGAGTGGCCACGTGACCTGCGAAGTGGGACTGGAAAAACTGAAGATGAAAGGTCTTACGTACACAATGTGTGACAAAACAAAGTTCACATGGAAGAGAGCTCCAACAGACAGTGGGCATGATACAGTGGTCATGGAAGTCACATTCTCTGGAACAAAGCCCTGTAGGATCCCAGTCAGGGCAGTGGCACATGGATCTCCAGATGTGAACGTGGCCATGCTGATAACGCCAAACCCAACAATTGAAAACAATGGAGGTGGCTTCATAGAGATGCAGCTGCCCCCAGGGGATAACATCATCTATGTTGGGGAACTGAGTCATCAATGGTTCCAAAAAGGGAGCAGCATCGGAAGGGTTTTCCAAAAGACCAAGAAAGGCATAGAAAGATTGACAGTGATAGGAGAGCACGCCTGGGACTTCGGTTCTGCTGGAGGCTTTCTGAGTTCAATTGGGAAGGCGGTGCACACGGTCCTTGGTGGTGCTTTCAACAGCATCTTCGGGGGAGTGGGGTTTCTACCAAAGCTTCTATTAGGAGTGGCATTGGCTTGGTTGGGCCTGAACATGAGAAACCCTACAATGTCCATGAGCTTTCTCTTGGCTGGAGGTCTGGTCTTGGCCATGACCCTTGGAGTGGGGGCTGATGTTGGCTGCGCTGTGGACACGGAACGAATGGAGCTCCGCTGTGGCGAGGGCCTGGTCGTGTGGAGAGAGGTCTCAGAATGGTATGACAACTATGCCTACTACCCGGAGACACCGGGGGCCCTTGCATCAGCCATAAAGGAGACATTTGAGGAGGGAAGTTGTGGCGTAGTCCCCCAGAACAGGCTCGAGATGGCCATGTGGAGAAACTCGGTCACAGAGCTGAATCTGGCTCTGGCGGAAGGAGAGGCAAATCTCACAGTGGTGGTGGACAAGTTTGACCCCACTGACTACCGAGGTGGTGTCCCTGGTTTACTGAAAAAAGGAAAGGACATAAAAGTCTCCTGGAAAAGCTGGGGCCATTCAATGATCTGGAGCATTCCTGAGGCCCCCCGTCGCTTCATGGTGGGCACGGAAGGACAAAGTGAGTGTCCCCTAGAGAGACGGAAGACAGGTGTTTTCACGGTGGCAGAATTCGGGGTTGGCCTGAGAACAAAGGTCTTCTTGGATTTCAGACAGGAACCAACACATGAGTGTGACACAGGAGTGATGGGAGCTGCCGTCAAGAACGGCATGGCAGTCCACACAGATCAAAGTCTCTGGATGAGGTCAATGAAAAATGACACAGGCACTTACATAGTTGAACTTTTGGTCACTGACCTGAGGAACTGCTCGTGGCCTGCTAGCCACACTATCGATAATGCTGACGTGGTGGACTCGGAGTTATTCCTTCCGGCGAGCCTGGCAGGACCCAGATCCTGGTACAACAGGATACCTGGTTATTCAGAGCAGGTGAAAGGGCCATGGAAGTACACGCCTATCCGAGTCATCAGAGAGGAGTGTCCCGGCACGACCGTTACCATCAACGCCAAGTGTGACAAAAGAGGAGCATCTGTGAGGAGTACCACAGAGAGCGGCAAGGTTATCCCAGAATGGTGCTGCCGAGCATGCACAATGCCACCAGTGACGTTCCGGACTGGAACTGATTGCTGGTATGCCATGGAAATACGGCCAGTCCATGACCAGGGGGGGCTTGTTCGCTCAATGGTGGTTGCGGACAATGGTGAATTACTTAGTGAGGGAGGGGTCCCCGGAATAGTGGCATTGTTTGTGGTCCTTGAATACATCATCCGTAGGAGACCCTCCACGGGAGCGACGGTTGTGTGGGGGGGCATCGTCGTTCTCGCTCTGCTTGTCACCGGGATGGTCAGGATAGAGAGCCTGGTGCGCTATGTGGTGGCAGTGGGGATCACATTCCACCTTGAGCTAGGGCCAGAGATCGTGGCATTGATGCTACTCCAGGCTGTGTTTGAGCTGAGGGTGGGTTTGCTCAGCGCATTTGCACTGCGTAGAAGCCTCACCGTCCGAGAGATGGTGACCACCTACTTTCTCTTGCTGGTCCTGGAATTGGGGCTGCCGGGTGCGAGCCTTGAGGATTTCTGGAAATGGGGTGATGCACTGGCTATGGGGGCGCTGATATTCAGGGCTTGCACGGCGGAAGGAAAGACTGGAGCGGGGCTCTTGCTCATGGCTCTCATGACACAGCAGGATGTGGTGACTGTGCATCATGGACTGGTGTGCTTCCTGAGTGTAGCTTCGGCTTGCTCGGTCTGGAGGCTGCTCAAGGGACACAGAGAGCAGAAGGGATTGACCTGGATTGTCCCCCTGGCTAGATTGCTTGGGGGAGAGGGCTCTGGAATCAGACTGCTGGCGTTTTGGGAGCTGTCAGCTCGCAGAGGGAGACGATCCTTCAGTGAACCACTAACTGTGGTAGGAGTCATGCTAACACTGGCCAGCGGCATGATGCGACACACCTCCCAGGAGGCTCTCTGTGCACTCGCAGTGGCCTCGTTTCTCTTGTTGATGCTGGTGCTGGGGACAAGAAAGATGCAGCTGGTTGCCGAATGGAGTGGCTGCGTGGAATGGCACCCGGAACTAGTGAATGAGGGTGGAGAGGTTAGCCTGCGGGTCCGTCAGGACGCGATGGGAAACTTTCACTTGACTGAGCTCGAGAAAGAAGAGAGAATGATGGCTTTTTGGCTGCTTGCCGGCTTGGCAGCCTCGGCCATTCATTGGTCAGGCATTCTTGGTGTGATGGGACTGTGGACGCTCACGGAAATGCTGAGGTCATCCCGAAGGTCTGACCTGATTTTCTCTGGACAGGGGGGTCGAGAGCGTGGTGACAGACCTTTCGAGGTTAGGGACGGTGTCTACAGGATTTTCAGCCCCGGCTTGTTCTGGGGTCAGAACCAGGTGGGAGTTGGCTACGGTTCCAAGGGTGTTTTACACACGATGTGGCACGTGACAAGAGGAGCGGCGCTGTCTATTGATGATGCTGTAGCCGGTCCCTACTGGGCTGATGTGAGGGAAGATGTCGTGTGCTACGGAGGAGCCTGGAGTCTGGAGGAAAAATGGAAAGGTGAAACAGTACAGGTCCATGCCTTCCCACCGGGGAAGGCCCATGAGGTGCATCAGTGCCAGCCTGGGGAGTTGATCCTTGACACCGGAAGGAAGCTCGGGGCAATACCAATTGATTTGGTAAAAGGAACATCAGGCAGCCCCATTCTTAACGCCCAGGGAGTGGTTGTGGGGCTATATGGAAATGGCCTAAAAACTAATGAGACCTACGTCAGCAGCATTGCTCAAGGAGAAGCAGAGAGGAGTCGACCCAACCTTCCACGGGCTGTTGTGGGTACTGGCTGGACATCAAAGGGTCAGATCACAGTGCTGGACATGCACCCAGGCTCGGGGAAGACCCACAGAGTCCTCCCGGAGCTCATTCGCCAATGCATTGACAGGCGCCTGAGAACGTTGGTGTTGGCTCCAACTCGTGTGGTACTCAAAGAAATGGAGCGCGCCTTGAATGGGAAACGGGTCAGGTTCCACTCACCAGCAGTCAGTGACCAACAGGCTGGAGGGGCAATTGTCGATGTGATGTGTCACGCAACCTATGTCAACAGAAGGCTACTCCCACAGGGGAGACAAAATTGGGAGGTGGCAATCATGGATGAGGCCCACTGGACGGACCCCCACAGCATAGCTGCCAGAGGTCATTTGTACACTCTGGCAAAAGAAAACAAGTGTGCACTGGTCTTGATGACAGCGACACCTCCTGGCAAGAGTGAACCCTTTCCGGAGTCTAACGGAGCCATTACTAGTGAGGAAAGACAGATTCCTGATGGAGAGTGGCGCGACGGGTTTGACTGGATCACTGAGTATGAAGGGCGCACCGCTTGGTTTGTCCCTTCGATTGCAAAAGGTGGGGCTATAGCTCGCACCTTGAGACAGAAGGGGAAAAGTGTGATCTGTTTGAACAGCAAAACCTTTGAAAAGGACTACTCCAGAGTGAGGGATGAGAAGCCTGACTTTGTGGTGACGACTGATATTTCGGAGATGGGAGCCAACCTTGACGTGAGCCGCGTCATAGATGGGAGGACAAACATCAAGCCTGAAGAGGTTGATGGGAAGGTCGAGCTCACCGGGACCAGGCGAGTGACCACGGCTTCCGCTGCCCAACGGCGCGGGAGAGTTGGTCGGCAAGACGGACGAACAGATGAATACATATACTCTGGACAGTGTGATGATGATGACAGTGGATTAGTGCAATGGAAAGAGGCGCAAATACTTCTTGACAACATAACAACCTTGCGGGGGCCCGTGGCCACCTTCTATGGACCAGAACAAGACAAGATGCCGGAGGTGGCTGGTCACTTTCGACTCACTGAAGAGAAAAGAAAGCACTTCCGACATCTTCTCACCCATTGTGACTTCACACCGTGGCTGGCATGGCACGTCGCAGCAAATGTGTCCAGCGTCACGGATCGAAGCTGGACATGGGAAGGGCCGGAGGCAAATGCCGTGGATGAGGCCAGTGGTGACTTGGTCACCTTTAGGAGCCCGAATGGGGCGGAGAGAACTCTCAGGCCGGTGTGGAAGGACGCACGCATGTTCAAAGAGGGACGTGACATCAAAGAGTTCGTGGCGTACGCGTCTGGGCGTCGCAGCTTTGGAGATGTTCTGACAGGAATGTCGGGAGTTCCTGAGCTTTTGCGGCACAGGTGCGTCAGTGCCCTGGATGTCTTCTACACACTTATGCATGAGGAACCTGGCAGCAGGGCAATGAGAATGGCGGAGAGAGATGCCCCAGAGGCCTTTCTGACTATGGTTGAGATGATGGTACTGGGCTTGGCAACCCTGGGTGTCATCTGGTGCTTCGTCGTCCGGACTTCAATCAGCCGTATGATGCTGGGCACGCTGGTCCTGCTGGCCTCCTTGCTACTTTTGTGGGCAGGTGGCGTCGGCTATGGAAATATGGCCGGAGTGGCTCTCATCTTTTACACGTTGCTGACAGTGCTGCAGCCCGAGGCGGGAAAACAGAGAAGCAGTGACGACAACAAACTGGCATATTTCTTGCTGACGCTCTGCAGCCTTGCTGGACTGGTTGCAGCCAATGAGATGGGTTTTCTGGAGAAGACCAAGGCAGACTTGTCCACGGTGCTGTGGAGTGAACAGGAGGAACCCCGGCCATGGAGTGAATGGACGAATGTGGACATCCAGCCAGCGAGGTCCTGGGGGACCTACGTGCTGGTGGTGTCTCTGTTTACACCGTACATCATCCACCAACTGCAAACCAAAATCCAACAACTTGTCAACAGTGCCGTGGCATCTGGTGCACAGGCCATGAGAGACCTTGGGGGAGGTGCCCCCTTCTTTGGTGTGGCGGGACATGTCATGACCCTCGGGGTGGTGTCACTGATTGGGGCTACTCCCACCTCACTGATGGTGGGCGTTGGCTTGGCGGCACTCCATCTGGCCATTGTGGTGTCTGGCCTGGAGGCTGAATTGACACAGAGAGCTCATAAGGTCTTTTTCTCTGCAATGGTGCGCAACCCCATGGTGGATGGGGATGTCATCAACCCATTCGGGGAGGGGGAGGCAAAACCTGCTCTATATGAAAGGAAAATGAGTCTGGTGTTGGCCATAGTGTTGTGCCTCATGTCGGTGGTCATGAACCGAACGGTGGCTTCCATAACAGAAGCTTCAGCTGTGGGACTGGCAGCAGCGGGACAGCTGCTTAGACCGGAGGCTGACACGCTGTGGACGATGCCGGTTGCTTGTGGCATGAGTGGTGTGGTCAGGGGTAGCCTGTGGGGGTTTCTGCCTCTTGGGCATAGACTCTGGCTTCGAGCTTCTGGGGGCAGACGTGGCGGTTCTGAGGGAGACACGCTTGGAGATCTCTGGAAACGGAGGCTGAACAACTGCACCAGGGAGGAATTCTTTGTGTACAGGCGCACTGGCATCCTTGAGACGGAACGTGACAAGGCTAGAGAGTTGCTCAGAAGAGGAGAGACCAATATGGGATTGGCTGTCTCTCGGGGCACGGCAAAGCTTGCCTGGCTTGAGGAACGCGGATATGCCACCCTCAAGGGAGAGGTGGTAGATCTTGGATGTGGAAGGGGCGGCTGGTCCTACTATGCGGCATCCCGACCGGCAGTCATGAGTGTCAGGGCGTACACCATTGGTGGAAGAGGGCACGAGGCTCCAAAGATGGTAACAAGCCTGGGTTGGAACTTGATTAAATTTAGATCAGGAATGGACGTGTTCAGCATGCAGCCACACCGGGCTGACACTGTCATGTGTGACATCGGAGAGAGCAGCCCAGATGCCGCTGTGGAAGGTGAGAGGACAAGGAAAGTGATATTGCTCATGGAGCAATGGAAAAACAGGAATCCCACGGCTGCCTGTGTGTTCAAGGTGCTGGCCCCATACCGCCCAGAAGTGATAGAAGCACTGCACAGATTCCAACTGCAGTGGGGGGGGGGTCTGGTGAGGACCCCTTTTTCAAGGAACTCCACCCATGAGATGTATTACTCAACAGCTGTCACTGGGAACATAGTGAACTCCGTCAATGTACAGTCGAGGAAACTTTTGGCTCGGTTTGGAGACCAGAGAGGGCCAACCAGGGTGCCTGAACTTGACCTGGGAGTTGGAACGAGGTGTGTGGTCTTAGCTGAGGACAAGGTGAAAGAACAAGACGTACAAGAGAGGATCAGAGCGTTGCGTGAGCAATACAGCGAAACCTGGCATATGGACGAGGAACACCCGTACCGGACATGGCAGTATTGGGGCAGCTACCGCACGGCACCAACCGGCTCGGCGGCGTCACTGATCAATGGAGTTGTGAAACTTCTCAGCTGGCCATGGAACGCACGGGAAGATGTGGTGCGCATGGCTATGACTGACACAACGGCTTTTGGACAGCAGAGAGTGTTCAAGGACAAAGTTGACACAAAGGCACAGGAGCCTCAGCCCGGTACAAGAGTCATCATGAGAGCAGTAAATGATTGGATTTTGGAACGACTGGCGCAGAAAAGCAAACCGCGCATGTGCAGCAAAGAAGAATTCATAGCAAAAGTGAAATCAAATGCAGCCTTGGGAGCTTGGTCAGATGAGCAAAACAGATGGGCAAGTGCAAGAGAGGCTGTAGAGGATCCTGCATTTTGGCACCTCGTGGATGAAGAGAGAGAAAGGCACCTCATGGGGAGATGCGCGCACTGCGTGTACAACATGATGGGCAAGAGAGAGAAGAAACTGGGAGAGTTCGGAGTGGCAAAAGGAAGTCGGGCCATTTGGTACATGTGGCTGGGGAGTCGCTTTCTGGAGTTCGAAGCTCTTGGATTCTTGAATGAAGACCATTGGGCCTCTAGAGAGTCCAGTGGAGCTGGAGTTGAAGGAATAAGCTTGAACTACCTGGGCTGGCACCTCAAGAAGTTGTCAACCCTGAATGGAGGACTCTTCTATGCAGATGACACAGCTGGCTGGGACACGAAAGTTACCAATGCAGACTTAGAGGATGAAGAACAGATCCTACGGTACATGGAGGGTGAGCACAAACAATTGGCAACCACAATAATGCAAAAAGCATACCATGCCAAAGTCGTGAAGGTCGCGAGGCCCTCCCGTGATGGAGGCTGCATCATGGATGTCATCACAAGAAGAGACCAAAGAGGCTCGGGCCAGGTTGTGACCTATGCCCTTAACACCCTCACCAACATAAAGGTGCAACTAATCCGAATGATGGAAGGGGAAGGGGTCATAGAGGCAGCGGATGCACACAACCCGAGACTGCTTCGAGTGGAGCGCTGGCTGAAAGAACATGGAGAAGAGCGTCTTGGAAGAATGCTCGTTAGTGGTGACGATTGTGTGGTGAGGCCCTTGGATGACAGATTTGGCAAAGCACTCTACTTTCTGAATGACATGGCCAAGACCAGGAAGGACATTGGGGAATGGGAGCACTCGGCCGGCTTTTCAAGCTGGGAGGAGGTCCCCTTTTGTTCACACCATTTCCACGAGCTAGTGATGAAGGACGGACGCACCCTGGTGGTGCCGTGCCGAGACCAAGATGAACTCGTTGGGAGGGCGCGCATCTCACCGGGGTGCGGCTGGAGTGTCCGCGAGACGGCCTGCCTTTCAAAAGCCTACGGGCAGATGTGGCTGCTGAGCTATTTCCACCGGCGAGACCTGAGGACGCTCGGGCTCGCCATCAACTCAGCAGTGCCTGTCGATTGGGTTCCCACCGGCCGCACGACGTGGAGCATCCATGCCAGTGGGGCCTGGATGACCACAGAAGACATGCTGGACGTTTGGAACCGGGTGTGGATCCTGGACAACCCTTTCATGCAGAACAAGGAAAAGGTCATGGAGTGGAGGGATGTTCCGTACCTCCCTAAAGCTCAGGACATGTTATGTTCCTCCCTTGTTGGGAGGAGAGAAAGAGCAGAATGGGCTAAGAACATCTGGGGAGCGGTGGAAAAGGTGAGGAAGATGATAGGTCCTGAAAAGTTCAAGGACTATCTCTCCTGTATGGACCGCCATGACCTGCACTGGGAGCTCAGACTGGAGAGCTCAATAATCtaaacccagactgtgacagagcaaaacccggagggctcgcaaaagattgtccggaaccaaaacagaagcaagcaactcacagagatagagctcggactggagagctctttaaacaaaaaagccagaattgagctgaacctggagagctcattaaatatagtccagacaaaacaaaacatgacaaagtaaataggctgagctaaaagctcccaccacgggactgcttcatagcggtttgtggggggaggctaggaggcgaagccacagatcatggagtgatgcggcagcgcgcgagagcgacggggaagtggtcgcacccgacgcactatccatgaagcaatacttcgtgagacccccctggccagcaaagggggcagactggtcaggggtaagggatgcccccagagtgcattacggcagcacgccagtgagagtggcgacgggaaaatggtcgatcccgacgtagggcactctgaaaaattttgtgagaccccctgcatcatgacaaggccgaacatggtgcatgaaaaggggaggcccccggaagcacgcttccgggaggagggaagagagaaattggcagctctcttcaggatttttcctcctcctatacaaaattccccctcggtagagggggggcggttcttgttctccctgagccaccatcacccagacacagatagtctgacaaggaggtgatgtgtgactcggaaaaacacccgctGGGTCGGCATGGCATCTCCACCTCCTCGCGGTCCGACCTGGGCATCCGAAGGAGGACGTCGTCCACTCGGATGGCTAAGGGAGAGCTCggatccaccggatctagataactgatcataatcagccataccacatttgtagaggttttacttgctttaaaaaacctcccacacctccccctgaacctgaaacataaaatgaatgcaattgttgttgttAACTTGTTTATTGCAGCTTATAATGGTTACAAATAAAGCAATAGCATCACAAATTTCACAAATAAAGCATTTTTTTCACTGCATTCTAGTTGTGGTTTGTCCAAACTCATCAATGTATCTTAtagtaatcaattacggggtcattagttcatagcccatatatggagttccgCGTTACATAACTTACGGTAAATGGCCCGCCTGGCTGACCGCCCAACGACCCCCGCCCATTGACGTCAATAATGACGTATGTTCCCATAGTAACGCCAATAGGGACTTTCCATTGACGTCAATGGGTGGAGTATTTACGGTAAACTGCCCACTTGGCAGTACATCAAGTGTATCATATGCCAAGTACGCCCCCTATTGACGTCAATGACGGTAAATGGCCCGCCTGGCATTATGCCCAGTACATGACCTTATGGGACTTTCCTACTTGGCAGTACATCTACGTATTAGTCATCGCTATTACCATG |
| infectiousTBEVnluc | agattttcttgcacgtgcatgcgtttgcttcggacagcaatagcagcggttggtttgaaagagatattcttttgtttctaccagtcgtgaacgtgttgagaaaaagacagcttaggagaacaagagctggggATGGTCAAGAAGGCCATCCTGAAAGGTAAGGGGGGCGGTCCCCCTCGACGAGTGTCGAAAGAGACCGCAACGATGGTCTTCACACTCGAAGATTTCGTTGGGGACTGGCGACAGACAGCCGGCTACAACCTGGACCAAGTCCTTGAACAGGGAGGTGTGTCCAGTTTGTTTCAGAATCTCGGGGTGTCCGTAACTCCGATCCAAAGGATTGTCCTGAGCGGTGAAAATGGGCTGAAGATCGACATCCATGTCATCATCCCGTATGAAGGTCTGAGCGGCGACCAAATGGGCCAGATCGAAAAAATTTTTAAGGTGGTGTACCCTGTGGATGATCATCACTTTAAGGTGATCCTGCACTATGGCACACTGGTAATCGACGGGGTTACGCCGAACATGATCGACTATTTCGGACGGCCGTATGAAGGCATCGCCGTGTTCGACGGCAAAAAGATCACTGTAACAGGGACCCTGTGGAACGGCAACAAAATTATCGACGAGCGCCTGATCAACCCCGACGGCTCCCTGCTGTTCCGAGTAACCATCAACGGAGTGACCGGCTGGCGGCTGTGCGAACGCATTCTGGCGGGAAGCGGAGCTACTAACTTCAGCCTGCTGAAGCAGGCTGGAGACGTGGAGGAGAACCCTGGACCTATGGTGAAAAAAGCTATTCTCAAGGGCAAAGGCGGAGGACCTCCAAGGAGGGTCAGCAAGGAAACAGCCACCAAGACGCGTCAACCCAGAGTCCAAATGCCAAATGGGCTTGTGTTGATGCGCATGATGGGGATCTTGTGGCATGCCGTAGCTGGCACCGCGAGAAACCCCGTATTGAAGGCGTTTTGGAACTCGGTCCCTCTGAAACAGGCCACAGCAGCACTGCGGAAGATCAAAAGGACGGTGAGTGCTCTCATGGTTGGCTTGCAAAAACGTGGGAAAAGGAGGTCAGCGACGGACTGGATGAGCTGGTTGCTGGTCATTACTCTGTTGGGGATGACGCTTGCTGCAACGGTGAGGAAAGAAAGGGACGGCTCAACTGTGATCAGAGCTGAAGGAAAGGATGCAGCAACTCAGGTGCGTGTGGAGAATGGCACCTGTGTGATCCTGGCTACTGACATGGGGTCATGGTGTGATGATTCACTGTCCTATGAGTGTGTGACCATAGATCAAGGAGAGGAGCCTGTTGACGTGGACTGCTTTTGCCGGAACGTTGATGGAGTCTATCTGGAGTATGGACGCTGTGGGAAACAGGAAGGCTCACGGACAAGGCGCTCAGTGCTGATCCCATCCCATGCTCAGGGAGAGCTGACGGGGAGGGGACACAAATGGCTAGAAGGAGACTCGCTGCGAACACACCTCACTAGAGTTGAGGGATGGGTCTGGAAGAACAAGCTACTTGCCTTGGCGATGGTTACCGTTGTGTGGTTGACCCTGGAGAGTGTGGTGACCAGGGTCGCCGTTCTGGTTGTGCTCCTGTGTTTGGCACCGGTCTACGCTTCTCGTTGCACACACTTGGAAAACAGGGACTTTGTGACTGGAACTCAGGGGACTACGAGGGTCACCTTGGTGCTGGAACTGGGTGGATGTGTTACCATAACAGCTGAGGGGAAGCCTTCGATGGATGTGTGGCTTGACGCCATTTACCAGGAGAACCCTGCTAAGACACGTGAGTACTGTTTGCACGCCAAGTTGTCGGACACTAAGGTTGCAGCCAGATGTCCAACAATGGGACCAGCCACTTTGGCTGAAGAACACCAGGGTGGCACAGTGTGCAAGAGAGATCAGAGTGATCGAGGCTGGGGCAACCACTGTGGACTGTTTGGAAAGGGTAGCATTGTGGCCTGTGTCAAGGCGGCTTGTGAGGCAAAAAAGAAAGCCACAGGACATGTGTACGACGCCAACAAAATAGTGTACACGGTTAAAGTCGAACCACACACGGGAGACTATGTTGCCGCAAACGAGACACACAGTGGGAGGAAGACGGCATCCTTCACGGTCTCTTCAGAGAAAACCATTCTAACTATGGGTGAATATGGAGATGTGTCTTTGTTGTGCAGGGTCGCTAGTGGCGTTGACTTGGCCCAGACCGTCATCCTTGAGCTTGACAAGACAGTGGAACACCTTCCAACGGCTTGGCAGGTCCACAGGGACTGGTTCAATGATCTGGCTCTGCCATGGAAACATGAGGGAGCGCAAAACTGGAACAACGCAGAAAGACTGGTTGAATTTGGGGCTCCTCACGCTGTCAAGATGGACGTGTACAACCTCGGAGACCAGACTGGAGTGTTACTGAAGGCTCTCGCTGGGGTTCCTGTGGCACACATTGAGGGAACCAAGTACCACCTGAAGAGTGGCCACGTGACCTGCGAAGTGGGACTGGAAAAACTGAAGATGAAAGGTCTTACGTACACAATGTGTGACAAAACAAAGTTCACATGGAAGAGAGCTCCAACAGACAGTGGGCATGATACAGTGGTCATGGAAGTCACATTCTCTGGAACAAAGCCCTGTAGGATCCCAGTCAGGGCAGTGGCACATGGATCTCCAGATGTGAACGTGGCCATGCTGATAACGCCAAACCCAACAATTGAAAACAATGGAGGTGGCTTCATAGAGATGCAGCTGCCCCCAGGGGATAACATCATCTATGTTGGGGAACTGAGTCATCAATGGTTCCAAAAAGGGAGCAGCATCGGAAGGGTTTTCCAAAAGACCAAGAAAGGCATAGAAAGATTGACAGTGATAGGAGAGCACGCCTGGGACTTCGGTTCTGCTGGAGGCTTTCTGAGTTCAATTGGGAAGGCGGTGCACACGGTCCTTGGTGGTGCTTTCAACAGCATCTTCGGGGGAGTGGGGTTTCTACCAAAGCTTCTATTAGGAGTGGCATTGGCTTGGTTGGGCCTGAACATGAGAAACCCTACAATGTCCATGAGCTTTCTCTTGGCTGGAGGTCTGGTCTTGGCCATGACCCTTGGAGTGGGGGCTGATGTTGGCTGCGCTGTGGACACGGAACGAATGGAGCTCCGCTGTGGCGAGGGCCTGGTCGTGTGGAGAGAGGTCTCAGAATGGTATGACAACTATGCCTACTACCCGGAGACACCGGGGGCCCTTGCATCAGCCATAAAGGAGACATTTGAGGAGGGAAGTTGTGGCGTAGTCCCCCAGAACAGGCTCGAGATGGCCATGTGGAGAAACTCGGTCACAGAGCTGAATCTGGCTCTGGCGGAAGGAGAGGCAAATCTCACAGTGGTGGTGGACAAGTTTGACCCCACTGACTACCGAGGTGGTGTCCCTGGTTTACTGAAAAAAGGAAAGGACATAAAAGTCTCCTGGAAAAGCTGGGGCCATTCAATGATCTGGAGCATTCCTGAGGCCCCCCGTCGCTTCATGGTGGGCACGGAAGGACAAAGTGAGTGTCCCCTAGAGAGACGGAAGACAGGTGTTTTCACGGTGGCAGAATTCGGGGTTGGCCTGAGAACAAAGGTCTTCTTGGATTTCAGACAGGAACCAACACATGAGTGTGACACAGGAGTGATGGGAGCTGCCGTCAAGAACGGCATGGCAGTCCACACAGATCAAAGTCTCTGGATGAGGTCAATGAAAAATGACACAGGCACTTACATAGTTGAACTTTTGGTCACTGACCTGAGGAACTGCTCGTGGCCTGCTAGCCACACTATCGATAATGCTGACGTGGTGGACTCGGAGTTATTCCTTCCGGCGAGCCTGGCAGGACCCAGATCCTGGTACAACAGGATACCTGGTTATTCAGAGCAGGTGAAAGGGCCATGGAAGTACACGCCTATCCGAGTCATCAGAGAGGAGTGTCCCGGCACGACCGTTACCATCAACGCCAAGTGTGACAAAAGAGGAGCATCTGTGAGGAGTACCACAGAGAGCGGCAAGGTTATCCCAGAATGGTGCTGCCGAGCATGCACAATGCCACCAGTGACGTTCCGGACTGGAACTGATTGCTGGTATGCCATGGAAATACGGCCAGTCCATGACCAGGGGGGGCTTGTTCGCTCAATGGTGGTTGCGGACAATGGTGAATTACTTAGTGAGGGAGGGGTCCCCGGAATAGTGGCATTGTTTGTGGTCCTTGAATACATCATCCGTAGGAGACCCTCCACGGGAGCGACGGTTGTGTGGGGGGGCATCGTCGTTCTCGCTCTGCTTGTCACCGGGATGGTCAGGATAGAGAGCCTGGTGCGCTATGTGGTGGCAGTGGGGATCACATTCCACCTTGAGCTAGGGCCAGAGATCGTGGCATTGATGCTACTCCAGGCTGTGTTTGAGCTGAGGGTGGGTTTGCTCAGCGCATTTGCACTGCGTAGAAGCCTCACCGTCCGAGAGATGGTGACCACCTACTTTCTCTTGCTGGTCCTGGAATTGGGGCTGCCGGGTGCGAGCCTTGAGGATTTCTGGAAATGGGGTGATGCACTGGCTATGGGGGCGCTGATATTCAGGGCTTGCACGGCGGAAGGAAAGACTGGAGCGGGGCTCTTGCTCATGGCTCTCATGACACAGCAGGATGTGGTGACTGTGCATCATGGACTGGTGTGCTTCCTGAGTGTAGCTTCGGCTTGCTCGGTCTGGAGGCTGCTCAAGGGACACAGAGAGCAGAAGGGATTGACCTGGATTGTCCCCCTGGCTAGATTGCTTGGGGGAGAGGGCTCTGGAATCAGACTGCTGGCGTTTTGGGAGCTGTCAGCTCGCAGAGGGAGACGATCCTTCAGTGAACCACTAACTGTGGTAGGAGTCATGCTAACACTGGCCAGCGGCATGATGCGACACACCTCCCAGGAGGCTCTCTGTGCACTCGCAGTGGCCTCGTTTCTCTTGTTGATGCTGGTGCTGGGGACAAGAAAGATGCAGCTGGTTGCCGAATGGAGTGGCTGCGTGGAATGGCACCCGGAACTAGTGAATGAGGGTGGAGAGGTTAGCCTGCGGGTCCGTCAGGACGCGATGGGAAACTTTCACTTGACTGAGCTCGAGAAAGAAGAGAGAATGATGGCTTTTTGGCTGCTTGCCGGCTTGGCAGCCTCGGCCATTCATTGGTCAGGCATTCTTGGTGTGATGGGACTGTGGACGCTCACGGAAATGCTGAGGTCATCCCGAAGGTCTGACCTGATTTTCTCTGGACAGGGGGGTCGAGAGCGTGGTGACAGACCTTTCGAGGTTAGGGACGGTGTCTACAGGATTTTCAGCCCCGGCTTGTTCTGGGGTCAGAACCAGGTGGGAGTTGGCTACGGTTCCAAGGGTGTTTTACACACGATGTGGCACGTGACAAGAGGAGCGGCGCTGTCTATTGATGATGCTGTAGCCGGTCCCTACTGGGCTGATGTGAGGGAAGATGTCGTGTGCTACGGAGGAGCCTGGAGTCTGGAGGAAAAATGGAAAGGTGAAACAGTACAGGTCCATGCCTTCCCACCGGGGAAGGCCCATGAGGTGCATCAGTGCCAGCCTGGGGAGTTGATCCTTGACACCGGAAGGAAGCTCGGGGCAATACCAATTGATTTGGTAAAAGGAACATCAGGCAGCCCCATTCTTAACGCCCAGGGAGTGGTTGTGGGGCTATATGGAAATGGCCTAAAAACTAATGAGACCTACGTCAGCAGCATTGCTCAAGGAGAAGCAGAGAGGAGTCGACCCAACCTTCCACGGGCTGTTGTGGGTACTGGCTGGACATCAAAGGGTCAGATCACAGTGCTGGACATGCACCCAGGCTCGGGGAAGACCCACAGAGTCCTCCCGGAGCTCATTCGCCAATGCATTGACAGGCGCCTGAGAACGTTGGTGTTGGCTCCAACTCGTGTGGTACTCAAAGAAATGGAGCGCGCCTTGAATGGGAAACGGGTCAGGTTCCACTCACCAGCAGTCAGTGACCAACAGGCTGGAGGGGCAATTGTCGATGTGATGTGTCACGCAACCTATGTCAACAGAAGGCTACTCCCACAGGGGAGACAAAATTGGGAGGTGGCAATCATGGATGAGGCCCACTGGACGGACCCCCACAGCATAGCTGCCAGAGGTCATTTGTACACTCTGGCAAAAGAAAACAAGTGTGCACTGGTCTTGATGACAGCGACACCTCCTGGCAAGAGTGAACCCTTTCCGGAGTCTAACGGAGCCATTACTAGTGAGGAAAGACAGATTCCTGATGGAGAGTGGCGCGACGGGTTTGACTGGATCACTGAGTATGAAGGGCGCACCGCTTGGTTTGTCCCTTCGATTGCAAAAGGTGGGGCTATAGCTCGCACCTTGAGACAGAAGGGGAAAAGTGTGATCTGTTTGAACAGCAAAACCTTTGAAAAGGACTACTCCAGAGTGAGGGATGAGAAGCCTGACTTTGTGGTGACGACTGATATTTCGGAGATGGGAGCCAACCTTGACGTGAGCCGCGTCATAGATGGGAGGACAAACATCAAGCCTGAAGAGGTTGATGGGAAGGTCGAGCTCACCGGGACCAGGCGAGTGACCACGGCTTCCGCTGCCCAACGGCGCGGGAGAGTTGGTCGGCAAGACGGACGAACAGATGAATACATATACTCTGGACAGTGTGATGATGATGACAGTGGATTAGTGCAATGGAAAGAGGCGCAAATACTTCTTGACAACATAACAACCTTGCGGGGGCCCGTGGCCACCTTCTATGGACCAGAACAAGACAAGATGCCGGAGGTGGCTGGTCACTTTCGACTCACTGAAGAGAAAAGAAAGCACTTCCGACATCTTCTCACCCATTGTGACTTCACACCGTGGCTGGCATGGCACGTCGCAGCAAATGTGTCCAGCGTCACGGATCGAAGCTGGACATGGGAAGGGCCGGAGGCAAATGCCGTGGATGAGGCCAGTGGTGACTTGGTCACCTTTAGGAGCCCGAATGGGGCGGAGAGAACTCTCAGGCCGGTGTGGAAGGACGCACGCATGTTCAAAGAGGGACGTGACATCAAAGAGTTCGTGGCGTACGCGTCTGGGCGTCGCAGCTTTGGAGATGTTCTGACAGGAATGTCGGGAGTTCCTGAGCTTTTGCGGCACAGGTGCGTCAGTGCCCTGGATGTCTTCTACACACTTATGCATGAGGAACCTGGCAGCAGGGCAATGAGAATGGCGGAGAGAGATGCCCCAGAGGCCTTTCTGACTATGGTTGAGATGATGGTACTGGGCTTGGCAACCCTGGGTGTCATCTGGTGCTTCGTCGTCCGGACTTCAATCAGCCGTATGATGCTGGGCACGCTGGTCCTGCTGGCCTCCTTGCTACTTTTGTGGGCAGGTGGCGTCGGCTATGGAAATATGGCCGGAGTGGCTCTCATCTTTTACACGTTGCTGACAGTGCTGCAGCCCGAGGCGGGAAAACAGAGAAGCAGTGACGACAACAAACTGGCATATTTCTTGCTGACGCTCTGCAGCCTTGCTGGACTGGTTGCAGCCAATGAGATGGGTTTTCTGGAGAAGACCAAGGCAGACTTGTCCACGGTGCTGTGGAGTGAACAGGAGGAACCCCGGCCATGGAGTGAATGGACGAATGTGGACATCCAGCCAGCGAGGTCCTGGGGGACCTACGTGCTGGTGGTGTCTCTGTTTACACCGTACATCATCCACCAACTGCAAACCAAAATCCAACAACTTGTCAACAGTGCCGTGGCATCTGGTGCACAGGCCATGAGAGACCTTGGGGGAGGTGCCCCCTTCTTTGGTGTGGCGGGACATGTCATGACCCTCGGGGTGGTGTCACTGATTGGGGCTACTCCCACCTCACTGATGGTGGGCGTTGGCTTGGCGGCACTCCATCTGGCCATTGTGGTGTCTGGCCTGGAGGCTGAATTGACACAGAGAGCTCATAAGGTCTTTTTCTCTGCAATGGTGCGCAACCCCATGGTGGATGGGGATGTCATCAACCCATTCGGGGAGGGGGAGGCAAAACCTGCTCTATATGAAAGGAAAATGAGTCTGGTGTTGGCCATAGTGTTGTGCCTCATGTCGGTGGTCATGAACCGAACGGTGGCTTCCATAACAGAAGCTTCAGCTGTGGGACTGGCAGCAGCGGGACAGCTGCTTAGACCGGAGGCTGACACGCTGTGGACGATGCCGGTTGCTTGTGGCATGAGTGGTGTGGTCAGGGGTAGCCTGTGGGGGTTTCTGCCTCTTGGGCATAGACTCTGGCTTCGAGCTTCTGGGGGCAGACGTGGCGGTTCTGAGGGAGACACGCTTGGAGATCTCTGGAAACGGAGGCTGAACAACTGCACCAGGGAGGAATTCTTTGTGTACAGGCGCACTGGCATCCTTGAGACGGAACGTGACAAGGCTAGAGAGTTGCTCAGAAGAGGAGAGACCAATATGGGATTGGCTGTCTCTCGGGGCACGGCAAAGCTTGCCTGGCTTGAGGAACGCGGATATGCCACCCTCAAGGGAGAGGTGGTAGATCTTGGATGTGGAAGGGGCGGCTGGTCCTACTATGCGGCATCCCGACCGGCAGTCATGAGTGTCAGGGCGTACACCATTGGTGGAAGAGGGCACGAGGCTCCAAAGATGGTAACAAGCCTGGGTTGGAACTTGATTAAATTTAGATCAGGAATGGACGTGTTCAGCATGCAGCCACACCGGGCTGACACTGTCATGTGTGACATCGGAGAGAGCAGCCCAGATGCCGCTGTGGAAGGTGAGAGGACAAGGAAAGTGATATTGCTCATGGAGCAATGGAAAAACAGGAATCCCACGGCTGCCTGTGTGTTCAAGGTGCTGGCCCCATACCGCCCAGAAGTGATAGAAGCACTGCACAGATTCCAACTGCAGTGGGGGGGGGGTCTGGTGAGGACCCCTTTTTCAAGGAACTCCACCCATGAGATGTATTACTCAACAGCTGTCACTGGGAACATAGTGAACTCCGTCAATGTACAGTCGAGGAAACTTTTGGCTCGGTTTGGAGACCAGAGAGGGCCAACCAGGGTGCCTGAACTTGACCTGGGAGTTGGAACGAGGTGTGTGGTCTTAGCTGAGGACAAGGTGAAAGAACAAGACGTACAAGAGAGGATCAGAGCGTTGCGTGAGCAATACAGCGAAACCTGGCATATGGACGAGGAACACCCGTACCGGACATGGCAGTATTGGGGCAGCTACCGCACGGCACCAACCGGCTCGGCGGCGTCACTGATCAATGGAGTTGTGAAACTTCTCAGCTGGCCATGGAACGCACGGGAAGATGTGGTGCGCATGGCTATGACTGACACAACGGCTTTTGGACAGCAGAGAGTGTTCAAGGACAAAGTTGACACAAAGGCACAGGAGCCTCAGCCCGGTACAAGAGTCATCATGAGAGCAGTAAATGATTGGATTTTGGAACGACTGGCGCAGAAAAGCAAACCGCGCATGTGCAGCAAAGAAGAATTCATAGCAAAAGTGAAATCAAATGCAGCCTTGGGAGCTTGGTCAGATGAGCAAAACAGATGGGCAAGTGCAAGAGAGGCTGTAGAGGATCCTGCATTTTGGCACCTCGTGGATGAAGAGAGAGAAAGGCACCTCATGGGGAGATGCGCGCACTGCGTGTACAACATGATGGGCAAGAGAGAGAAGAAACTGGGAGAGTTCGGAGTGGCAAAAGGAAGTCGGGCCATTTGGTACATGTGGCTGGGGAGTCGCTTTCTGGAGTTCGAAGCTCTTGGATTCTTGAATGAAGACCATTGGGCCTCTAGAGAGTCCAGTGGAGCTGGAGTTGAAGGAATAAGCTTGAACTACCTGGGCTGGCACCTCAAGAAGTTGTCAACCCTGAATGGAGGACTCTTCTATGCAGATGACACAGCTGGCTGGGACACGAAAGTTACCAATGCAGACTTAGAGGATGAAGAACAGATCCTACGGTACATGGAGGGTGAGCACAAACAATTGGCAACCACAATAATGCAAAAAGCATACCATGCCAAAGTCGTGAAGGTCGCGAGGCCCTCCCGTGATGGAGGCTGCATCATGGATGTCATCACAAGAAGAGACCAAAGAGGCTCGGGCCAGGTTGTGACCTATGCCCTTAACACCCTCACCAACATAAAGGTGCAACTAATCCGAATGATGGAAGGGGAAGGGGTCATAGAGGCAGCGGATGCACACAACCCGAGACTGCTTCGAGTGGAGCGCTGGCTGAAAGAACATGGAGAAGAGCGTCTTGGAAGAATGCTCGTTAGTGGTGACGATTGTGTGGTGAGGCCCTTGGATGACAGATTTGGCAAAGCACTCTACTTTCTGAATGACATGGCCAAGACCAGGAAGGACATTGGGGAATGGGAGCACTCGGCCGGCTTTTCAAGCTGGGAGGAGGTCCCCTTTTGTTCACACCATTTCCACGAGCTAGTGATGAAGGACGGACGCACCCTGGTGGTGCCGTGCCGAGACCAAGATGAACTCGTTGGGAGGGCGCGCATCTCACCGGGGTGCGGCTGGAGTGTCCGCGAGACGGCCTGCCTTTCAAAAGCCTACGGGCAGATGTGGCTGCTGAGCTATTTCCACCGGCGAGACCTGAGGACGCTCGGGCTCGCCATCAACTCAGCAGTGCCTGTCGATTGGGTTCCCACCGGCCGCACGACGTGGAGCATCCATGCCAGTGGGGCCTGGATGACCACAGAAGACATGCTGGACGTTTGGAACCGGGTGTGGATCCTGGACAACCCTTTCATGCAGAACAAGGAAAAGGTCATGGAGTGGAGGGATGTTCCGTACCTCCCTAAAGCTCAGGACATGTTATGTTCCTCCCTTGTTGGGAGGAGAGAAAGAGCAGAATGGGCTAAGAACATCTGGGGAGCGGTGGAAAAGGTGAGGAAGATGATAGGTCCTGAAAAGTTCAAGGACTATCTCTCCTGTATGGACCGCCATGACCTGCACTGGGAGCTCAGACTGGAGAGCTCAATAATCtaaacccagactgtgacagagcaaaacccggagggctcgcaaaagattgtccggaaccaaaacagaagcaagcaactcacagagatagagctcggactggagagctctttaaacaaaaaagccagaattgagctgaacctggagagctcattaaatatagtccagacaaaacaaaacatgacaaagtaaataggctgagctaaaagctcccaccacgggactgcttcatagcggtttgtggggggaggctaggaggcgaagccacagatcatggagtgatgcggcagcgcgcgagagcgacggggaagtggtcgcacccgacgcactatccatgaagcaatacttcgtgagacccccctggccagcaaagggggcagactggtcaggggtaagggatgcccccagagtgcattacggcagcacgccagtgagagtggcgacgggaaaatggtcgatcccgacgtagggcactctgaaaaattttgtgagaccccctgcatcatgacaaggccgaacatggtgcatgaaaaggggaggcccccggaagcacgcttccgggaggagggaagagagaaattggcagctctcttcaggatttttcctcctcctatacaaaattccccctcggtagagggggggcggttcttgttctccctgagccaccatcacccagacacagatagtctgacaaggaggtgatgtgtgactcggaaaaacacccgct |

#### Table S4. Sequence composition of flavivirus genomes and reporter proteins.

| **Group** | **Sequence (strain/species)** | **GC%** | **Length (nt)** |
| --- | --- | --- | --- |
| **Flaviviruses** | TBEV (Haselmuhl Tiho1) | 54 | 11098 |
|  | JEV (JaOArS982) | 51 | 10976 |
|  | WNV (NY99) | 51 | 11029 |
|  | KUNV (FLSDX) | 50 | 11022 |
|  | YFV (17D) | 50 | 10862 |
|  | ZIKV (H/PF/2013) | 51 | 10807 |
|  | DENV2 (16681) | 46 | 10723 |
|  | DENV4 (H241) | 47 | 10664 |
| **Fluorescent reporters** | eGFP (*Aequorea victoria*) | 62 | 717 |
|  | eYFP (*Aequorea victoria*) | 62 | 717 |
|  | Clover2 (*Aequorea victoria*) | 62 | 717 |
|  | mAmetrine (*Aequorea victoria*) | 62 | 717 |
|  | oxGFP (*Aequorea victoria*) | 62 | 717 |
|  | mCherry (*Discosoma sp.*) | 64 | 708 |
|  | turboGFP (*Pontellina plumata*) | 64 | 696 |
|  | bfloGFPa1 (*Branchiostoma floridae*) | 57 | 669 |
|  | mCardinal (*Entacmaea quadricolor*) | 58 | 732 |
|  | mNeptune2 (*Entacmaea quadricolor*) | 58 | 732 |
|  | LSSmKate2 (*Entacmaea quadricolor*) | 58 | 738 |
|  | miRFP703 (*Rhodopseudomonas palustris*) | 65 | 945 |
| **Bioluminescent reporters** | Nanoluciferase (*Oplophorus gracilirostris*) | 53 | 515 |
|  | Firefly luciferase (*Photinus pyralis*) | 45 | 1653 |
|  | Renilla luciferase (*Renilla reniformis*) | 36 | 936 |
|  | Gaussia luciferase (*Gaussia princeps*) | 59 | 555 |

### SUPPLEMENTARY FIGURES

#### Figure S1. Phylogenetic analysis of TBEV-Eu Haselmühl

**(A-D)** Maximum likelihood phylogenic trees of TBEV-Eu viruses based on (A) whole-genome sequences, (B) the coding sequence (CDS), (C) the 5′ UTR, and (D) the 3′ UTR of the indicated viruses. The trees include complete and partial genomes from TBEV-Eu strains with published recombinant viruses (Berankova et al., 2025; Haviernik et al., 2021; Hoornweg et al., 2023), and additional strains used in recent phylogenetic analyses (Bestehorn-Willmann et al., 2023; Kutschera and Wolfinger, 2022). Only viruses from the Haselmühl clade defined in A were used in C-D. Only complete genomes were used for panels C and D. For panels B–D, pairwise percent identity to Haselmühl is annotated for each segment. Branch lengths are proportional to substitutions per site; node labels show bootstrap support. **(E)** Top panel: the cHP and polyA were highlighted on the predicted 5′ and 3′ UTR RNA secondary structures of the Neudoerfl strain, as reported by Kutschera and Wolfinger (2022). Bottom panel: pairwise alignment of the Neudoerfl and Haselmühl sequences across the Neudoerfl internal polyA region and flanking nucleotides. Coordinates are shown relative to the Haselmühl sequence.

**Figure S2. TBEV Haselmühl growth curves in mammalian cell lines**

The indicated cell lines were infected with Haselmühl at the indicated MOI. RNA was harvested at the indicated time points and viral replication was quantified by qPCR. Each point represents the means and standard deviation from 2 biological replicates.

#### Figure S3. Supporting data for figures 1

**(A)** Images captured by widefield microscopy (x20) during TBEVgfp rescue, after 48h of p0 transfection (top) and after 24h of p1 infection (bottom). Images were centered on positive cells when possible. **(B)** TBEVgfp was amplified on the indicated cell lines and unless cells displayed significant CPE, supernatants were collected and titrated at the indicated times. Each dot represents one sample, with connected dots corresponding to serial measurements from the same culture.

#### Figure S4. Supporting data for figure 3

**(A)** GFP and 4G2 mean intensities (left) and infected-cell ratios (right) correlated across wells (Spearman rho = 0.78-0.87). n = 16 wells. **(B)** Mean signal intensity per infected cell after subtraction of background signal in infected and stained wells. Numbers indicate the signal-to-background ratios. Statistical significance was determined by a t-test, **** p < 0.0001, n = 16 wells. **(C)** Quantification of nuclei in infected wells live-stained with Hoechst and fixed with PFA (direct) or fixed with PFA and stain with 4G2/Alexa647 and Hoechst (after IFA), n = 16 wells. **(D)** Single-cell analysis of the experiment analyzed at well-level in Fig. 2E using the mean intensity measured per cell. Left panel: GFP and 4G2 mean intensities correlated across cells (Spearman rho = 0.81) shows no 4G2+/GFP- cells but several GFP+/4G2-. Right panel: Image analysis based on Hoechst, GFP and 4G2 signals normalized as a percentage of the siCTRL. Violin plots, truncated below 0 and above 400, represent the single-cell distribution of at least 35000 cells. Solid lines represent the median, and dotted lines indicate the 25^th^ and 75^th^ percentiles. Single-cell mean intensities were analyzed using a linear mixed-effects model with siRNA condition as a fixed effect and well identity as a random intercept. *P*-values are shown in italics. The plain line represents the median and the dotted lines represent the 25^th^ and 75^th^ percentiles.

#### Figure S5. Supporting data for figure 4

Cell viability assay of 7DMA-treated cells, n=8 biological replicates.

#### Figure S6. Phylogenetic analysis of fluorophores used in reporter orthoflaviviruses

MUSCLE alignments of fluorophore DNA sequences found on Addgene plasmids visualized with AliView highlighting majority consensus nucleotide. *Aequorea* vars share 100% coverage and 93-98% identity with eGFP. Non-*Aequorea* vars share 85% to 94% coverage and 43% to 48% identity with eGFP (lowest = Branchiostoma, highest = Entacmaea). Most abundant GFP recombination sites are indicated with their position on both TBEVgfp and eGFP sequences, and relative abundance.
