## Supplementary material for "Field-isolate recombinant tick-borne encephalitis viruses define reporter-stability guidelines for antiviral testing in flaviviruses": besson_REV_table_1.docx

**Table 1. List of recombinant flavivirus systems expressing a reporter protein.** Abbreviations: GFPvar, Aequorea victoria GFP variant; GFPtrunc, truncated GFP observed by PCR; ub, ubiquitin; fs, frameshift. Parentheses indicate optional elements; “/” indicates alternatives.

| **Virus** | **Reporter** | **Insertion site** | **Reported stability** | **Passaged in** | **Rescue** | **Reference** |
| --- | --- | --- | --- | --- | --- | --- |
| YFV | Nluc | C-(lethal)-rep-2A | Stable until p10 | Vero | Full-length clone | Baker et al., 2020 |
| ZIKV | Nluc | C-(lethal)-rep-2A | Stable until p10 | Vero | Full-length clone | Baker et al., 2020 |
| TBEV | Turbo-GFP | C-2A-rep-2A | Excised from p7 | BHK-21 | ISA | Berankova et al., 2025 |
| YFV | GFPvar (eGFP) | E-NS1 | Stable until p10, GFPtrunc detected from p2 | Vero | Full-length clone | Bonaldo et al., 2007 |
| DENV2 | GFPvar (OxGFP), mCherry, Nluc | C-rep-2A | Excised from p1 to p4 | Vero | Full-length clone | Cherkashchenko et al., 2023 |
| DENV4 | GFPvar (OxGFP), mCherry, Nluc | C-rep-2A | Excised from p1 to p4 | Vero | Full-length clone | Cherkashchenko et al., 2023 |
| KUNV | GFPvar (OxGFP), mCherry, Nluc | C-rep-2A | Excised from p1 to p4 | Vero | Full-length clone | Cherkashchenko et al., 2023 |
| ZIKV | GFPvar (OxGFP), mCherry, Nluc | C-rep-2A | Excised from p3 to p4 | Vero | Full-length clone | Cherkashchenko et al., 2023 |
| POWV | GFPvar (split eGFP = res. 1-214) | NS1-Cter | NA | NA | CPER | Conde et al., 2023 |
| CxFV | GFPvar (GFP) | C-rep-2A | Excised from p4 | C6/36 | CPER | Dong et al., 2023 |
| YFV | Nluc | C-rep-2A | Excised from p3 to p4 | BHK-21 | Full-length clone | Dong et al., 2021 |
| DENV4 | Nluc | C-rep-2A | NA | NA | Full-length clone | Fang et al., 2022 |
| ZIKV | GFPvar (eGFP) | C-rep-2A | Excised from p1 to p4 | Vero | Full-length clone | Gadea et al., 2016 |
| ZIKV | GFPvar (eGFP) | C-rep-2A | Stable until p5 (data not shown) | NA | Full-length clone | Gao et al., 2021 |
| LGTV | GFPvar (eGFP) | E-NS1 | NA | NA | Full-length clone | Grabowski et al., 2017 |
| TBEV | mCherry | C-2A-rep-2A | Excised from p1 to p4 | BHK-21 | ISA | Haviernik et al., 2021 |
| DTMUV | Nluc | C-rep-2A | Stable until p10 | BHK-21 | Full-length clone | He et al., 2019 |
| WNV | mCherry | C-2A-rep-2A | Stable until p5 | Vero | Full-length clone | Kobayashi et al., 2023 |
| JEV | GFPvar (eGFP) | C-rep-2A | Stable until p10 | BHK-21 | Full-length clone | Li et al., 2022 |
| DENV2 | mCherry | C-rep-2A | Excised from p4 to p6 | Vero | Full-length clone | Li et al., 2020 |
| JEV | Rluc | C-rep-2A | Excised from p2 to p5 | BHK-21 | Full-length clone | Li et al., 2017 |
| ZIKV | GFPvar (eGFP), mCherry, Nluc | C-rep-(ub)-2A | Excised from p2-p4 (eGFP, mCherry), stable until p4 (Nluc) | Vero | Full-length clone | Mutso et al., 2017 |
| DENV4 | GFPvar (eGFP) | C-2A-rep-2A | Stable until p3 | Vero | ISA | Park et al., 2023 |
| WNV | GFPvar (GFP) | 3'UTR | Excised from p2 to p4 | HEK293T | Full-length clone | Pierson et al., 2005 |
| YFV | Nluc | C-rep-2A-ub | Stable until p3 | BHK-21 | Full-length clone | Ramanathan et al., 2020 |
| DENV2 | GFPvar (GFP) | C-rep-2A | Excised from p1 to p4 | Vero | Full-length clone | Schoggins et al., 2012 |
| DENV2 | GFPvar (eGFP, Clover2), bfloGFP | C-(2A)-rep-2A | Excised from p1-2 to p4 | Vero | Full-length clone | Suphatrakul et al., 2018 |
| DENV2 | GFPvar (mAmetrine), mCherry, miRFP703, LSSmKate2, mCardinal, mNeptune2 | C-(2A)-rep-2A | NA | NA | Full-length clone | Suphatrakul et al., 2018 |
| ZIKV | GFPvar (GFP), Nluc | C-(fs2A)rep-2A/ub,  E-NS1 | Stable until p10,  GFPtrunc detected | Vero | Full-length clone | Volkova et al., 2020 |
| YFV | GFPvar = Venus | C-rep-2A | NA | NA | Full-length clone | Yi et al., 2011 |
| JEV | GFPvar (GFP) | 3'UTR | NA | BHK-21 | Full-length clone | Yun et al., 2003 |
| ZIKV | GFPvar (GFP), Nluc | 3'UTR | Excised at p5 (data not shown) | BHK-21 | Full-length clone | Yun et al., 2020 |
| JEV | GFPvar (eGFP) | 3'UTR | Excised from p5 | C6/36 | Full-length clone | Zhang et al., 2020 |
