## Supplementary figures and images for "Field-isolate recombinant tick-borne encephalitis viruses define reporter-stability guidelines for antiviral testing in flaviviruses"

### FIG_S1.tiff

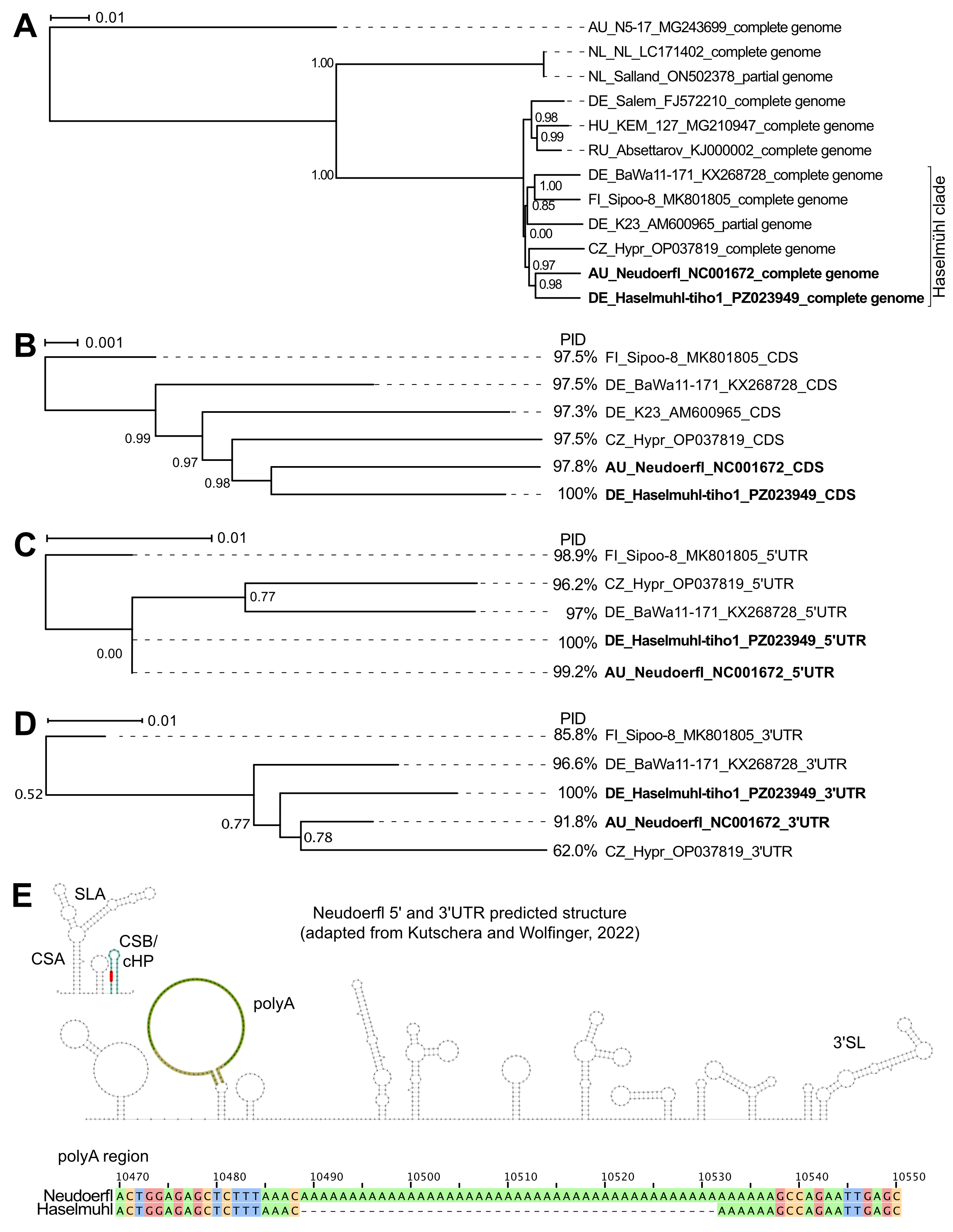

### FIG_S2.tiff

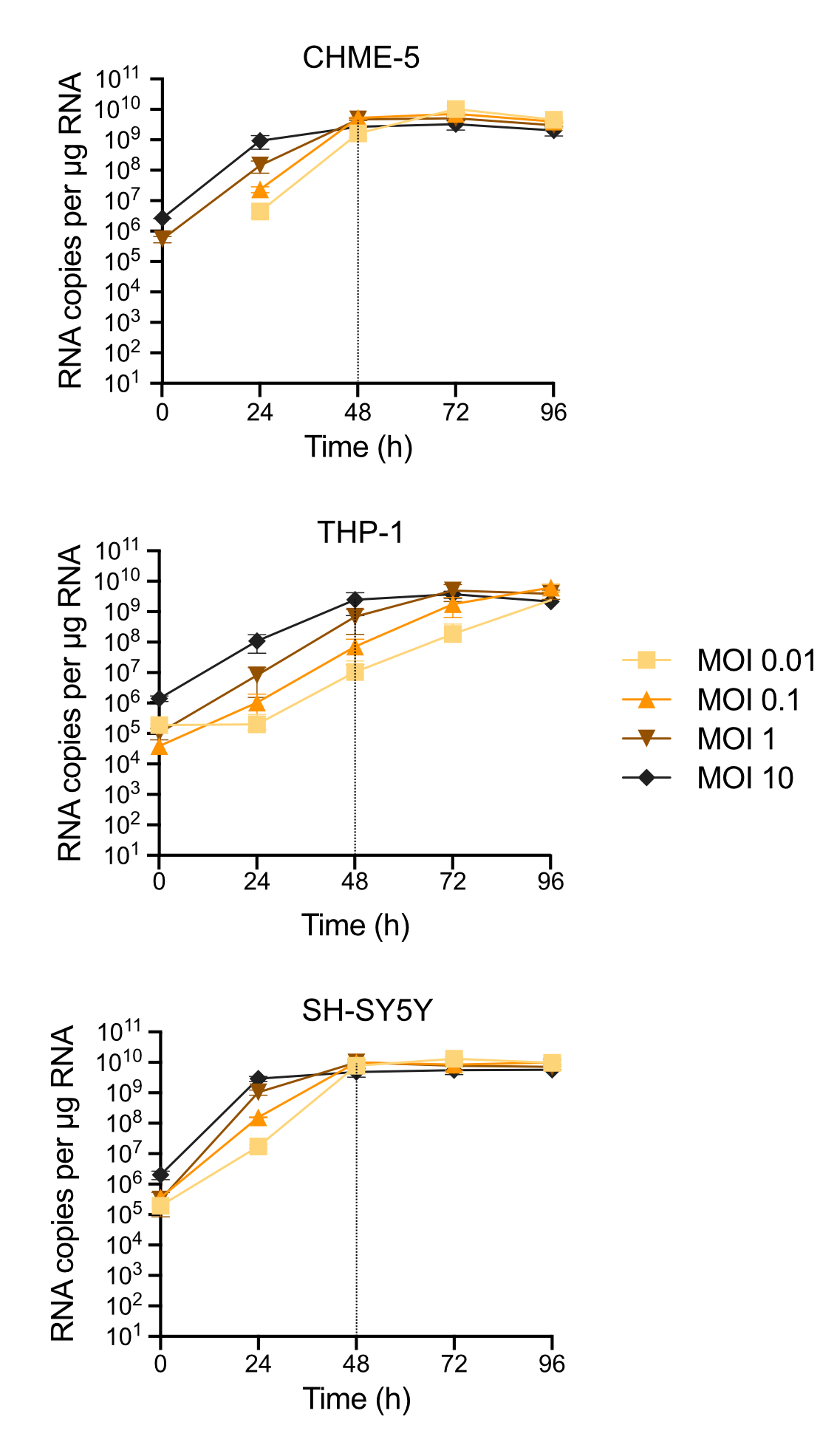

### FIG_S3.tiff

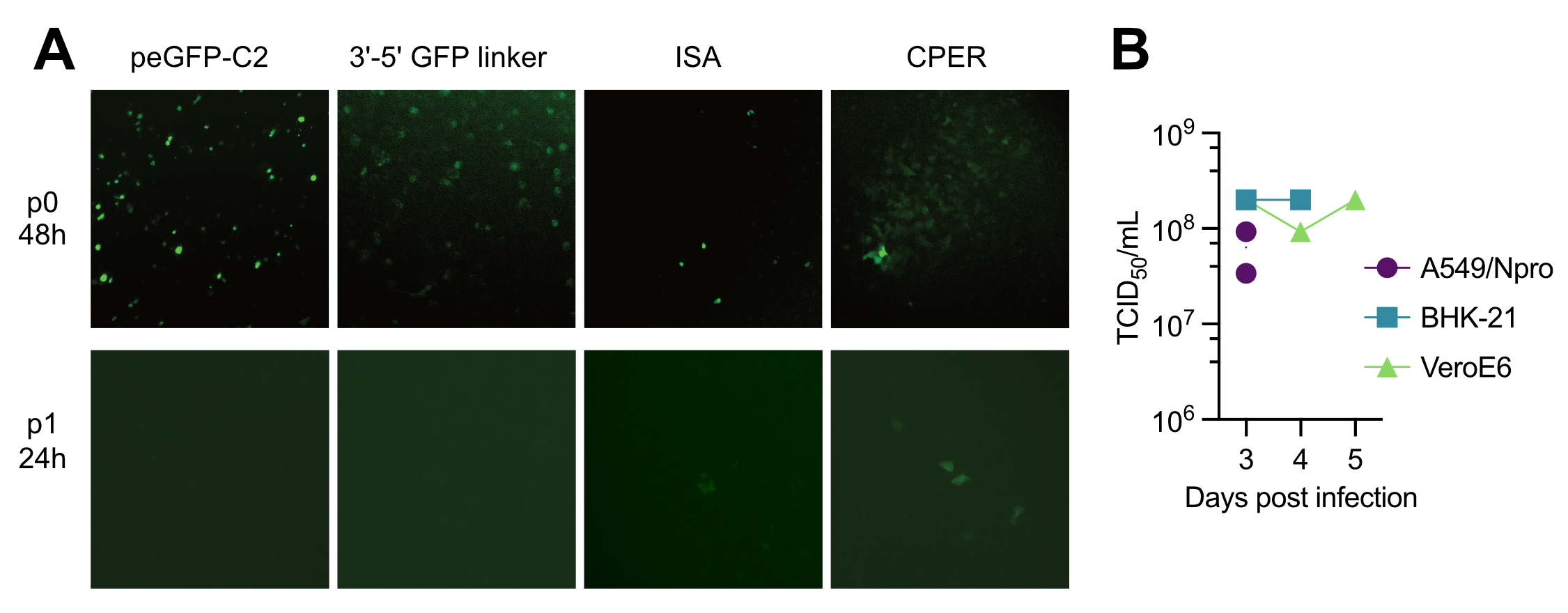

### FIG_S4.tiff

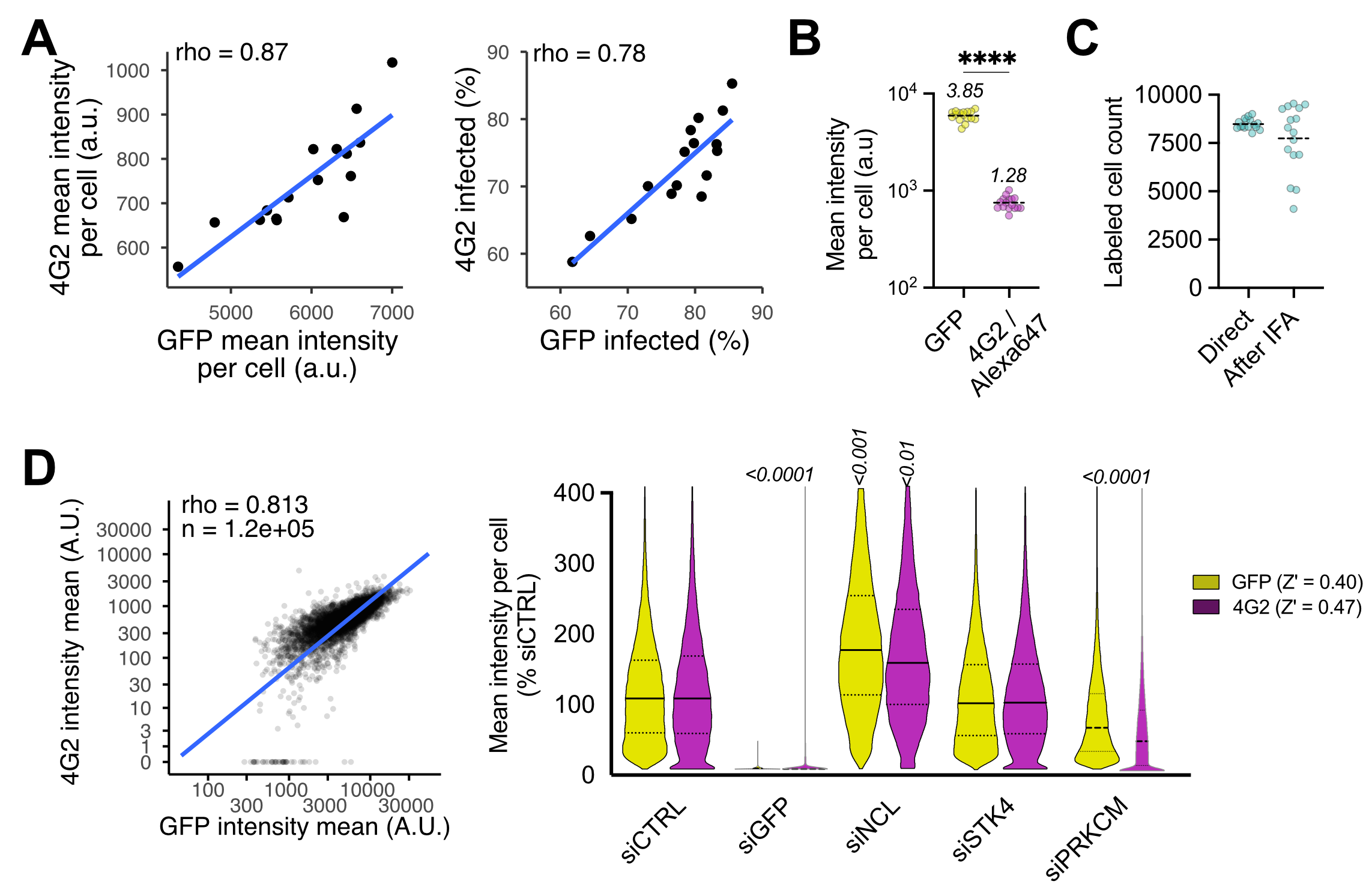

### FIG_S5.tiff

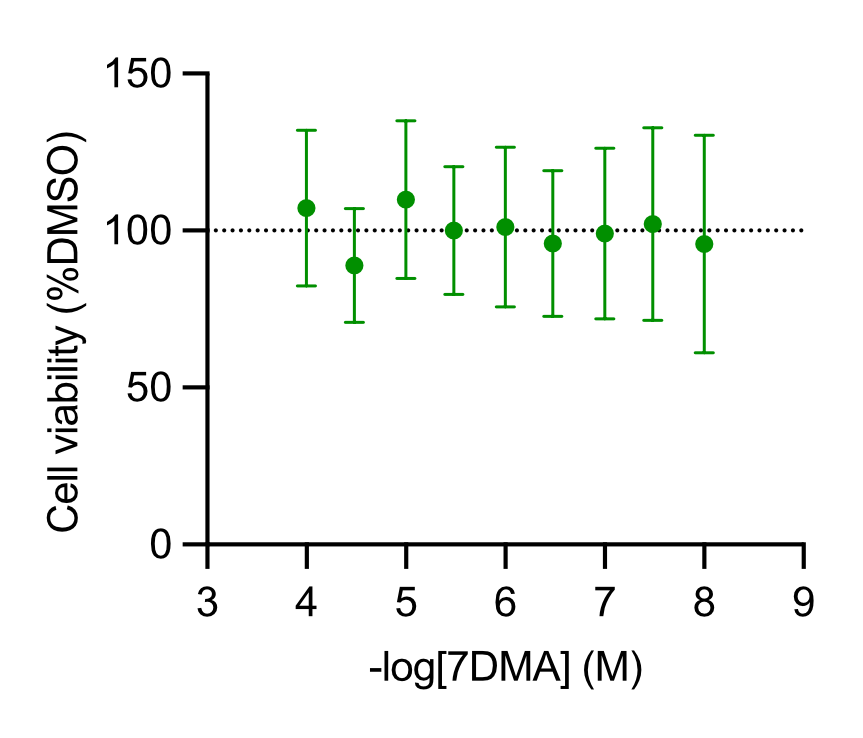

### FIG_S6.tiff

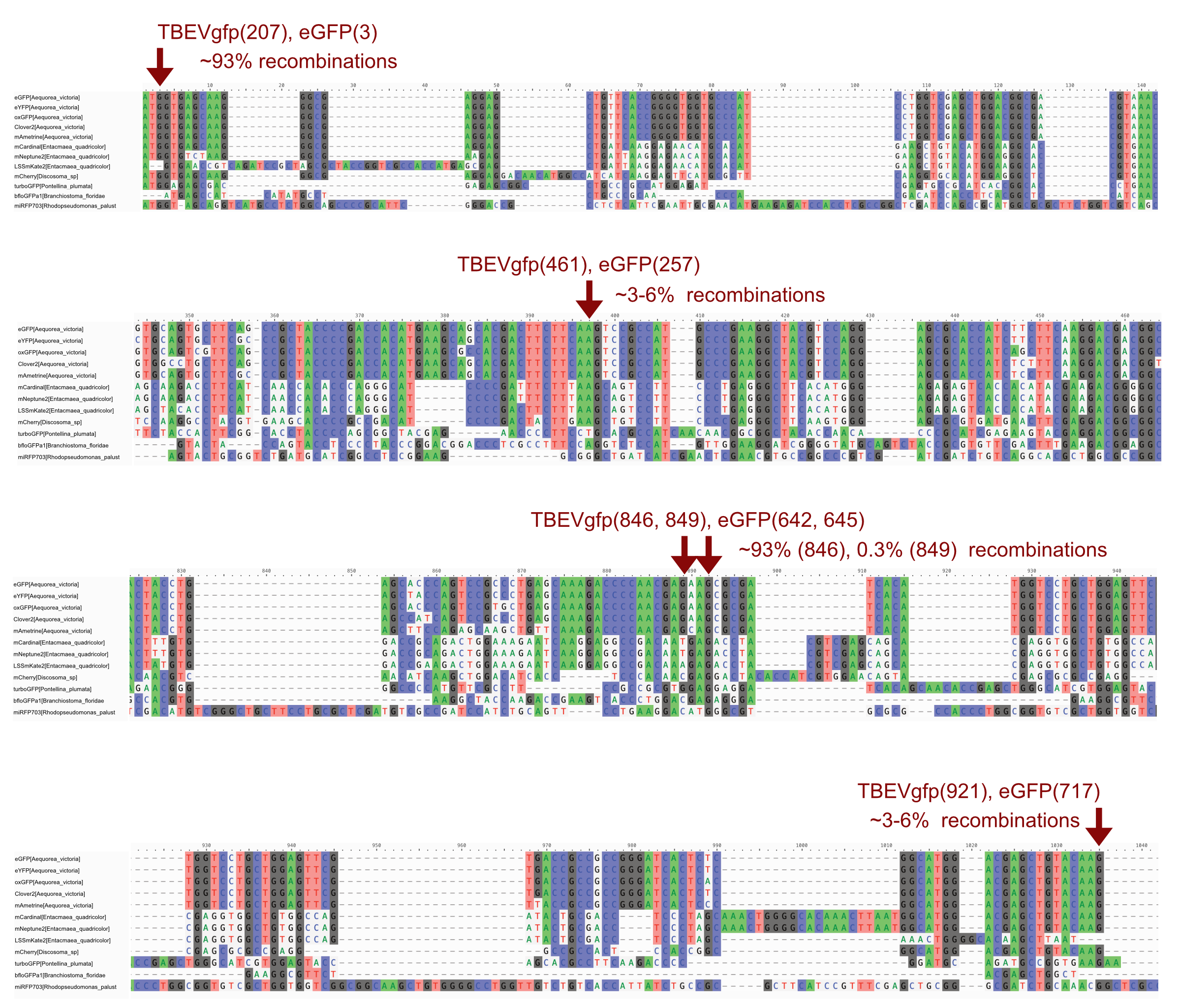
